# Pseudomonas aeruginosa uses kinases NahK and RetS to control the motile-biofilm switch

**DOI:** 10.1101/2025.09.24.678285

**Authors:** Jason M. Withorn, Karishma Ramcharan, Natalie E. Alfano, Alicia G. Mendoza, Jiayuan Fu, Danielle Guercio, Aine MacDermott, Kendra E. Byrne, Elizabeth M. Boon

**Author notes:** **Materials and Correspondence** Correspondence and requests for materials should be addressed to Elizabeth M. Boon.

## Abstract

The multidrug-resistant bacterium *Pseudomonas aeruginosa* (*Pa)* poses a significant threat to public health. This Gram-negative bacterium establishes pathogenicity through formation of multicellular communities, known as biofilms, that result in significant resistance to antibiotics and host immune systems. In *Pa*, the motile-to-biofilm transition is regulated through an interconnected signaling network known as the Gac Multikinase Network (Gac-MKN). This network comprises two regulatory branches: the HptB signaling network and GacS/A signaling network. In the Gac-MKN, several histidine kinases converge to regulate the activity of the post-transcriptional regulator protein, RsmA. Although previous studies have assessed the role of individual kinases in this network, the role of each Gac-MKN kinase in regulating RsmA activity has not been quantitatively characterized and compared in side-by-side experiments in the same reference strain, which is presented here. In this study, we show that kinases NahK and RetS are the predominant regulators of the Gac-MKN. Through controlled testing of RsmA-dependent phenotypes, we demonstrate loss of *nahK* or *retS* leads to complete inactivation of RsmA, triggering biofilm formation. Our results support previous findings that RetS regulates RsmA through the GacS/A network but present the new finding that NahK is the central kinase involved in HptB phosphorylation; previous studies have attributed HptB phosphorylation to PA1611 and SagS. Our findings demonstrate that NahK signaling controls RsmA activity to rapidly transition between the motile and biofilm states. We anticipate the results of this study will facilitate the use of targeting the Gac-MKN to trigger biofilm dispersal for improved antibiotic treatment.

## Introduction

Bacteria have evolved over time to grow and reside in various hostile environments. As an individual bacterium, the cell is more likely to succumb to antibiotics, host immune responses, and unfavorable environmental conditions. Bacteria form multicellular communities called biofilms to circumvent these conditions. Bacteria encapsulated in a biofilm are not only protected from harsh extracellular conditions, but are also 10-to 1000-fold more resistant to antibiotics.^1^ This transition from individual, motile bacteria to the biofilm, sessile state is tightly regulated by many intracellular signaling networks that respond to environmental stimuli and bacterial signaling molecules.^2^

*Pseudomonas aeruginosa (Pa)* is a Gram-negative, opportunistic pathogen that utilizes these multicellular biofilms to establish a wide range of host infections, such as lung infections in cystic fibrosis patients, ventilator-associated pneumonia, and both topical and catheter-based infections.^3,4^ Over the last 30 years, a multitude of studies have probed various signaling networks within the *Pa* genome and have identified an interconnected signaling system that the bacterium primarily utilizes to regulate the transition between the motile and biofilm states, the Gac Multikinase Network (Gac-MKN).^5-7^

The Gac-MKN has two signaling branches: the HptB signaling network and the GacS/A signaling network (**Figure 1A**).^8-10^ In the hypervirulent *Pa* strain UCBPP-PA14 (PA14), the Gac-MKN encompasses five sensory hybrid-histidine kinases: GacS (PA14_52260), RetS (PA14_64230), PA1611 (PA14_43670; hereby referred to as PA1611), NahK (PA14_38970; previously referred to as ErcS’), and SagS (PA14_27550).^8-10^ Although the environmental signals and intracellular cues that regulate the activity of these kinases remain mostly uncharacterized, several signals have been shown to directly regulate the kinases in this network, such as glucose-6-phosphate, kin cell lysis, mucin-glycans, and nitric oxide (NO). ^11-14^

**Figure 1.**
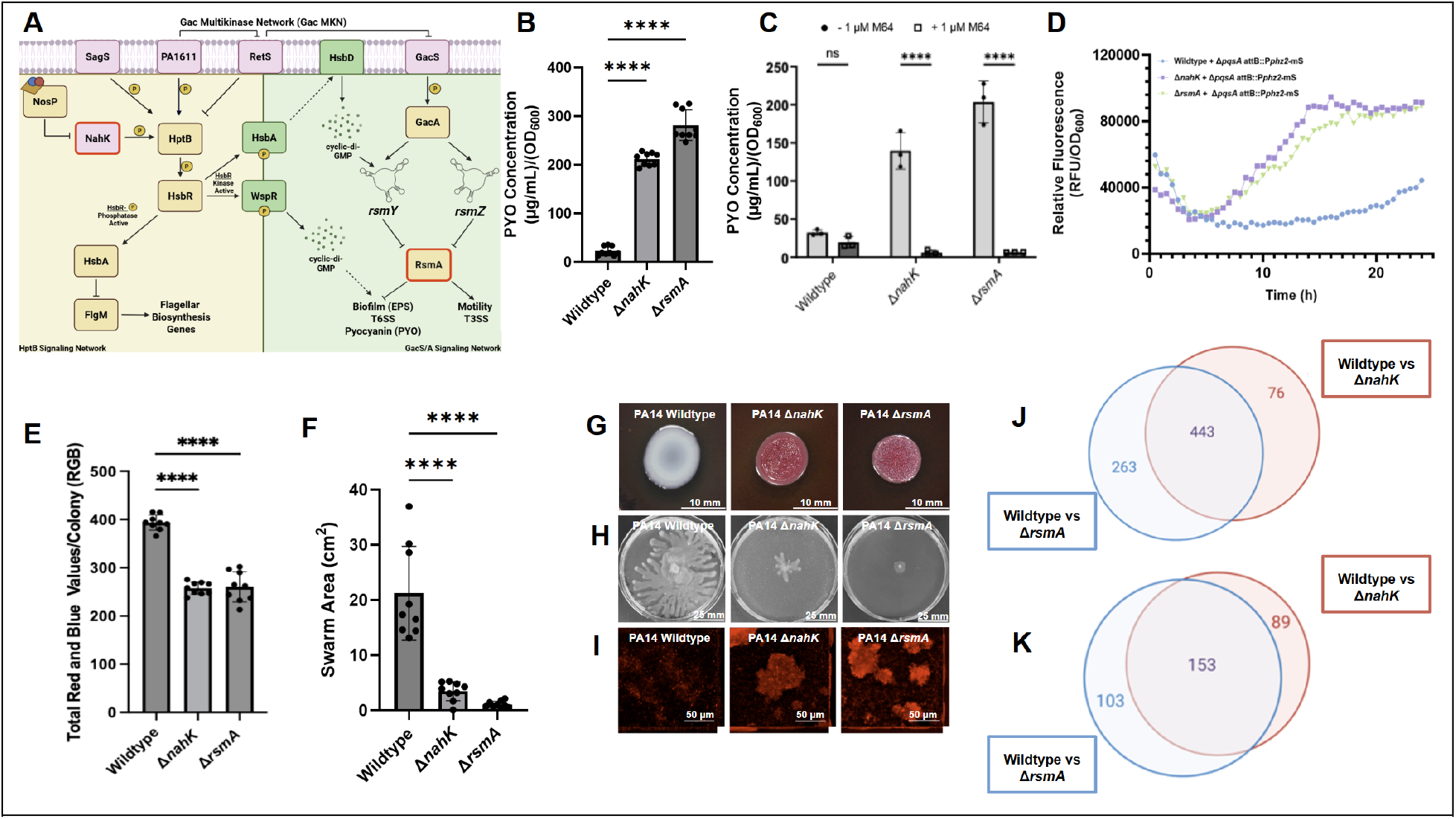
Loss of *nahK* signaling leads to inactivation of RsmA. **(A)** Schematic of Gac-MKN. HptB branch (yellow); GacS/A branch (green). NahK, PA1611, RetS, and SagS all modulate the phosphorylation state of HptB. Without HptB phosphorylation, intracellular cyclic-di-GMP increases and the transcription of *rsmY* is upregulated. GacS phosphorylates transcription factor GacA to activate transcription of both small regulatory RNAs *rsmY* and *rsmZ*. These ncRNAs inactivate RsmA to trigger biofilm formation. **(B)** PYO [(μL/mL)/OD_600_)] of PA14 strains from planktonic, liquid LB cultures (n=9), and **(C)** with/without supplementation of 1 µM M64 PqsR inhibitor (n=3).^29^ (520 nm; ε = 17.072^53^). **(D)** RFU/OD_600_ of mScarlett reporter for *phz2* phenazine biosynthesis operon genomically integrated in Δ*pqsABC* reporter strain cocultured with planktonic PA14 strains (n=3). **(E)** RGB quantification for EPS staining of PA14 colony biofilms cultured on 1% tryptone, 1% BactoAgar media supplemented with 40 µg/mL Congo-Red dye and 20 µg/mL Brilliant Blue dye. (n = 9). **(F)** Swarming area (cm^2^) of PA14 strains cultured on low-density agar media. (n=9) **(G)** Representative images of PA14 static Congo-Red colony biofilms. Increased red colony color correlates to the quantity of EPS secreted by each strain (n = 9). **(H)** Representative images of PA14 swarming motility on low-density agar (n=9). Tendril formation correlates to increased flagellar-mediated motility. **(I)** Representative CLSM images of PA14 strains grown under continuous flow conditions. Cells visualized with nucleic acid stain SYTO62, images displayed as compiled z-stack (35 slices; 0.5 µm slices) flanked by the XY and YZ side profiles. Visible biofilm macrostructures observed for Δ*nahK* and Δ*rsmA* (n = 9). **(J)** Venn diagram of upregulated genes detected in Δ*nahK* and Δ*rsmA* relative to wildtype. 443 genes commonly upregulated in each deletion strain. **(K)** Venn diagram of downregulated genes in Δ*nahK* and Δ*rsmA* relative to wildtype. 153 genes are commonly downregulated in each deletion strain. Error bars represent standard deviation from the mean. Significant differences are indicated with asterisks (N.S. *P* > 0.05; \**P* < 0.05; \*\**P* < 0.01; \*\*\**P* < 0.001; \*\*\*\**P* < 0.0001; one-way ANOVA and a Tukey multiple comparisons test).

NO is a well-documented signaling molecule capable of triggering biofilm dispersal across numerous species of biofilm-forming bacteria.^15-17^ Our lab has been interested in characterizing how NO mediates the biofilm-to-motile transition across pathogenic bacteria.^18^ In *Pa*, NO has been shown to inhibit autophosphorylation of NahK, preventing phosphotransfer to the histidine phosphotransfer protein, HptB.^14^ Additional kinases in the Gac-MKN have also been shown to modulate the phosphorylation state of HptB, such as PA1611, SagS, and RetS (**Figure 1A**).^9,19-20^ Although RetS has a degenerative histidine kinase domain, it has been shown to function as a phosphatase for HptB and to inhibit PA1611 through heterodimerization.^9,21^ It has been reported that collectively, NahK, PA1611, RetS, and SagS control the HptB branch of the Gac-MKN by regulating HptB phosphorylation. HptB phosphorylation promotes flagellar biosynthesis through downstream sequestration of the flagellar anti-sigma factor FlgM, while de-phosphorylation of HptB leads to increased transcription of the small regulatory RNA *rsmY*, through increasing levels of intracellular cyclic-di-GMP, to promote biofilm formation.^22-24^

GacS is the major kinase involved in the GacS/A branch of the Gac-MKN. GacS phosphorylates the response regulator GacA to induce transcription of both small regulatory RNAs *rsmY* and *rsmZ*, which function to inactivate the post-transcriptional regulatory protein, RsmA.^5^ RsmA controls the translation of over 500 genes in *Pa*, including those responsible for quorum-sensing (QS), antibiotic resistance, virulence, and proliferation (**Figure 1A**).^25,26^ RetS also participates in the GacS/A branch to inhibit GacS phosphotransfer through heterodimerization.^21^ GacS-RetS interaction prevents phosphorylation of GacA, keeping RsmA in the active state.

Previous work by our group has shown that deletion of *nahK* in PA14 increases pyocyanin (PYO) production through increased biosynthesis of the *Pseudomonas* quinolone signal (PQS) and cell elongation in the presence of NO under anaerobic conditions.^27,28^ In this study, we determine Δ*nahK* phenotypes, such as hyper-biofilm formation, decreased motility, and PYO overproduction result from inactivation of RsmA. We find that NahK has predominant control over HptB phosphorylation, as opposed to SagS and/or PA1611, and that NahK-to-HptB phosphorylation has significant control over RsmA activity. Indeed, the transcriptional regulon and biological phenotypes of Δ*nahK* and Δ*rsmA* strains largely overlap. NahK-to-HptB phosphorylation modulates expression of *rsmY* and *rsmZ* through regulation of intracellular cyclic-di-GMP; however, we report this regulation is not solely controlled by HsbR/A/D and WspR systems as previously reported. ^23,24^ Finally, we provide further insight into the role of PA1611 and RetS in the GacS-MKN through kinase overexpression; the main function of PA1611 is to regulate the RetS-to-GacS heterodimerization. Together, this study outlines the basal signaling cascade of the Gac-MKN and provides insight into how RsmA activity varies due to Gac-MKN kinase fluctuations.

## Results

### Deletion of nahK leads to inactivation of RsmA

In a previous study, we found that Δ*nahK* overproduces PYO by activating the PQS QS system, which upregulates transcription of the *phz2* PYO biosynthetic pathway.^27^ Further, we demonstrated that overexpression of *rsmA* in Δ*nahK* complements this PYO overproduction phenotype.^27^ Presumably, overexpression of *rsmA* results in an excess of uninhibited RsmA, suggesting the main driver of Δ*nahK* phenotypes is reduced RsmA activity. As a result, we wondered if Δ*nahK* and Δ*rsmA* might have similar phenotypes. Thus, we obtained and characterized a Δ*rsmA* strain with respect to its PYO production phenotypes. Indeed, our data indicates that both Δ*nahK* and Δ*rsmA* increase PYO production by the same order of magnitude when compared to the PA14 wildtype strain (**Figure 1B**). We also found that in the presence of M64,^29^ an inhibitor of the PQS regulator PqsR, neither Δ*nahK* nor Δ*rsmA* overproduce PYO (**Figure 1C**). Further, we cocultured wildtype, Δ*nahK*, and Δ*rsmA* strains with a *Pa* mutant that is no longer able to produce PQS, but encodes a fluorescent reporter of the *phz2* operon (Δ*pqsA* attB::P*phz2-*mScarlet), and found that both Δ*nahK* and Δ*rsmA* produce sufficient PQS to activate the fluorescence in the reporter strain (**Figure 1D**), confirming these strains overproduce PQS. Overall, these data suggest that loss of NahK signaling results in inactivation of RsmA, leading to increased PYO production through a PQS-dependent mechanism.

In addition to PYO production, RsmA regulates the translation of systems that contribute to the planktonic-to-biofilm switch, such as bacterial motility and secretion of extracellular polymeric substance (EPS).^30,31^ Therefore, Δ*nahK* and Δ*rsmA* were then screened using three phenotypic assays controlled by these systems: swarming motility, Congo-Red staining to view EPS production, and continuous flow biofilm culturing for macrostructure visualization (**Figure 1 E-I**). Swarming is a complex, QS-and flagellar-mediated bacterial movement across low-density nutrient agar.^32^ Δ*rsmA*, as previously reported,^30^ results in an abolishment of swarming motility and Δ*nahK* results in substantially reduced swarming (**Figure 1F,1H**, **Supplementary Figures S1B-S1C**). Biofilm-forming bacteria secrete EPS to promote biofilm formation by facilitating cell-to-surface adhesion and cell-to-cell cohesion.^33^ Congo-Red dye can be used to visualize the production of EPS as the anionic dye interacts with the secreted polysaccharides in the EPS, resulting in red coloring of cultured colonies.^34^ Both Δ*nahK* and Δ*rsmA* show increased Congo-Red dye uptake relative to the wildtype (**Figure 1E,1G**). Loss of bacterial motility and increased EPS production suggest loss of either *nahK* or *rsmA* results in increased biofilm formation. Culturing biofilm-forming bacteria under continuous-flow, coupled with confocal-laser scanning microscopy (CLSM), allows for non-invasive imaging of mature biofilms under dynamic conditions.^35,36^ After 72 hours of culturing under continuous flow, visible macrostructures were observed in both Δ*nahK* and Δ*rsmA*, while no macrostructures were detected in the wildtype strain (**Figure 1I**). The lack of macrostructure in PA14 wildtype is expected due to degradation of the *psl* exopolysaccharide biosynthesis operon.^37-39^ Overall, Δ*nahK* behaves phenotypically identically to Δ*rsmA* in our assays.

RNA-sequencing was used to further compare NahK and RsmA by determining the extent of transcriptional overlap between the Δ*nahK* and Δ*rsmA* strains, relative to the wildtype strain (**Figure 1J,1K**, **Supplementary Figure S1G**,**S1H, Supplementary Tables S4-5**). Transcriptome analysis shows that out of all genes upregulated in the Δ*nahK* strain relative to the wildtype, approximately 63% are also upregulated in the Δ*rsmA* strain relative to wildtype (**Figure 1J**). Similarly, out of all genes downregulated in the Δ*nahK* strain relative to wildtype, approximately 60% are also downregulated in the Δ*rsmA* strain relative to wildtype (**Figure 1K**). Indeed, when directly comparing Δ*nahK* to Δ*rsmA*, only a few genes are up-or downregulated to significantly different extents, underscoring the similarity in transcriptional profiles between these strains (**Table S4**). In agreement with the phenotypes examined in **Figure 1B-I**, many transcripts dysregulated in both genomic deletion strains with respect to the wildtype strain are involved in PYO production (*phz1* and *phz2* phenazine biosynthesis), QS (*qslA/pqs* biosynthesis), motility (flagellar biosynthesis, type IV pili assembly, and *cupD* fimbae), and biofilm formation (*pel* polysaccharide) (**Supplementary Tables S4-5**). Transcripts for additional systems previously linked to RsmA-regulation were also detected in both Δ*nahK* and Δ*rsmA*, such as transcripts involved in lipopolysaccharide (LPS) biosynthesis, O-specific antigen (OSA) biosynthesis, and protein catabolic processes.^25^ Interestingly, we identified transcripts of novel systems highly dysregulated in both Δ*nahK* and Δ*rsmA*, such as the production of r-bodies and mercuric resistance.^40,41^ Overall, Δ*nahK* has a similar transcriptional regulon as Δ*rsmA*, suggesting that both gene products function to regulate the same phenotypes.

In addition to determining how the regulon of Δ*nahK* is different than Δ*rsmA*, we also compared a double deletion strain of co-cistronic genes *nosP*, the NO sensor in *Pa*, and *nahK* (Δ*nosP*Δ*nahK*) to the wildtype (**Supplementary Table S5**). While the analysis of the Δ*nosP*Δ*nahK* double deletion relative to wildtype was done under different growth media than the Δ*nahK* deletion alone, we observed similar phenotypes and transcriptional differences between these two strains, suggesting that NahK exerts the primary functions of this operon.

However, there are differences between Δ*nahK* and Δ*rsmA*, specifically regarding the regulation of the secretion systems. It has been shown that RsmA regulates the Type 3 Secretion System (T3SS) and Type 6 Secretion System (T6SS) inversely depending on if RsmA is active or inhibited, respectively.^42-43^ However, Δ*nahK* does not have this inverse effect on the T3SS and T6SS. Upregulation of transcripts specific to T3SS effectors (*exoT* and *exoY*) and the T3SS translocation apparatus (*popBD, pcrV*) were identified in Δ*nahK* relative to wildtype, but were downregulated in Δ*rsmA* relative to wildtype. Further research could focus on how NahK and RsmA differentially regulate these systems, as this points to a divergence of the two signaling pathways.

Collectively, these data suggest that NahK signaling plays a major role in modulating the activity of RsmA to regulate multiple cellular processes in *Pa*. It is well established that NahK is a kinase that contributes to RsmA regulation through the HptB branch of the Gac MKN (**Figure 1A**).^8-10,14,18,22-24,27-28^ The simplest explanation of these data are consistent with NahK being the major regulator of HptB, as opposed to PA1611, RetS, or SagS. Thus, inhibition of NahK autophosphorylation leads to downstream inhibition of RsmA, promoting biofilm formation.

It remains unclear how NahK, or other members of the GacS MKN, control the post-transcriptional regulation of RsmA, as Δ*rsmA* has many additional targets outside of the transcriptional regulon observed in Δ*nahK*. It is of interest to us that Δ*nahK* appears to have additional targets outside of RsmA control. Future research will focus on understanding how this subset of genes differs from those influenced by RsmA. One potential mechanism for these differences may be through signaling downstream of HptB, as it is currently unknown how HptB signaling influences systems outside of modulating of intracellular cyclic-di-GMP and flagellar biosynthesis.^9,22-24,40,44^

### NahK, HptB, and RetS function to keep RsmA active

We then aimed to determine how other signal transduction proteins within the Gac-MKN play a role in modulating the activity of RsmA. To do this, we compared individual genomic deletion mutants of every gene in the Gac-MKN for PYO/PQS production, biofilm formation, and swarming motility (**Figure 2A-2H**). When comparing PYO production for each genomic deletion mutant in the Gac-MKN, we found that deletions of *nahK, hptB, retS*, and *rsmA* all have hyper-PYO phenotypes; these mutants each overproduce PYO by the same order of magnitude (**Figure 2A**). Interestingly, Δ*rsmY* and Δ*rsmZ*, strains deleting the small regulatory RNAs that directly inhibit RsmA, produce intermediate levels of PYO, relative to levels produced by wildtype and any of the hyper-PYO mutants. However, a double deletion of *rsmY* and *rsmZ* produces wildtype levels of PYO. Quantitative-PCR for *rsmY* and *rsmZ* in each single deletion strain was done to determine if loss of one RNA affected the transcription of the other (**Supplementary Figure S2**). Loss of *rsmY* resulted in upregulation of *rsmZ*, however, loss of *rsmZ* did not affect the transcription of *rsmY*. Further research could focus on the interconnected regulation of these small RNAs, as this points towards unknown regulatory feedback mechanisms for each RNA. We then determined that the hyper-PYO phenotype observed in Δ*hptB*, Δ*retS* and Δ*rsmA* was through the same PQS-*phz2*-mediated production as Δ*nahK*. When cultured with the M64 PQS inhibitor, a complete reduction of PYO production was observed in each of the hyper-PYO strains, suggesting PQS mediates phenazine production in each strain (**Figure 2B**).^29^ We then confirmed the PQS secreted by these strains activated a *phz2* mScarlet reporter in a co-culture assay as previously described (**Figure 2C**).^27^ Together, this data suggests that Δ*nahK*, Δ*hptB*, Δ*retS*, and Δ*rsmA* dysregulate PQS QS systems leading to an increase in PYO production.

**Figure 2:**
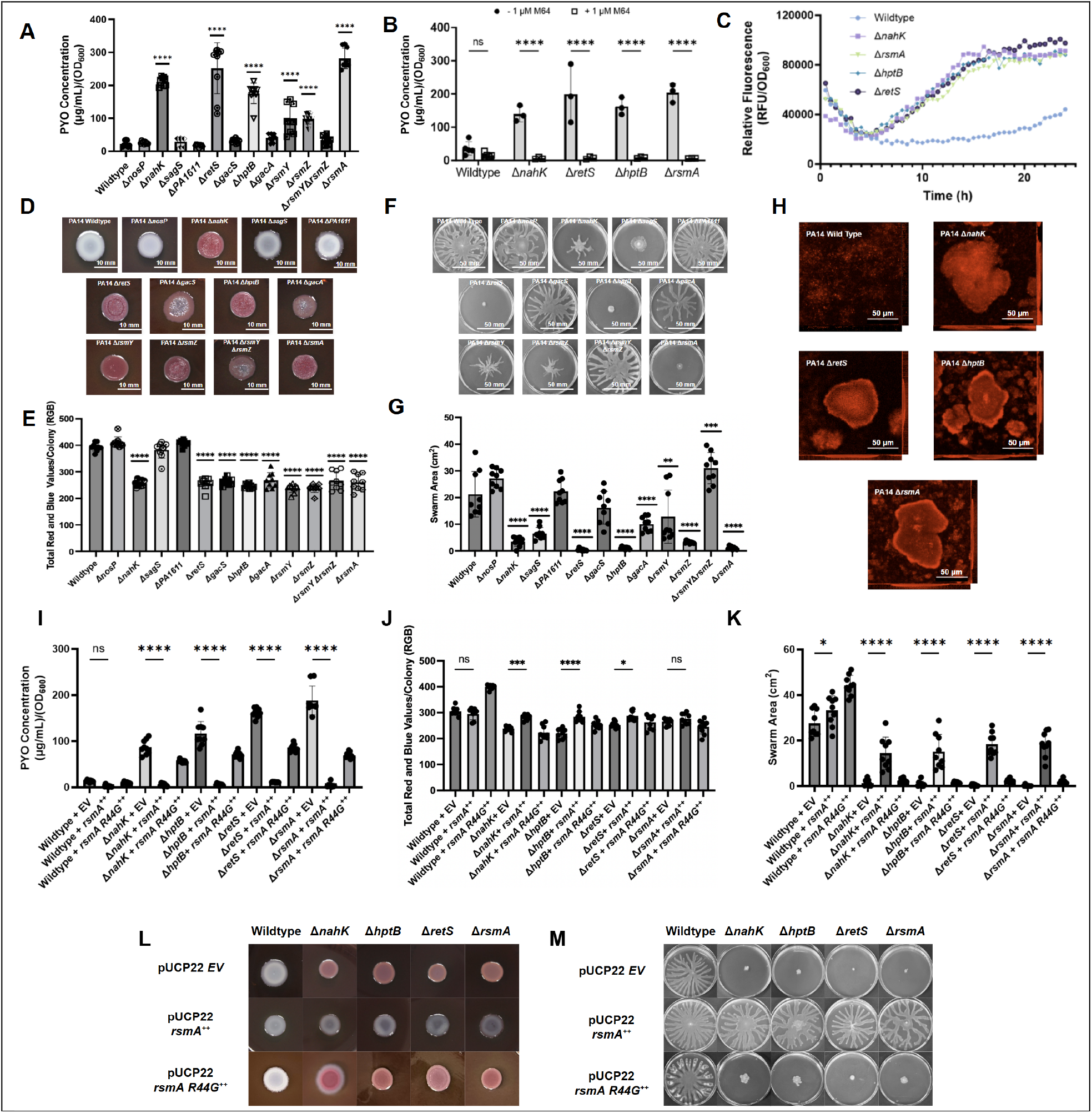
Deletion of *nahK, hptB* and r*etS* phenocopy RsmA inactivation. **(A)** PYO [(μL/mL)/OD_600_)] of PA14 strains from planktonic, liquid LB cultures (n=9), and **(B)** with/without supplementation of 1 µM M64 PqsR inhibitor (n=3). **(C)** RFU/OD_600_ of mScarlett reporter for *phz2* phenazine biosynthesis operon genomically integrated in Δ*pqsABC* reporter strain cocultured with planktonic PA14 strains (n=3). **(D)** Representative images of PA14 static Congo-Red colony biofilms. (n = 9). **(E)** RGB quantification for EPS staining of PA14 colony biofilms cultured on 1% tryptone, 1% BactoAgar media supplemented with 40 µg/mL Congo-Red dye and 20 µg/mL Brilliant Blue dye. (n = 9). **(F)** Representative images of PA14 swarming motility on low-density agar (n=9). **(G)** Swarming area (cm^2^) of PA14 strains cultured on low-density agar media. (n=9) **(H)** Representative images of swarming motility (n = 9). **(I)** Representative CLSM images of PA14 strains grown under continuous flow conditions. Cells visualized with SYTO62, images displayed as compiled z-stack (35 slices; 0.5 µm slices) flanked by the XY and YZ side profiles. Visible biofilm macrostructures observed for Δ*nahK*, Δ*hptB*, Δ*retS*, and Δ*rsmA* (n = 9). **(I-M)** Phenotypic characterization for RsmA-mediated phenotypes of PA14 strains containing pUCP22 overexpression vector containing either functional wildtype RsmA or nonfunctional RsmA R44G. **(I)** PYO [(μL/mL)/OD_600_)] of PA14 strains from planktonic, liquid LB cultures (n=9) **(J)** RGB quantification for EPS staining of PA14 colony biofilms cultured on 1% tryptone, 1% BactoAgar media supplemented with 40 µg/mL Congo-Red dye and 20 µg/mL Brilliant Blue dye. (n = 9). **(K)** Swarming area (cm^2^) of PA14 strains cultured on low-density agar media. (n=9). **(L)** Representative images of *P. aeruginosa* static Congo-Red colony biofilms. (n = 9). **(M)** Representative images of PA14 swarming motility on low-density agar (n=9). Error bars represent standard deviation from the mean. Significant differences are indicated with asterisks (N.S. *P* > 0.05; \**P* < 0.05; \*\**P* < 0.01; \*\*\**P* < 0.001; \*\*\*\**P* < 0.0001; one-way ANOVA and a Tukey multiple comparisons test).

Next, we explored how each gene in the Gac-MKN contributes to EPS production (**Figure 2D, 2E**), swarming motility (**Figure 2F,2G**), and biofilm macrostructure formation (**Figure 2H**), i.e., other phenotypes Δ*nahK* and Δ*rsmA* share. We demonstrated that the same mutants that have hyper-PYO production, Δ*nahK*, Δ*hptB*, Δ*retS* and Δ*rsmA*, also have increased Congo-Red dye uptake relative to wildtype and all other single mutant strains of the Gac-MKN system. The same trend was also observed when the Gac-MKN mutants were compared with respect to swarming motility (**Figure 2F,2G**). Here, we observed that Δ*hptB*, Δ*retS* and Δ*rsmA* all have minimal swarming motility compared to the other mutants in the Gac-MKN (**Figure 2F**,**2G**). Interestingly, while Δ*nahK* does have a significant decrease in swarming motility, it is not as drastic as Δ*hptB*, Δ*retS* and Δ*rsmA* (**Figure 2F,2G**). This may suggest that although NahK is the predominant regulator of HptB phosphorylation, alternative Gac-MKN may play additional roles in HptB-controlled swarming motility. We then examined if both Δ*hptB* and Δ*retS* could induce macrostructure formation in PA14 in the same manner as Δ*nahK* and Δ*rsmA* (**Figure 1I**) when cultured under continuous flow. Similarly to the phenotypes previously examined, Δ*nahK*, Δ*hptB*, Δ*retS*, and Δ*rsmA* are all capable of establishing mature biofilm macrostructures compared to the wildtype, which forms a flat monolayer along the surface when cultured under continuous flow (**Figure 2H**).

These characterizations revealed a unique cluster of single mutants in the Gac-MKN with similar phenotypes: Δ*nahK*, Δ*hptB*, Δ*retS*, and Δ*rsmA*. The grouping suggests that NahK, HptB, and RetS are all necessary to maintain RsmA activity to regulate these phenotypes. Complementation of the phenotypes illustrated in **Figure 2A-2G**, are shown in **Supplementary Figures S1A-S1F**; each gene and native promoter was encoded into the pUCP22 high-copy *Pseudomonas* stabilization plasmid, which returned the phenotypes to wildtype levels, demonstrating the phenotypes observed in **Figure 2** are due to the gene deletions.^45^ To determine if RsmA activity is important for modulating these responses in all hyper PYO-mutants, we then overexpressed *rsmA* in Δ*nahK*, Δ*hptB*, and Δ*retS*. Overexpression of *rsmA* in Δ*nahK*, Δ*hptB*, and Δ*retS* complements PYO production, biofilm formation, and swarming motility (**Figure 2I-2M**). In addition, overexpressing a non-functional mutant of RsmA (R44G) no longer complements the observed phenotypes.^46^ From this, we conclude the observed phenotypes are due to downstream signaling influenced by RsmA-mediated post-transcriptional regulation (**Figure 2I-2M**).

### *nahK, hptB*, and *retS* inhibit RsmA by increased transcription of *rsmY* and *rsmZ*

Once the phenotypic cluster was identified, we set out to determine the mechanism of how NahK, HptB, and RetS modulate the activity of RsmA. RetS has previously been shown to directly affect the transcription of *rsmY* and *rsmZ* by inhibition of the GacS/A signaling system through GacS heterodimerization,^21^ however, little is known about NahK or HptB downstream mechanisms. NahK is one of four kinases capable of regulating the phosphorylation state of HptB.^9^ When HptB is phosphorylated, the signal is transduced to HsbR to trigger HsbA dephosphorylation, leading to flagellar biosynthesis (**Figure 1A**). Presumably, this is an RsmA-active state, maintaining basal levels of both cyclic-di-GMP and the small RNAs *rsmY* and *rsm*Z. However, if HptB is not phosphorylated, HsbA remains phosphorylated and intracellular levels of cyclic-di-GMP elevate through stimulation of the diguanylate cyclase HsbD to increase transcription of *rsmY* (**Figure 1A**).^23^

To understand whether NahK and HptB regulate RsmA activity through the small RNAs *rsmY* and *rsmZ*, we introduced *lacZ*-transcriptional reporters for *rsmY* and *rsmZ*^5^ into PA14 wildtype, Δ*nahK*, and Δ*hptB* to quantitatively monitor each RNA’s transcription as a function of growth (**Figure 3A, 3B**). In both Δ*nahK* and Δ*hptB*, increased promoter activity for both *rsmY* and *rsmZ* relative to the wildtype was observed. As both *rsmY* and *rsmZ* were upregulated, a triple genomic deletion strain, Δ*nahK*Δ*rsmY*Δ*rsmZ*, was generated to determine the contribution of the upregulated RNAs on the previously identified RsmA-mediated phenotypes. Δ*nahK*Δ*rsmY*Δ*rsmZ* was then screened for the phenotypes observed in the Δ*nahK*, Δ*hptB*, Δ*retS*, Δ*rsmA* cluster mutants: PYO (**Figure 3C**), Congo-Red EPS visualization (**Figure 3D,3F**), and swarming motility (**Figure 3E,3G**). Interestingly, the Δ*nahK*Δ*rsmY*Δ*rsmZ* triple mutant predominantly restores PYO production and swarming motility to wildtype levels, but the Congo-Red-EPS visualization still shows increased EPS production compared to the wildtype strain (**Figure 3D,3F**). These data suggest that the ability of NahK, and therefore HptB, to inhibit RsmA is due to the increased transcription of small RNAs *rsmY* and *rsmZ*.

**Figure 3:**
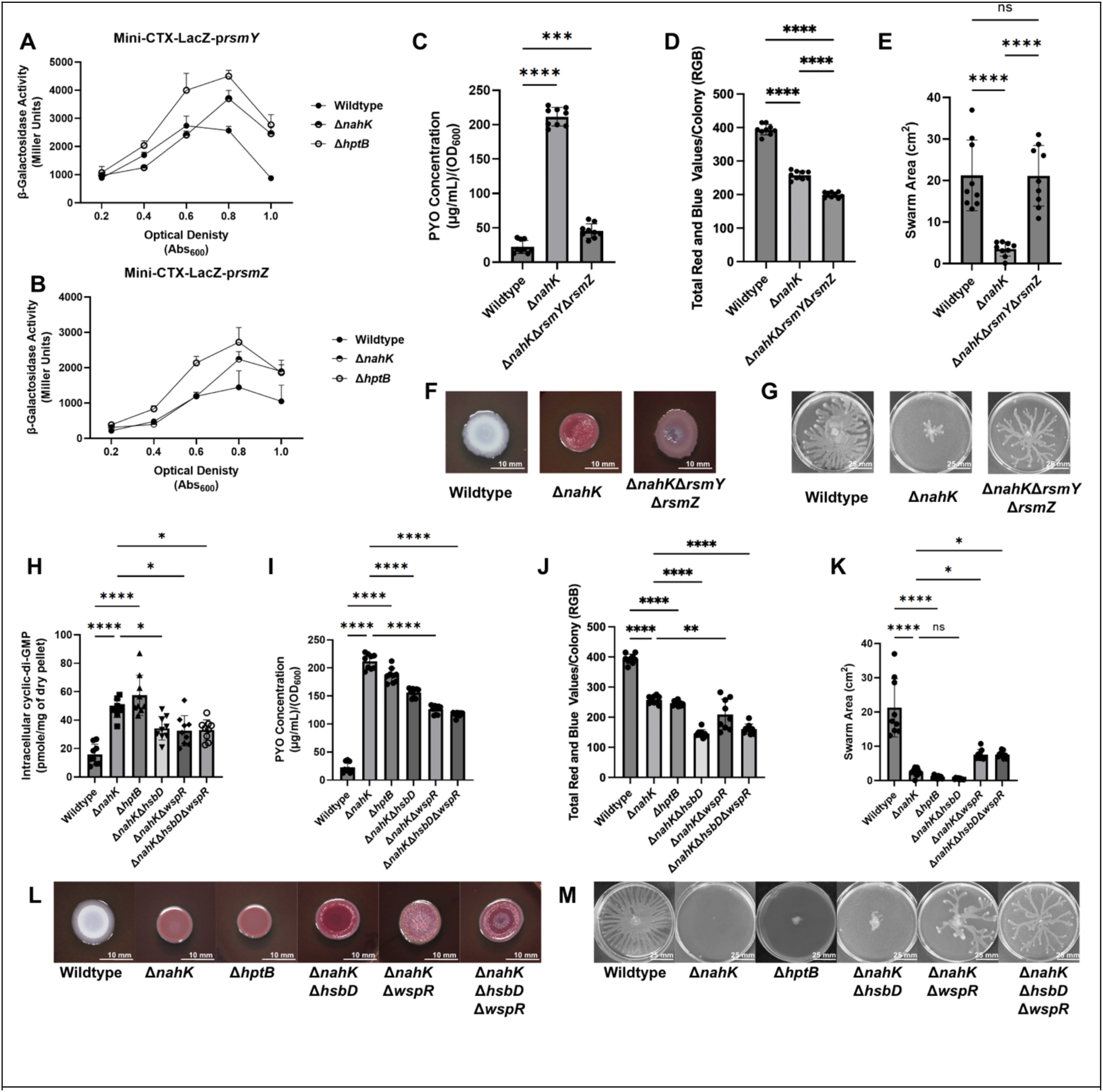
Increased transcription of the non-coding RNAs, *rsmY* and *rsmZ* lead to inactivation of RsmA in a cyclic di-GMP dependent manner. **(A)** Miller units, measuring activity of *lacZ*-promoter fusions, for *rsmY* promoter activity (**A**), and *rsmZ* promoter activity (**B**). Production of O-nitrophenol was measured at 420 nm and normalized to reaction times at each optical density; OD_600_ = 0.2, 0.4, 0.6, 0.8, and 1.0 for promoter activity. (n=3). **(C)** PYO [(μL/mL)/OD_600_)] of PA14 strains from planktonic, liquid LB cultures (n=9). **(D)** RGB quantification for EPS staining of PA14 colony biofilms cultured on 1% tryptone, 1% BactoAgar media supplemented with 40 µg/mL Congo-Red dye and 20 µg/mL Brilliant Blue dye. (n = 9). **(E)** Swarming area (cm^2^) of PA14 strains cultured on low-density agar media. (n=9). **(F)** Representative images of PA14 static Congo-Red colony biofilms. (n = 9). **(G)** Representative images of PA14 swarming motility on low-density agar (n=9). **(H)** Concentration of intracellular cyclic-di-GMP (pmol/mg of dry pellet) collected from planktonic PA14 strains cultured in M9 minimal media supplemented with 0.4% glucose. Cyclic-di-GMP extracted at OD_600_ = 0.8, isolated and quantified using HPLC. (n=9). **(I)** PYO [(μL/mL)/OD_600_)] of PA14 strains from planktonic, liquid LB cultures (n=9). **(J)** RGB quantification for EPS staining of PA14 colony biofilms cultured on 1% tryptone, 1% BactoAgar media supplemented with 40 µg/mL Congo-Red dye and 20 µg/mL Brilliant Blue dye. (n = 9). **(K)** Swarming area (cm^2^) of PA14 strains cultured on low-density agar media. (n=9). **(L)** Representative images of PA14 static Congo-Red colony biofilms. (n = 9). **(M)** Representative images of PA14 swarming motility on low-density agar (n=9). Error bars represent standard deviation from the mean. Significant differences are indicated with asterisks (N.S. *P* > 0.05; \**P* < 0.05; \*\**P* < 0.01; \*\*\**P* < 0.001; \*\*\*\**P* < 0.0001; one-way ANOVA and a Tukey multiple comparisons test).

We wondered if the intermediate EPS production for Δ*nahK*Δ*rsmY*Δ*rsmZ* could be explained by differences in how NahK and HptB influence the transcription of *rsmY* and *rsmZ*, in comparison to GacS/A. GacS/A regulate *rsmY* and *rsmZ* transcription through phosphoryl signaling, however, *rsmY* and *rsmZ* transcription can also be regulated in an uncharacterized cyclic-di-GMP-dependent manner.^47^ As a result, we extracted and quantified intracellular cyclic-di-GMP in the wildtype, Δ*nahK*, and Δ*hptB* strains (**Figure 3H**). Both Δ*nahK* and Δ*hptB* have significantly increased intracellular levels of cyclic-di-GMP compared to the wildtype. Next we wanted to determine which diguanylate cyclase enzymes were responsible for inducing these elevated levels. Previous reports linked HptB activity to two independent diguanylate cyclases, HsbD and WspR.^9,23,24^ To test if these cyclases play a role in EPS production controlled by NahK and HptB, we constructed double genomic deletions of each diguanylate cyclase in Δ*nahK* to generate Δ*nahK*Δ*hsbD* and Δ*nahK*Δ*wspR*, as well as a triple genomic deletion strain, Δ*nahK*Δ*hsbD*Δw*spR*, of both diguanylate cyclase enzymes in Δ*nahK*. Although we observed a reduction of intracellular levels of cyclic-di-GMP in the Δ*nahK*Δ*hsbD* strain (**Figure 3H**), which coincides with initial investigations into this system,^23^ increased PYO, increased EPS production and reduced motility were still observed (**Figure 3I-3M**). Reduced intracellular cyclic-di-GMP was also observed in the Δ*nahK*Δ*wspR* and Δ*nahK*Δ*hsbD*Δw*spR* mutants; however, a significant increase in swarming motility was observed. These data suggest that although NahK and HptB appear to regulate the transcription of small RNAs *rsmY* and *rsmZ* in a cyclic-di-GMP dependent manner, this regulation is not exclusively achieved through the previously characterized HsbD/WspR diguanylate cyclase enzymes. Additionally, the cyclic-di-GMP produced downstream of NahK and HptB by specific diguanylate cyclases may influence localized systems, as only reduced cyclic-di-GMP from loss of *wspR*, and not *hsbD*, affected partially restored swarming motility. Therefore, an additional diguanylate cyclase enzyme may be involved in this signaling cascade, coinciding with recent investigations into HsbA signaling through non-HsbD dependent mechanisms.^24,48^ This suggests that although NahK influences *rsmY* and *rsmZ* transcription through a HsbR/A/D/WspR dependent mechanism, alternative signaling cascades through additional diguanylate cyclases may apply additional influence on RsmA activity.

### Inter-kinase regulation tailors RsmA-mediated phenotypes in the Gac-MKN

As described, our studies have revealed a unique cluster of genes that drastically affect the activity of RsmA-mediated phenotypes: *nahK, hptB, retS*, and *rsmA*. However, the use of these mutants may not provide an accurate understanding for non-phosphoryl signaling interactions taking place between the proteins in the Gac-MKN. Several of the hybrid histidine kinases of the Gac-MKN have been shown to have additional regulatory events, including heterodimerization and small molecule stimuli. ^11-14,19,21,49^

The absence of these regulatory factors in our experiments may mask otherwise noticeable changes in RsmA-mediated phenotypes upon genomic deletion. Thus, we generated protein overexpression plasmids for each kinase involved in the Gac-MKN using the pUCP22 high copy, *Pseudomonas* stabilized vector.^45^ The promoter (−500 bp) and genetic sequences for *gacS, nahK, pa1611, retS*, and *sagS* were each ligated into the pUCP22 vector and transformed into each markerless kinase deletion mutant. We then screened the 30 newly generated overexpression strains against the three previously utilized RsmA-mediated phenotypes (**Figure 4A-E, Supplementary Figure S4A-T**). We found that fluctuations in RsmA activity were most impactful on PYO production (**Figure 4A-E**), as opposed to swarming motility (**Supplementary Figure S4B-S4C**,**S4F-S4G**,**S4J-S4K**,**S4N-S4O**,**S4R-S4S**) and Congo-Red EPS visualization (**Supplementary Figure S4A**,**S4D-S4E**,**S4H-S4I**,**S4L-S4M**,**S4P-S4Q**,**S4T**). Overproduction of PYO through the *phz2* (PYO) biosynthesis operon is uniquely and highly upregulated during RsmA inactivity, while swarming motility and Congo-Red dye uptake can be altered through additional signaling cascades outside of RsmA’s unique control.^27,50,51^

**Figure 4:**
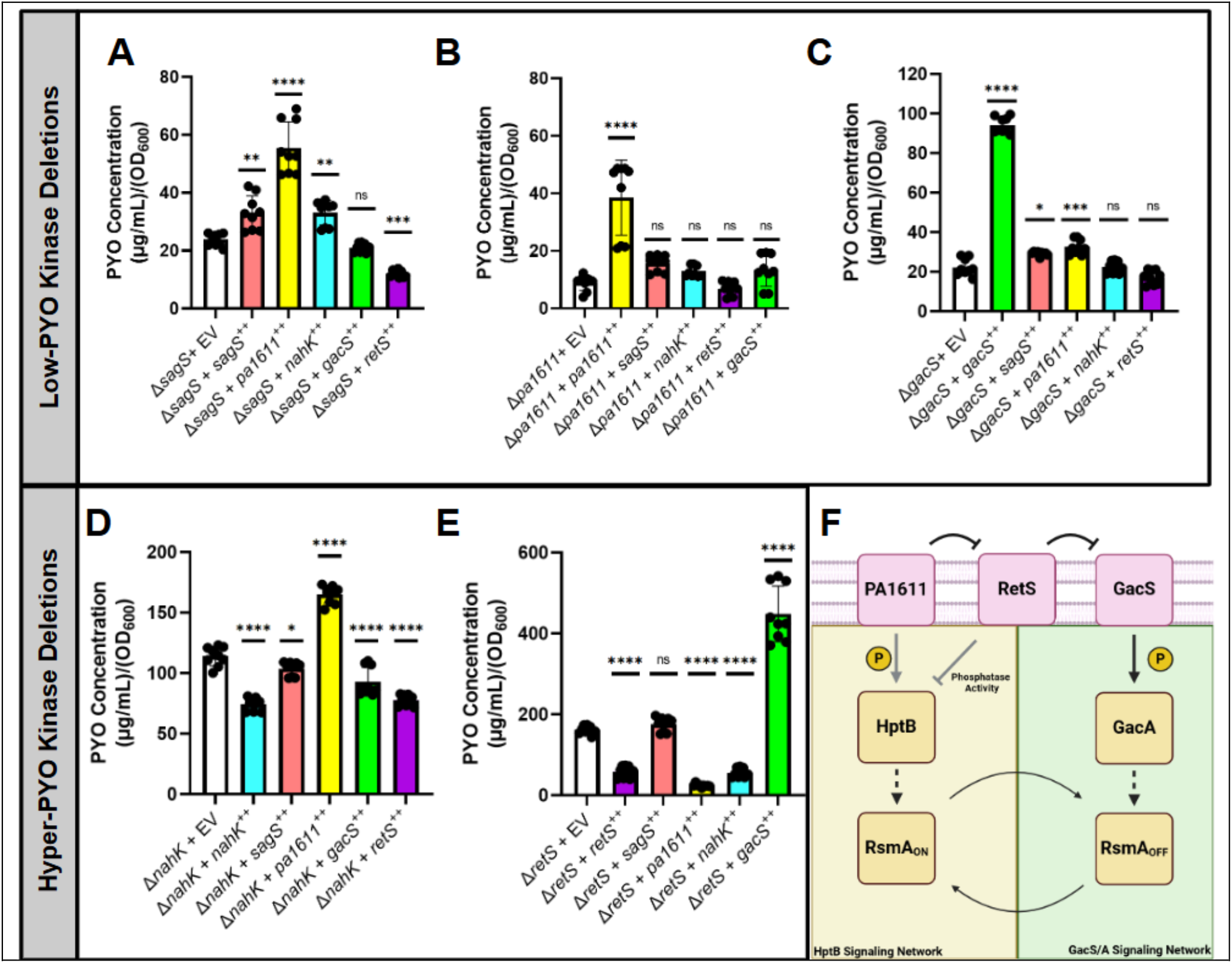
Gac-MKN heterodimerization tailors RsmA activity to control planktonic-to-biofilm switch. **(A-E)** PYO [(μL/mL)/OD_600_)] of PA14 strains from planktonic, liquid LB cultures (n=9). PA14 strains with single kinase deletions were electroporated with pUCP22 vectors containing one of the following: an empty vector (white bars), *sagS* (coral bars), *pa1611* (yellow bars), *nahK* (cyan bars), *gacS* (green bars), or *retS* (purple bars). The overexpression vectors were introduced into each single, markerless kinase deletion; **(A)** Δ*sagS*, **(B)** Δ*pa1611*, **(C)** Δ*gacS*, **(D)** Δ*nahK*, and **(E)** Δ*retS*. Error bars represent standard deviation from the mean. Significant differences are indicated with asterisks (N.S. *P* > 0.05; \**P* < 0.05; \*\**P* < 0.01; \*\*\**P* < 0.001; \*\*\*\**P* < 0.0001; one-way ANOVA and a Tukey multiple comparisons test). **(F)** Schematic representation of kinase heterodimerization within the Gac MKN. RetS mainly functions to inhibit GacS kinase activity, however, can dephosphorylate HptB in the absence of GacS. PA1611 mainly functions to inhibit RetS, however, can phosphorylate HptB in the absence of RetS.

The single markerless kinase deletion mutants were either classified as low-PYO (**Figure 4A-C**) or hyper-PYO (**Figure 4D-E**). Overexpression of the Gac-MKN kinases in the low-PYO kinase deletions (Δ*sagS*, Δ*pa1611*, and Δ*gacS*) resulted in minor, but significant fluctuations in PYO production. However, overexpression of *pa1611* resulted in increased PYO production noticeable in all low-PYO kinase deletion strains (**Figure 4A-C:yellow**). The same trend in PYO production can also be observed in the hyper-PYO kinase deletion strain, Δ*nahK*, when *pa1611* is overexpressed (**Figure 4D:yellow**). Interestingly, when *pa1611* is overexpressed in a Δ*retS* strain the increased PYO phenotype is no longer visible (**Figure 4E:yellow**), where overexpression of *pa1611* in Δ*retS* resulted in a significant decrease in PYO production. Overall, this data suggests that PA1611 may participate in two different roles depending on the presence of RetS. Previous reports have linked RetS to two functions: dephosphorylation of HptB and heterodimerization with both PA1611 and GacS.^9,21,49^ Together, this data suggests that in the presence of RetS, PA1611 may function to inhibit RetS via heterodimerization to prevent RetS-GacS heterodimerization. However, in the absence of RetS, PA1611 functions to phosphorylate HptB (**Figure 4F**). This model is further supported by overexpression of *gacS* in both the low-PYO and hyper-PYO kinase deletion kinase mutants (**Figure 4A-E:green**). In the presence of RetS-expressing genomic deletion strains (Δ*sagS*, Δ*pa1611*, and Δ*nahK*), overexpression of *gacS* results in minor-to-no change in PYO production. However, overexpression of *gacS* in a Δ*retS* strain results in a significant increase in PYO production. Interestingly, overexpression of *gacS* in a Δ*gacS* strain results in a significant increase in PYO production. This may be explained by increased promoter activity upon deletion of the *gacS* gene in an attempt to restore RsmA homeostasis by the cell. Therefore, in the presence of the pUCP22 high-copy plasmid encoding *gacS* under control of the native promoter, overproduction of GacS would result in a hyper-PYO phenotype.

Although it remains unclear how overexpression of individual kinases can slightly, but significantly, alter RsmA-mediated phenotypes, this kinase overexpression study provides clear insight into both the heterodimeric interactions taking place between PA1611, RetS, and GacS, and also provides additional evidence connecting HptB phosphorylation to the GacS/A signaling network of the Gac-MKN. Future research into the loss of kinase activity for each kinase within the Gac-MKN may help elucidate the role of each kinase in regulating RsmA activity further, as it remains unclear how loss of kinase signaling may affect signaling and promoter activity at various stages of *Pa* growth.

## Discussion

The biofilm macrostructure is a diverse landscape where signaling systems are constantly responding to external stimuli to regulate cellular events, such as the motile-to-biofilm switch. Previous investigations of the Gac-MKN have assessed the roles of various two-component signaling systems across numerous strains of *Pa*. Here, we performed a complete assessment of the Gac-MKN regulatory networks and show that histidine kinases NahK and RetS play a crucial role in regulating the motile-to-biofilm transition through the post-transcriptional regulator RsmA. Although RetS has previously been implicated in the regulation of RsmA activity, we provide the novel insight that NahK is a primary regulator of RsmA activity.

NahK is one of three hybrid histidine kinases previously identified to phosphorylate the histidine phosphotransfer protein HptB. Through *in vivo* characterization, we show that NahK has predominant control over the HptB phosphorylation state, as opposed to SagS or PA1611. Loss of *hptB* has been shown to completely abolish swarming motility and later shown to have elevated levels of intracellular cyclic-di-GMP facilitating biofilm formation.^9,22-24,44^ We propose these data can be explained by the lack of NahK-to-HptB phosphotransfer. The HptB-dependent increase in cyclic-di-GMP had previously been linked to the diguanylate cyclase HsbD,^23^ and later WspR.^24^ We show that although deletion of these diguanylate cyclases, both individually and in tandem, do reduce cyclic-di-GMP, RsmA-inhibited phenotypes are still apparent, suggesting that additional cyclic-di-GMP metabolizing enzymes may be involved in HptB-mediated RsmA inhibition (**Figure 5**).

**Figure 5:**
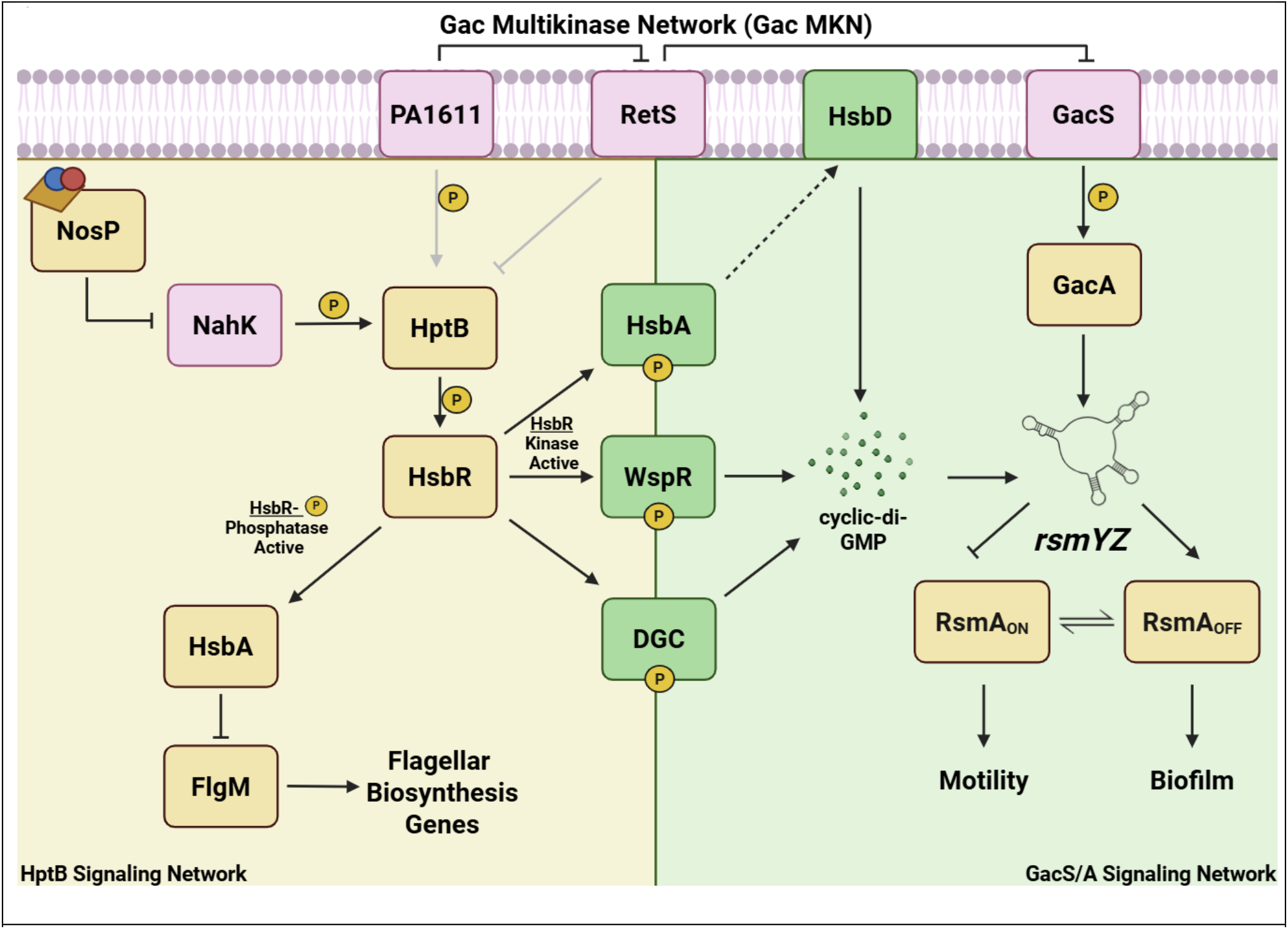
NahK is the predominant regulator of the HptB Signaling Network of the Gac MKN. Schematic summary of our investigation into Gac-MKN signal transduction. HptB branch (yellow); GacS/A branch (green). NahK has predominant control over HptB phosphorylation, resulting in complete control over RsmA activity. The phosphorylated state of HptB promotes flagellar biosynthesis through FlgM sequestration. Unphosphorylated HptB results in activation of several diguanylate cyclases to stimulate intracellular cyclic-di-GMP production to induce transcription of *rsmY* and *rsmZ*, driving RsmA equilibrium towards the RsmA_OFF_ state. GacS phosphorylates transcription factor GacA to activate transcription of both *rsmY* and *rsmZ* as well, but remains primarily inhibited by RetS. PA1611 predominantly functions to inhibit RetS, but can alternatively phosphorylate HptB in the absence of RetS. RetS primarily functions to inhibit GacS, but can dephosphorylate HptB in the absence of GacS.

The post-transcriptional regulator RsmA, regulates the translation of 500+ genes within *Pa*. This work highlights the pivotal role RsmA plays in regulating several processes involved in *Pa* pathogenicity: virulence (PYO), motility (swarming), QS (PQS), secretion systems (T3SS/T6SS), and macrostructure formation (EPS). Our laboratory has also shown that NahK-dependent regulation of RsmA activity also plays a critical role in regulating additional cellular processes, such as denitrification and NO-stress resistance.^28,52^ These systems are metabolically taxing for *Pa* to maintain over large periods of time, and explains why complete loss of *rsmA* results in an earlier onset of stationary phase (**Supplementary Figure S3**). As a result, we hypothesize that RsmA remains in a constantly changing equilibrium between the active (RsmA_ON_) and inactive (RsmA_OFF_), and regulatory mechanisms must exist to readily transition between the two states. This study, as well as others, have provided key insight into the upstream regulatory mechanims that drive RsmA_OFF_, however, it remains unknown as to how *Pa* readily converts back to the RsmA_ON_ state or how RsmA equilibrium fluctuates throughout the diverse biofilm landscape. In this study, we discovered promising preliminary evidence that negative feedback mechanisms promoting RsmA_ON_ exist, such as interconnected regulation of *rsmY* and *rsmZ*. Although these systems remain unclear through the scope of genomic deletion and gene overexpression, further exploration into these systems would pave the way for developing promising strategies to target the Gac-MKN therapeutically while simultaneously avoiding interference of feedback systems that contribute to evolutionary antimicrobial resistance.

## Materials and Methods

### Bacterial strains, growth conditions, and media

The bacterial strains and plasmids used in this study are described in **Supplemental Table 1 and 2**, respectively. The oligonucleotides used in this study are described in **Supplemental Table 3**. Unless otherwise noted, *E. coli* and *P. aeruginosa* strains were grown aerobically in Luria-Bertani (LB) medium shaking at 37 °C. Culture medium was supplemented with antibiotics when appropriate: ampicillin 100 µg/mL and gentamicin 15 µg/mL for *E. coli* and gentamicin 60 µg/mL for *P. aeruginosa*.

Growth curves were measured for all *P. aeruginosa* strains obtained, and/or generated, over the course of the study. Overnight LB cultures of *P. aeruginosa* were diluted to an initial optical density (OD_600_) of 0.01 in 5.00 mL of fresh LB media. Diluted cultures were incubated at 37 °C for 24 hours. OD_600_ was recorded every hour for the first 16 hours and then recorded every 4 hours for the remainder of the 24-hour time period.

### Generation of PA14 Markerless Deletion Strains

Markerless deletions of *P. aeruginosa* were generated as previously described with slight modification.^54^ In brief, approximately 500 base pair regions upstream and downstream were amplified and fused using overlap extension PCR. Fused inserts were ligated into the pEX18Gm *sacB* suicide plasmid. Sequence confirmed plasmids were then transformed into *E. coli* donor strain SM10 (λ*pir*) and conjugated into *P. aeruginosa* UCBPP-PA14 recipient strains through biparental mating. Conjugates were isolated onto *Pseudomonas* Isolation Agar (PIA) containing 60 µg/mL gentamicin and 25 µg/mL irgasan, where upstream or downstream chromosomal integration was confirmed through colony PCR. Upstream and downstream integration colonies were three-phase streaked onto TYS10 (10 g/L tryptone, 5 g/L yeast extract, 10% w/v filtered sucrose, 15 g/L agar) and incubated at room temperature for 48 hours to allow for *sacB*-mediated counter selection to occur, followed by a 24-hour incubation at 37 °C to facilitate colony growth. Markerless deletions were confirmed through colony PCR, followed by DNA sequencing of the markerless genomic deletion region.

### Pyocyanin extraction

PYO extraction was performed as previously described.^27,53^ In brief, overnight cultures were diluted 100-fold into 100 mL of fresh LB media, where cultures were incubated for 16 hours at 37 °C. After 16 hours, OD_600_ was recorded, and cells were pelleted, retaining the PYO-containing supernatant. Chloroform was used to extract all phenazines from the bacterial supernatant. PYO was then isolated from the chloroform abstracts with 0.1 M HCl for quantification. Electron absorption spectrums of acidified PYO extracts were collected on a Varian Cary-100 Bio Spectrophotometer. PYO was quantified utilizing the Beer’s Lambert equation (extinction coefficient: 17.072 µg/mL.^53^ PYO quantification in the presence of M64 PqsR inhibitor were cultured in LB medium supplemented with 1 µM of M64 PqsR inhibitor.

### Transcriptional mScarlet Fluorescence Assay

For reporter assays, *P. aeruginosa* strains containing chromosomally integrated mScarlet *phz1* or *phz2* reporters were grown overnight in 5 mL LB. After 16 hours, the overnight cultures were then diluted 100-fold into black-walled, clear-bottom 96-well plated containing 200 μL succinate minimal media. Over 24 hours, mScarlet fluorescence (ex: 560 nm/em: 610 nm) and OD_600_ were recorded kinetically every 30 mins on a SpectraMax iD3 Multimode Plate Reader. For coculture assays, strains were mixed in a 1:1 ratio of fluorescent, mScarlet-expressing and non-fluorescent cells. Over 24 hours, mScarlet fluorescence (ex: 560 nm, em: 610 nm) and OD_600_ were recorded kinetically every 30 mins on a SpectraMax iD3 Multimode Plate Reader. Plates were continuously shaking in an orbital direction at 37 °C.

### Congo-Red Assay

Congo Red dye has previously been reported for the qualitative visualization of *P. aeruginosa* biofilm and EPS production.^34^ A mixture of 1% tryptone and 1% Bacto Agar was autoclaved and allowed to cool for 20 minutes prior to supplementation of 40 µg/mL Congo Red (MP Biomedical) and 20 µg/mL Brilliant Blue G (ACROS Organics). The final mixture was poured in 25.00 mL volumes into 100 x 15 mm sterile petri dishes. LB overnight cultures of *P. aeruginosa* were directly pipetted in 10 µL volumes onto the Congo Red/Brilliant Blue G containing tryptone medium. Plates were incubated at room temperature for 72 hours in total darkness to stimulate the production of cyclic-di-GMP prior to imaging. After 72 hours, plates were imaged and RGB values determined using ImageJ image processing software.

### Swarming Motility Assay

*P. aeruginosa* strains were grown in LB media at 37 °C shaking. After 16 hours, 40 µL of each overnight culture was spread across LB agar medium and incubated at 37 °C to form bacterial lawns of *P. aeruginosa* to encourage communal behavior. *P. aeruginosa* strains containing pUCP22-based overexpression plasmids were spread across LB agar medium supplemented with 60 µg/mL gentamicin. A sterile pipette tip was then used to transfer a 1 cm section of bacterial lawn onto the center of a swarming motility plate (4 mM NH_4_Cl, 2.4 mM Na_2_HPO_4_, 4.4 mM KH_2_PO_4_, 1.71 mM NaCl) containing 0.5% Difco agar, supplemented with 11 mM dextrose, 0.5% casamino acids, 1 mM MgSO_4_, and 0.1 mM CaCl_2_. The spotted cultures were then incubated at 30 °C for 16 hours to allow for bacterial swarming to occur. After 16 hours, swarming motility plates were imaged, and the area of total swarming motility was calculated utilizing ImageJ image processing software.

### Continuous Flow Microscopy

The flow cell system was adapted from previously published methods.^55^ Overnight cultures of *P. aeruginosa* were grown in LB medium at 37 °C shaking. After 16 hours, overnight cultures of *P. aeruginosa* were diluted 100-fold into fresh LB media and incubated at 37 °C until an OD_600_ of 0.6 was obtained. The newly cultured *P. aeruginosa* strain was then diluted to an OD_600_ of 0.05 (inoculum). Using a sterile syringe and needle, 150 µL of inoculum was injected into a 40 mm x 4 mm x 1 mm three-channel flow cell chamber preequilibrated with 1% tryptic soybean broth (TSB) medium at room temperature. The inoculated flow cell chamber was incubated for 1 hour in an inverted position to allow for bacterial attachment to the adhered glass cover slip. After 1 hour, the flow cell chamber was reverted to the upright position and supplied with 1% TSB medium at a constant flow rate of 3 mL/hour using a Cole-Parmer peristaltic pump (Cole Parmer Instrument Co., Germany).

After 72 hours of culturing under continuous flow, the flow of 1% TSB medium was stopped and bacterial cells were stained using the nucleic acid stain SYTO62 (Molecular Probes, Eugene, Oreg.) SYTO62 was diluted to a final concentration of 0.1 mM in 1% TSB medium. *P. aeruginosa* biofilms were stained for 30 mins in the dark before the flow of 1% TSB medium was then resumed to remove excess SYTO62 red dye for 1 hour at a flow rate of 5 mL/hour. The biofilms were observed using a Zeiss AXIO Examiner D1 microscope equipped with an ANDOR DSD2 laser free confocal module. *P. aeruginosa* biofilms cultured under continuous flow were imaged in triplicate through a 20x/0.80 DICII objective lens and an X-Cite 120LED mini–LED Illuminator to excite SYTO62 fluorescence. The acquired images were further assessed on ImageJ.

### RNA Sequencing

Overnight cultures of *P. aeruginosa* strains were grown in Luria-Bertani (LB) media at 37 °C shaking. After 16 hours, the overnight cultures were diluted 25-fold into 5.00 mL of fresh 1X M9 minimal media supplemented with 0.4% glucose. The M9 cultures were incubated at 37 °C shaking to an OD_600_ of 0.45; then, each culture was subjected to 3 hours of iron chelation with the addition of 500 µM of 2-2’ bipyrodile. After 3 hours, the cultures were diluted 50-fold into 50 mL of fresh 1X M9 minimal media (0.4% glucose) and grown at 37 °C shaking for 16 hours. Each *P. aeruginosa* strain was grown in biological duplicate. RNAprotect® Bacterial Reagent was used to prevent RNA degradation prior to RNA extraction. Total RNA was extracted using the QIAGEN RNeasy® Mini Kit. RNA quality was assessed utilizing a BioAnalyzer 2100 prior to RNA sequencing. RNA sequencing was performed on an Illumina NextSeq 550 on a Mid Output 150 cycle flow cell kit assembly at the Stony Brook University Genomics and Bioinformatics Core Facility. Illumina-generated sequencing reads were later processed and mapped to the *P. aeruginosa* UCBPP-PA14 genome sequence from the *Pseudomonas* Genome Database (www.pseudomonas.com).

### LacZ Beta-Galactosidase Assay

For beta-galactosidase quantifications, *P. aeruginosa* strains containing chromosomally integrated mini-CTX-*lacZ*, mini-CTX-*lacZ-*p*rsmY* and -p*rsmZ* constructs were grown in 5.00 mL LB overnight. After 16 hours, *P. aeruginosa* cultures were diluted to an initial OD_600_ = 0.01 into fresh M9 minimal media supplemented with 0.4% glucose and incubated at 37 °C shaking. At OD_600_ = 0.2, 0.4, 0.6, 0.8 and 1.0, 20 μL of the shaking cultures were added to 80 μL of permeabilization solution (100 mM Na_2_HPO_4,_ 20 mM KCl, 2mM MgSO_4_, 0.8 mg/mL CTAB, 0.4 mg/mL deoxycholate and 5.4 μL/ml BME). After the last 20 μL sample was added to the permeability solution, 600 μL of the substrate solution (60 mM Na_2_HPO4, 40 mM NaH_2_PO4, 1 mg/ml ONPG, and 2.7 μL/mL BME) was added to each sample and the time was noted. Once the samples developed into a noticeable yellow hue, indicating ONPG conversion to o-nitrophenol, 700 μL of the stop solution (1M Na_2_CO_3_) was immediately added, and the total reaction time was recorded. Samples were pelleted (14,000 RPM, 5 mins), and the absorbances of the supernatants were analyzed at 420 nm.

### Cyclic-di-GMP Extraction

For intracellular cyclic-di-GMP extractions, *P. aeruginosa* overnight strains were diluted to an OD_600_ of 0.05 in M9 minimal media supplemented with 0.4% glucose. Cultures were grown to an OD_600_ of 0.8, where 10 mL of the bacterial culture was pelleted in triplicate (4 °C, 2500 RPM, 20 mins). The supernatant was discarded, and pellets were resuspended in 300 μL of ice-cold extraction buffer (40% MeOH, 40% ACN, 20% H_2_O). Resuspended cells were incubated at −20 °C for 15 minutes and were immediately heated to 95 °C for 3 minutes to denature cyclic-di-GMP modulating enzymes. The samples were then cooled on ice for 5 minutes prior to centrifugation at 14000 RPM for 10 minutes. The supernatant was transferred to new 1.5 mL microcentrifuge tubes. Cell pellets were resuspended and extracted twice with 200 μL of extraction buffer, excluding the 95 °C heating step. Combined cell extracts were collected in a new 1.5 mL microcentrifuge tube, and the extraction buffer was removed under vacuum using a centrifugal evaporator. Dried cell pellets from extraction were also dried and weighed.

Solid cell extracts were resuspended in 120 μL of deionized water, followed by filtration of the supernatant through a 0.22 μm hydrophilic filter. 60 μL of each sample was injected into a reverse-phase Phenomenex Luna Omega Polar C-18 column (250 mm x 4.6 mm x 5 μm particle size). Samples were analyzed by a Shimadzu LC-2010A HT high-performance liquid chromatography that was equilibrated in running buffer (15 mM triethylammonium acetate in 6% methanol [pH 5.0, 30 °C]) at a flow rate of 0.75 ml min^-1^ for 100 min. Intracellular cyclic-di-GMP concentrations were calculated using the respective peak areas from the samples and were calibrated to a known cyclic-di-GMP standard curve. Final values were normalized to the weight of dried cell pellets from each sample.

## Acknowledgments

We thank B. Tseng for helpful insight into continuous flow microscopy and experimental design. We thank L.E.P. Dietrich for helpful discussion of the manuscript. We thank S. Laughlin and members of the S. Laughlin laboratory for training and access to confocal laser scanning microscope utilized in imaging bacterial macrostructures. We thank K. Oasima, S. Multani, and G. Sharma for early protocol development for the phenotypic screening of RsmA phenotypes. The RNA sequencing studies were supported by the Stony Brook University Renaissance School of Medicine Genomics Core Facility. This study was supported by a National Institute of Health R01GM118894 (to E.M.B.). J.M.W., K.R., and A.G.M. were supported by the National Institutes of Health T32 training program (T32GM136572). J.M.W. was supported by a Marcus and Kimberley Boehm Fellowship. A.M. was supported by a National Science Foundation REU program (CHE-2050541). D.G. was supported by the National Institutes of Health IRACDA training program (K12GM102778).

## Author Contributions

J.M.W., K.R., A.G.M., and E.M.B. conceived and designed the study, analyzed data, and wrote the manuscript; J.M.W., K.R., and N.E.A. contributed to various aspects of phenotypic studies; J.M.W., A.G.M., and J.F. prepared samples and carried out data analyses for RNA sequencing experiments; J.M.W. and K.R. prepared samples for *in vitro* quantification of promoter activity and secondary messengers; J.M.W., K.R., J.F., and A.M. contributed to the generation of in-frame markerless deletions; J.M.W. and N.E.A. constructed protein overexpression plasmids; J.M.W., D.G., and K.E.B. contributed to experimental design for phenotypic screening; J.M.W., K.R., and N.E.A carried out all other experiments and data analysis; A.M. contributed to manuscript editing; E.M.B. led and coordinated the project and was PI on the grants funding the project.

