## Supplementary Materials for "Pseudomonas aeruginosa uses kinases NahK and RetS to control the motile-biofilm switch"

**Supplementary Table S1:** Strains used in this study.

| Strain | Relevant Characteristics | Source |
|---|---|---|
| ***P. aeruginosa*** | | |
| UCBPP-PA14 (PA14) | Wildtype | Lab Collection |
| PA14 Δ*gacA* | Markerless, in-frame deletion of PA14_30650 (*gacA*) in PA14 | [13] |
| PA14 Δ*gacS* | Markerless, in-frame deletion of PA14_52260 (*gacS*) in PA14 | [13] |
| PA14 Δ*hptB* | Markerless, in-frame deletion of PA14_20800 (*hptB*) in PA14 | Generous donation from the Eric Deziel Laboratory at Institut National de la Recherche Scientifique |
| PA14 Δ*nahK* | Markerless, in-frame deletion of PA14_38970 (*nahK*) in PA14 | [27] |
| PA14 Δ*nahK*Δ*hsbD* | Markerless, in-frame deletion of PA14_38970 (*nahK*) and PA14_20820 (*hsbD*) in PA14 | This study |
| PA14 Δ*nahK*Δ*rsmY*Δ*rsmZ* | Markerless, in-frame deletion of PA14_38970 (*nahK*), PA14_06875 (*rsmY*), and PA3621.1 (*rsmZ*) in PA14 | This study |
| PA14 Δ*nahK*Δ*wspR* | Markerless, in-frame deletion of PA14_38970 (*nahK*) and PA14_16500 (*wspR*) in PA14 | This study |
| PA14 Δ*nosP* | Markerless, in-frame deletion of PA14_38990 (*nosP*) in PA14 | This study |
| PA14 Δ*nosP*Δ*nahK* | Markerless, in-frame deletion of PA14_38990 (*nosP*) and PA14_38970 (*nahK*) in PA14 | Generous donation from the Lars E.P. Dietrich Laboratory at Columbia University |
| PA14 Δ*pa1611* | Markerless, in-frame deletion of PA14_43670 (*pa1611*) in PA14 | This study |
| PA14 Δ*retS* | Markerless, in-frame deletion of PA14_64230 (*retS*) in PA14 | [13] |
| PA14 Δ*rsmA* | Markerless, in-frame deletion of PA14_52570 (*rsmA*) in PA14 | Generous donation from the Lars E.P. Dietrich Laboratory at Columbia University |

| | | |
|---|---|---|
| PA14 Δ*rsmY* | Markerless, in-frame deletion of PA14_06875 (*rsmY*) in PA14 | Generous donation from the Lars E.P. Dietrich Laboratory at Columbia University |
| PA14 Δ*rsmZ* | Markerless, in-frame deletion of PA3621.1 (*rsmZ*) in PA14 | Generous donation from the Lars E.P. Dietrich Laboratory at Columbia University |
| PA14 Δ*rsmY*Δ*rsmZ* | Markerless, in-frame deletion of PA14_06875 (*rsmY*) and PA3621.1 (*rsmZ*) in PA14 | Generous donation from the Lars E.P. Dietrich Laboratory at Columbia University |
| PA14 Δ*sagS* | Markerless, in-frame deletion of PA14_27550 (*sagS*) in PA14 | This study |
| PA14 Δ*pqsABC*::P*phz1* - mScarlet | Markerless, in-frame deletion of PA14_51430-51410 (*pqsABC*) with Pphz1-mScarlet integrated at the attB site using pLD3208 | [27] |
| PA14 Δ*pqsABC*::P*phz2* - mScarlet | Markerless deletion of PA14_51430-51410 (*pqsABC*) with Pphz2-mScarlet integrated at the attB site using pLD3208 | [27] |
| PA14 Δ*pqsR*::P*phz1* - mScarlet | Markerless, in-frame deletion of PA14_51340 (*pqsR*) with Pphz1-mScarlet integrated at the attB site using pLD3208 | [27] |
| PA14 Δ*pqsR*::P*phz2* - mScarlet | Markerless, in-frame deletion of PA14_51340 (*pqsR*) with Pphz2-mScarlet integrated at the attB site using pLD3208 | [27] |
| ***E. coli*** | | |
| DH5α | *fhuA2 lac(del)U169 phoA glnV44 Φ80' lacZ(del)M15 gyrA96 recA1 relA1 endA1 thi-1 hsdR17* | Lab Collection |
| WM3064 | *thrB1004 pro thi rpsL hsdS lacZΔM15 RP4-1360 Δ(araBAD)567 ΔdapA1341::[erm pir]* | Lab Collection |
| S17-1 | *StrR, TpR, F-RP4-2-Tc::Mu aphA::Tn7 recA λpir lysogen* | Generous donation from the Lars E.P. Dietrich Laboratory at Columbia University |
| SM10 | *thi thr leu tonA lacY supE recA::RP4-2-Tc::Mu Km λpir* | Generous donation from the Herbert P. Schweizer Laboratory at Northern Arizona University |

**Supplementary Table S2:** Plasmids used in this study.

| Plasmid | Relevant Characteristics | Source |
|---|---|---|
| pEX18Gm | Suicide vector in *P. aeruginosa SacB*, $Gm^R$ | Generous donation from the Herbert P. Schweizer Laboratory at Northern Arizona University |
| pEX18Gm Δ*nahK* | Suicide vector for Δ*nahK* mutation construction | This study |
| pEX18Gm Δ*hsbD* | Suicide vector for Δ*hsbD* mutation construction | This study |
| pEX18Gm Δ*wspR* | Suicide vector for Δ*wspR* mutation construction | This study |
| pEX18Gm Δ*nosP* | Suicide vector for Δ*nosP* mutation construction | This study |
| pEX18Gm Δ*pa1611* | Suicide vector for Δ*pa1611* mutation construction | This study |
| pEX18Gm Δ*sagS* | Suicide vector for Δ*sagS* mutation construction | This study |
| pFLP2 | Site-specific excision vector with cI857-controlled FLP recombinase encoding sequence, sacB, AmpR. | Generous donation from the Herbert P. Schweizer Laboratory at Northern Arizona University |
| pUCP22 | High-copy *P. aeruginosa* stabilized vector, $Amp^R$,$Gm^R$ | Generous donation from the Herbert P. Schweizer Laboratory at Northern Arizona University |
| pUCP22 PA14 *gacA* | High-copy *P. aeruginosa* stabilized vector containing PA14 *gacA* + 494 bp upstream, $Amp^R$,$Gm^R$ | This study |
| pUCP22 PA14 *gacS* | High-copy *P. aeruginosa* stabilized vector containing PA14 *gacS* + 443 bp upstream, $Amp^R$,$Gm^R$ | This study |
| pUCP22 PA14 *hptB* | High-copy *P. aeruginosa* stabilized vector containing PA14 *hptB* + 500 bp upstream, $Amp^R$,$Gm^R$ | This study |
| pUCP22 PA14 *nahK* | High-copy *P. aeruginosa* stabilized vector containing PA14 *nahK* + 500 bp upstream *nosP*, $Amp^R$,$Gm^R$ | This study |
| pUCP22 PA14 *nosP* | High-copy *P. aeruginosa* stabilized vector containing | This study |

| | | |
|---|---|---|
| | PA14 *nosP* + 500 bp upstream, $Amp^R$,$Gm^R$ | |
| pUCP22 PA14 *nosP nahK* | High-copy *P. aeruginosa* stabilized vector containing PA14 *nosP nahK* + 500 bp upstream, $Amp^R$,$Gm^R$ | This study |
| pUCP22 PA14 *pa1611* | High-copy *P. aeruginosa* stabilized vector containing PA14 *pa1611* + 465 bp upstream, $Amp^R$,$Gm^R$ | This study |
| pUCP22 PA14 *retS* | High-copy *P. aeruginosa* stabilized vector containing PA14 *retS* + 500 bp upstream, $Amp^R$,$Gm^R$ | This study |
| pUCP22 PA14 *rsmA* | High-copy *P. aeruginosa* stabilized vector containing PA14 *rsmA* + 500 bp upstream, $Amp^R$,$Gm^R$ | This study |
| pUCP22 PA14 *rsmA* R44G | High-copy *P. aeruginosa* stabilized vector containing PA14 *rsmA* R44G + 500 bp upstream, $Amp^R$,$Gm^R$ | This study |
| pUCP22 PA14 *rsmY* | High-copy *P. aeruginosa* stabilized vector containing PA14 *rsmY* + 455 bp upstream, $Amp^R$,$Gm^R$ | This study |
| pUCP22 PA14 *rsmZ* | High-copy *P. aeruginosa* stabilized vector containing PA14 *rsmZ* + 457 bp upstream, $Amp^R$,$Gm^R$ | This study |
| pUCP22 PA14 *sagS* | High-copy *P. aeruginosa* stabilized vector containing PA14 *sagS* + 447 bp upstream, $Amp^R$,$Gm^R$ | This study |

**Supplementary Table S3:** Oligonucleotides used in this study.

| Oligonucleotide | Oligonucleotide Sequence |
|---|---|
| ***Markerless Deletions*** | |
| Δ*hsbD* Upstream Fwd | 5'-catgaattctggcccaggtcagcg-3' |
| Δ*hsbD* Upstream Rev | 5'-ccatggactcaggccaccacgcacacctcttctcttgg-3' |
| Δ*hsbD* Downstream Fwd | 5'-ccaagagaagaggtgtgcgtggtggcctgagtccatgg-3' |
| Δ*hsbD* Downstream Rev | 5'-gataagcttcctggcgcagttcg-3' |
| Δ*hsbD* Upstream In check Rev | 5'-gccagccagacgagcatgc-3' |
| Δ*hsbD* Downstream In check Fwd | 5'-ctgtcggtcggcgtgtcc-3' |
| Δ*hsbD* Final Out Check Fwd | 5'-gacctcaagccacagc-3' |
| Δ*hsbD* Final Out Check Rev | 5'-caccttctgcgcatagac-3' |
| Δ*nahK* Upstream Rev | 5'-cgcctgaatcagaccgagcctgcagctcagcg-3' |
| Δ*nahK* Downstream Fwd | 5'-cgctgagctgcaggctcggtctgattcaggcg-3' |
| Δ*nahK* Downstream Rev | 5'-cataagcttcggtttcctgctggc-3' |
| Δ*nahK* Upstream In check Rev | 5'-gtccgatcagcgcctggttg-3' |
| Δ*nahK* Downstream In check Fwd | 5'-gatgatcaccgccaactacagcaac-3' |
| Δ*nahK* Final Out Check Fwd | 5'-tcgtgctgttcttctgttcc-3' |
| Δ*nahK* Final Out Check Rev | 5'-gtcgaacaagcagatcgc-3' |
| Δ*pa1611* Upstream Fwd | 5-cattctagagtgctctacggcaaggacag-3' |
| Δ*pa1611* Upstream Rev | 5'-ccttgtcgtagatcattccggctggctcatggttcttctccttc-3' |
| Δ*pa1611* Downstream Fwd | 5'-gaaggagaagaaccatgagccagccggaatgatctacgacaagg-3' |
| Δ*pa1611* Downstream Rev | 5'-cataagcttgttgccctgccagcc-3' |
| Δ*pa1611* Upstream In Check Rev | 5'-gtgagcaacagggtcagcagc-3' |
| Δ*pa1611* Downstream In Check Fwd | 5'-caacgcatactgcaacgctggatcg-3' |
| Δ*pa1611* Out Check Fwd | 5'-gtggaactgcgcacgc-3' |
| Δ*pa1611* Out Check Rev | 5'-gaggtgaacaggccgac-3' |
| Δ*sagS* Upstream Fwd | 5'-catgaattcgcagttgctcgaccacgc-3' |
| Δ*sagS* Upstream Rev | 5'-gcaccagcggtcgaggatcagccttggcgag-3' |
| Δ*sagS* Downstream Fwd | 5'-ctcgccaaggctgatcctcgaccgctggtgc-3' |
| Δ*sagS* Downstream Rev | 5'-cgatctagacaaccagctgtgtttcaag-3' |
| Δ*sagS* Upstream In check Rev | 5'-gatggatgcgtcctggctg-3' |
| Δ*sagS* Downstream In Check Fwd | 5'-ccggcatggacgactacctgg-3' |
| Δ*sagS* Final Out Check Fwd | 5'-ccagtagcgggagaac-3' |
| Δ*sagS* Final Out Check Rev | 5'-gatggcgtcgaagaatc-3' |
| pEX18Gm *genR* Upstream IN Fwd | 5'-caacgcgcttggtgcttatgtgatc-3' |
| pEX18Gm *sacB* Downstream IN Rev | 5'-gagttgcgcctcctgccag-3' |
| ***RT-qPCR*** | |
| qPCR *proC* Fwd | 5'-caggccgggcagttgctgtc-3' |
| qPCR *proC* Rev | 5'-ggtcaggcgcgaggctgtct-3' |
| qPCR *rsmY* Fwd | 5'-ggaagcgccaaagacaata-3' |

| | |
|---|---|
| qPCR *rsmY* Rev | 5'-gcagacctctatcctgacat-3' |
| qPCR *rsmZ* Fwd | 5'-gtacagggaacacgcaac-3' |
| qPCR *rsmZ* Rev | 5'-tcttcagtccctcgtcatc-3' |
| ***Complementation*** | |
| pUCP22 *gacA* Fwd | 5'-catggtaccaagcaatcctggatcgtcgg-3' |
| pUCP22 *gacA* Rev | 5'-cataagcttggcgctcatctagctggc-3' |
| pUCP22 *gacS* Fwd | 5'-catggtacccgacatcaggatcaccggc-3' |
| pUCP22 *gacS* Rev | 5'-cataagctttcagagttcgctggagtcg-3' |
| pUCP22 *hptB* Fwd | 5'-gtaggtaccccccatgtgctgcaactg-3' |
| pUCP22 *hptB* Rev | 5'-cataagcttggacgaaaacctcagcgatag-3' |
| pUCP22 *nosP* nahK Fwd | 5'-catcccgggtgacttccaagggggtcg-3' |
| pUCP22 *nosP* nahK Rev | 5'-cataagctttcagaccgaggcttcgc-3' |
| pUCP22 *nosP* Q5 Fwd | 5'-aagcttggcactggc-3' |
| pUCP22 *nosP* Q5 Rev | 5'-tcagcgacggacacc-3' |
| pUCP22 *nahK* Q5 Fwd | 5'-atggcatgcatatcaaccagac-3' |
| pUCP22 *nahK* Q5 Rev | 5'-gtacatgcccgctccc-3' |
| pUCP22 *pa1611* Fwd | 5'-catcccggggctctacggcaaggacag-3' |
| pUCP22 *pa1611* Rev | 5'-cataagcttgcagcctagtcatagactgttgc-3' |
| pUCP22 *retS* Fwd | 5'-gtaggtacccatggtccgcctggagtcc-3' |
| pUCP22 *retS* Rev | 5'-cataagctttcaggagggcagggcg-3' |
| pUCP22 *rsmA* Fwd | 5'-catggtacccttcacggtgcatcg-3' |
| pUCP22 *rsmA* Rev | 5'-cataagcttttaatggtttggctcttgatc-3' |
| pUCP22 *rsmY* Fwd | 5'-catggtacccgaaggtgttgcgttcgttgg-3' |
| pUCP22 *rsmY* Rev | 5'-cataagcttgctcgctgctggaaggc-3' |
| pUCP22 *rsmZ* Fwd | 5'-catgaattctggctgacggaactcacc-3' |
| pUCP22 *rsmZ* Rev | 5'-cattctagaggcgacgagtaaaacggc-3' |
| pUCP22 *sagS* Fwd | 5'-catggtacccgctcgatctcgatcagg-3' |
| pUCP22 *sagS* Rev | 5'-cataagcttctggtggtgggaggctag-3' |
| ***Site Directed Mutagenesis*** | |
| pUCP22 *rsmA* R44G Fwd | 5'-gcggaggaaatttaccagcgc-3' |
| pUCP22 *rsmA* R44G Rev | 5'-gtgcacggcgacttcc |

**Supplementary Table S4:** *Transcriptome analysis of PA14 WT, PA14 ΔnahK, and PA14 ΔrsmA in M9 minimal media supplemented with 0.4% glucose*

**See attached file*

**Supplementary Table S5:** *Transcriptome analysis of PA14 WT and PA14 ΔnosPΔnahK in Luria-Bertani (LB) and succinate-minimal media (SMM).*

**See attached file*

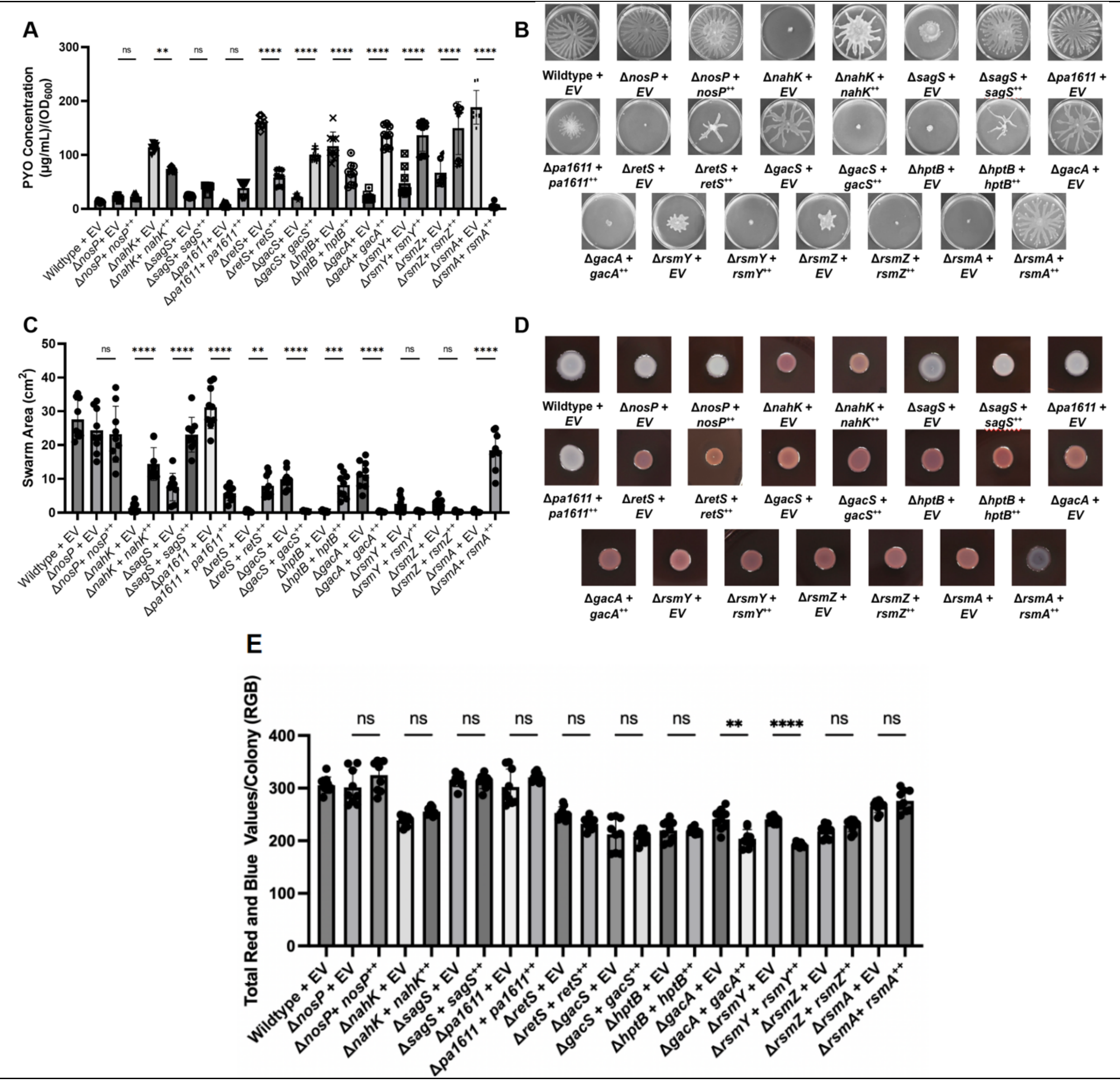


**Supplementary Figure S1: Genomic deletion of genes within the Gac MKN are complemented with pUCP22:** **(A-E)** Phenotypic characterization for RsmA-mediated phenotypes of PA14 strains containing pUCP22 overexpression vector of either the empty plasmid (EV) or the Gac MKN genomic deletion complement. **(A)** PYO [(μL/mL)/$OD_{600}$)] of PA14 strains from planktonic, liquid LB cultures (n=9). **(B)** Representative images of PA14 swarming motility on low-density agar (n=9). **(C)** Swarming area ($cm^2$) of PA14 strains cultured on low-density agar media. (n=9). **(D)** Representative images of PA14 static Congo Red colony biofilms. (n = 9). **(E)** RGB quantification for EPS staining of PA14 colony biofilms cultured on 1% tryptone, 1% BactoAgar media supplemented with 40 μg/mL Congo Red dye and 20 μg/mL Brilliant Blue dye. (n = 9). Error bars represent standard deviation from the mean. Significant differences are indicated with asterisks (N.S. $P > 0.05$; *$P < 0.05$; **$P < 0.01$; ***$P < 0.001$; ****$P < 0.0001$; one-way ANOVA and a Tukey multiple comparisons test).

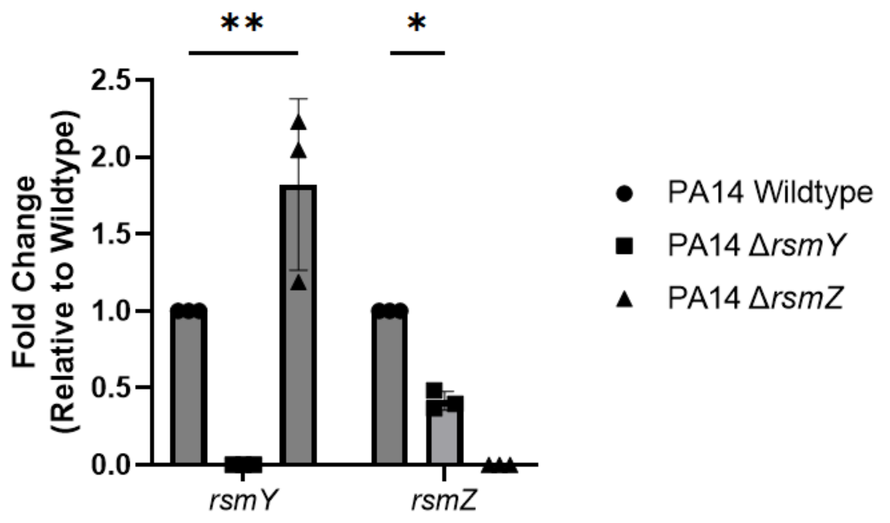


**Supplementary Figure S2: Single genomic deletion of non-coding regulatory RNAs interferes with transcription of corresponding non-coding regulatory RNA.** Qualitative PCR (qPCR) for *rsmY* and *rsmZ* in Δ*rsmY* and Δ*rsmZ* compared to wildtype. *proC* (delta 1-pyrroline-5-carboxylate reductase) was used as a housekeeping gene, and the fold change is reported relative to its expression in the wildtype control. p-values were calculated using two-way ANOVA and a Tukey multiple comparisons test (n = 3). ** $p \leq 0.01$, * $p \leq 0.05$. Error bars represent standard deviation from the mean. Significant differences are indicated with asterisks (*$P < 0.05$; **$P < 0.01$; one-way ANOVA and a Tukey multiple comparisons test).

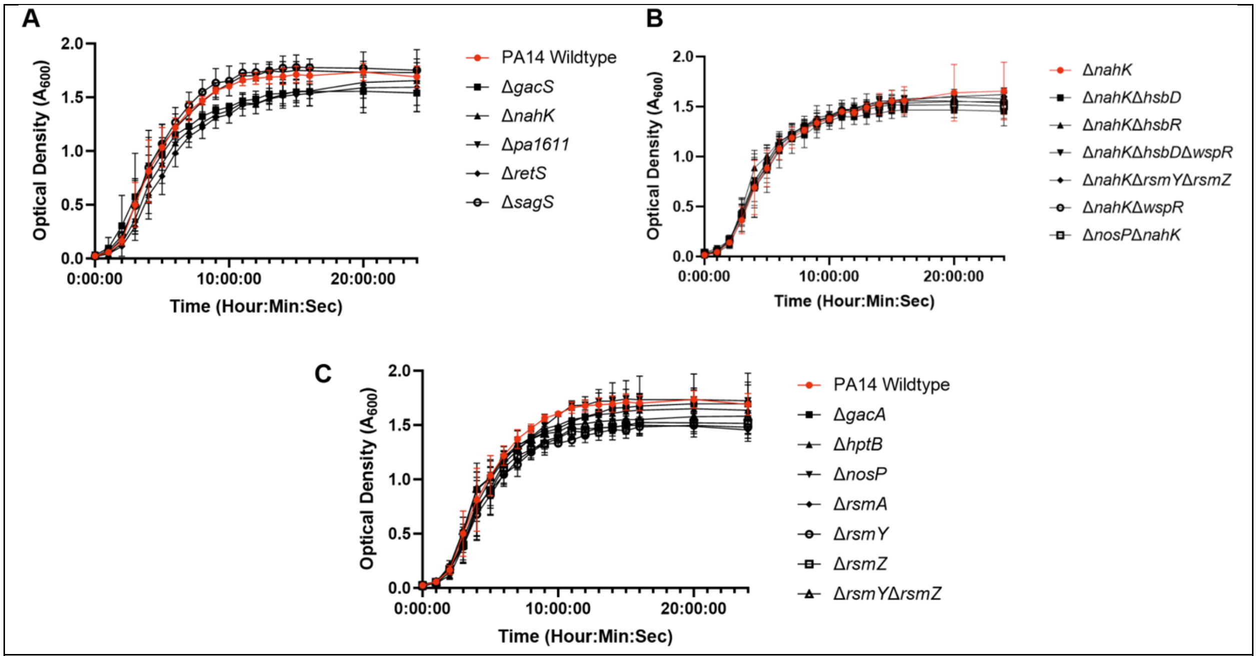


**Supplementary Figure S3: (A)** 24-hour growth curves of *P. aeruginosa* wildtype and single Gac MKN kinase markerless genomic deletions. *P. aeruginosa* UCBPP-PA14 wildtype colored in red for identification. **(B)** 24-hour growth curves of *P. aeruginosa* Δ*nahK* and dual genomic deletion of signal transduction proteins involved in the Gac MKN where the secondary gene was deleted from the *P. aeruginosa* Δ*nahK*. *P. aeruginosa* UCBPP-PA14 Δ*nahK* colored in red for identification. **(C)** Growth curves of *P. aeruginosa* wildtype and single Gac MKN non-kinase markerless genomic deletions over 24 hours. *P. aeruginosa* UCBPP-PA14 wildtype colored in red for identification.

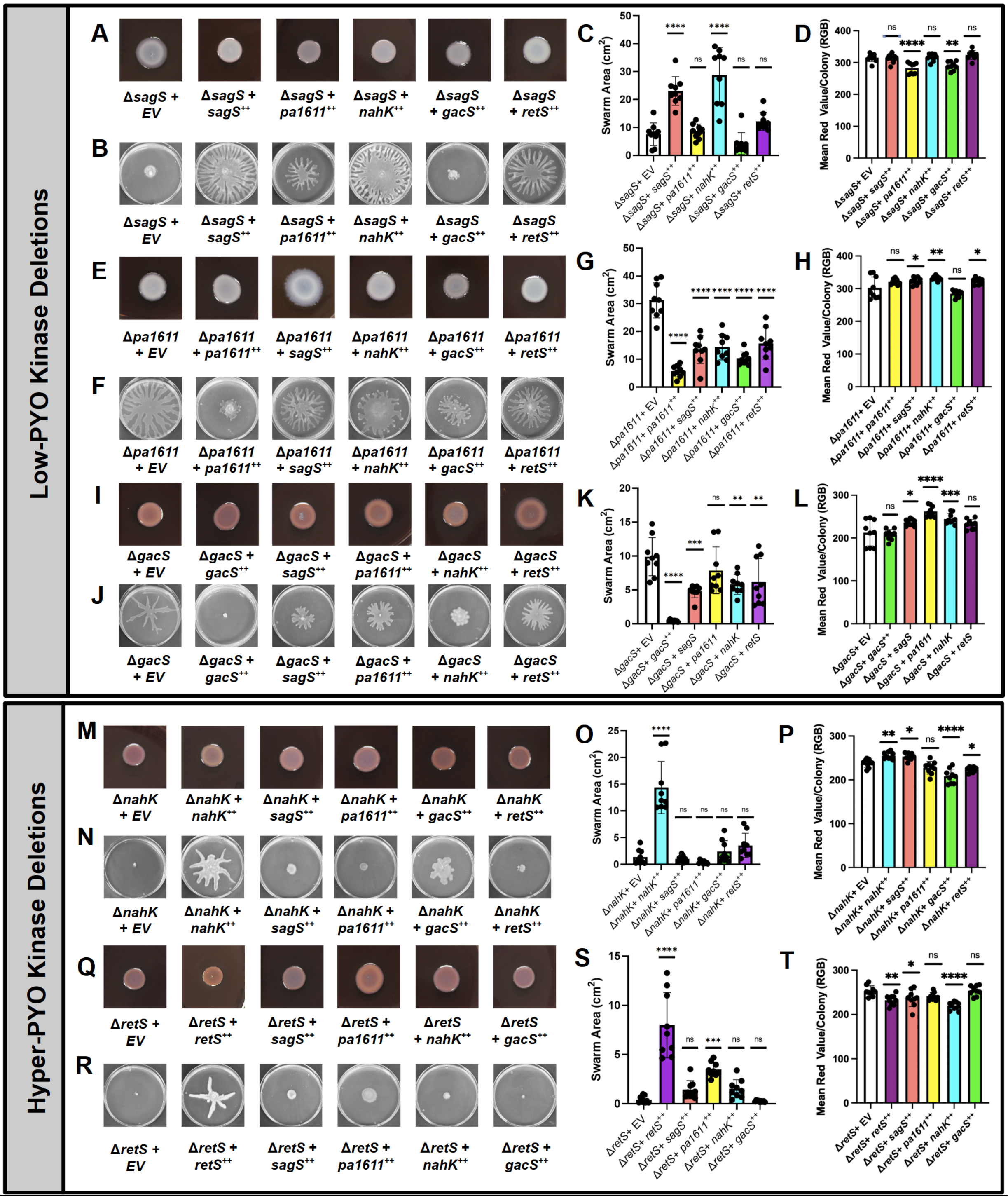


**Supplementary Figure S4: Gac MKN heterodimerization tailors RsmA activity to control planktonic-to-biofilm switch: (A-L)** Qualitative and quantitative phenotypic screening of EPS secretion and swarming motility of PA14 low-PYO producing kinase single deletion strains electroporated with pUCP22 vectors containing either empty vector, *sagS*, *pa1611*, *nahK*, *gacS*, or *retS*. **(M-T)** Qualitative and quantitative phenotypic screening of EPS secretion and swarming motility of PA14 hyper-PYO producing kinase single deletion strains electroporated with pUCP22 vectors containing either empty vector, *sagS*, *pa1611*, *nahK, gacS*, or *retS*. **(A,E,I,M,Q)** Representative images of PA14 static Congo Red colony biofilms. (n = 9). **(B,F,J,N,R)** Representative images of PA14 swarming motility on low-density agar (n=9). **(C,G,K,O,S)** Swarming area ($cm^2$) of PA14 strains electroporated with pUCP22 vectors containing one of the following: an empty vector (white bars), *sagS* (coral bars), *pa1611* (yellow bars), *nahK* (cyan bars), *gacS* (green bars), or *retS* (purple bars) cultured on low-density agar media. (n=9). **(D,H,LP,T)** RGB quantification for EPS staining of PA14 colony biofilms electroporated with pUCP22 vectors containing one of the following: an empty vector (white bars), *sagS* (coral bars), *pa1611* (yellow bars), *nahK* (cyan bars), *gacS* (green bars), or *retS* (purple bars) cultured on 1% tryptone, 1% BactoAgar media supplemented with 40 µg/mL Congo Red dye and 20 µg/mL Brilliant Blue dye. (n = 9). Error bars represent standard deviation from the mean. Significant differences are indicated with asterisks (N.S. $P > 0.05$; *$P < 0.05$; **$P < 0.01$; ***$P < 0.001$; ****$P < 0.0001$; one-way ANOVA and a Tukey multiple comparisons test).
