## Supplementary Table 4 for "Pseudomonas aeruginosa uses kinases NahK and RetS to control the motile-biofilm switch"

| Locus Tag | Gene | *nahK* vs wildtype ($log_2$FC ) | *rsmA* vs wildtype ($log_2$FC ) |
|---|---|---|---|
| PA14_00030 | recF | o | -0.52373 |
| PA14_00080 | | | 1.65094 |
| PA14_00130 | | -0.65098 | |
| PA14_00170 | trkA | 0.41803 | 0.76812 |
| PA14_00180 | rsmB | 0.82565 | 1.02924 |
| PA14_00190 | fmt | 0.58881 | 0.88011 |
| PA14_00200 | def | -0.61284 | |
| PA14_00250 | qor | | |
| PA14_00300 | | 1.23391 | 0.82851 |
| PA14_00310 | | 0.9861 | |
| PA14_00440 | trpA | 0.3989 | 1.52734 |
| PA14_00450 | trpB | | 1.11988 |
| PA14_00490 | tpsB2 | -0.80233 | 0.64609 |
| PA14_00510 | tpsA2 | -1.50677 | 1.02412 |
| PA14_00530 | | | 0.59495 |
| PA14_00560 | exoT | 0.71535 | -0.57097 |
| PA14_00570 | | | 1.50577 |
| PA14_00580 | | | 0.93585 |
| PA14_00590 | | | 0.98993 |
| PA14_00640 | phzH | 1.28843 | 1.83595 |
| PA14_00650 | | 1.11845 | 1.30886 |
| PA14_00660 | | | 1.45488 |
| PA14_00760 | | | |
| PA14_00770 | | -0.41348 | |
| PA14_00820 | tagQ | -0.6714 | 0.63877 |
| PA14_00830 | tagR | | 0.70701 |
| PA14_00850 | tagS | | 0.83511 |
| PA14_00860 | tagT | | 0.55092 |
| PA14_00875 | ppkA | | 1.01828 |
| PA14_00890 | pppA | | 0.9316 |
| PA14_00900 | tagF | -0.53755 | 0.71734 |
| PA14_00910 | tssM | -0.50024 | 0.72718 |
| PA14_00925 | tssL1 | -0.46305 | 1.04329 |
| PA14_00940 | tssK | | 1.24508 |
| PA14_00960 | tssJ | -0.52435 | 1.13063 |
| PA14_00990 | tssA | -0.42764 | |
| PA14_01010 | tssB | -0.57966 | 0.99873 |
| PA14_01020 | tssC | -0.56613 | 1.1743 |
| PA14_01030 | hcp1 | -1.4389 | 1.44988 |
| PA14_01040 | tagJ | -0.61424 | 1.2983 |
| PA14_01060 | tssE | -0.83852 | 1.09911 |
| PA14_01070 | tssF | -0.58041 | 1.81461 |
| PA14_01080 | tssG | | 1.32371 |
| PA14_01100 | tssH | -0.95979 | 1.18523 |
| PA14_01110 | vgrG1a | -1.11981 | 1.21193 |
| PA14_01120 | tsi6 | | 0.94767 |

| | | | |
|---|---|---|---|
| PA14_01130 | tis1 | 5.50634 | 7.39168 |
| PA14_01140 | tas1 | | 2.97134 |
| PA14_01150 | eagT6 | | 1.18401 |
| PA14_01160 | vgrG1b | | 1.18506 |
| PA14_01170 | | -0.62184 | 1.8012 |
| PA14_01180 | | -0.65535 | 2.20803 |
| PA14_01190 | | | 2.38392 |
| PA14_01200 | | 0.49602 | 4.47985 |
| PA14_01220 | | 5.26477 | 8.09802 |
| PA14_01230 | | -1.1073 | 1.15544 |
| PA14_01250 | | 1.43879 | 1.38519 |
| PA14_01380 | cyoE | | 0.78343 |
| PA14_01430 | | | 0.6424 |
| PA14_01460 | | | -1.1303 |
| PA14_01480 | | -0.54668 | |
| PA14_01490 | | 0.66242 | 2.34025 |
| PA14_01510 | parE | -0.76927 | |
| PA14_01660 | guaD | | -0.51144 |
| PA14_01680 | | 0.53068 | |
| PA14_01720 | ahpF | 0.74096 | |
| PA14_01730 | ppk2 | 0.53817 | 0.91314 |
| PA14_01760 | nuh | | |
| PA14_01770 | | 4.51453 | 3.97608 |
| PA14_01780 | | 1.31106 | 0.62995 |
| PA14_01810 | | | 0.61543 |
| PA14_01840 | | | -1.28191 |
| PA14_01860 | | | -1.23908 |
| PA14_01960 | triB | | 0.46487 |
| PA14_02050 | | | -0.94308 |
| PA14_02070 | | 0.53331 | 0.62039 |
| PA14_02090 | | | 0.72022 |
| PA14_02150 | siaA | 0.93718 | 0.99072 |
| PA14_02180 | cheB | | |
| PA14_02190 | cheD | | |
| PA14_02200 | | | |
| PA14_02230 | cheW | | |
| PA14_02310 | atsA | | |
| PA14_02360 | | -1.89755 | -1.72882 |
| PA14_02390 | | -2.769 | -2.45389 |
| PA14_02410 | | -1.28837 | -1.16889 |
| PA14_02435 | | -0.62157 | |
| PA14_02490 | tonB2 | -2.54325 | -1.56158 |
| PA14_02500 | exbB1 | -2.00824 | |
| PA14_02510 | | -1.59212 | |
| PA14_02520 | | | |
| PA14_02530 | | -5.4889 | -5.38342 |
| PA14_02910 | | 1.14157 | 1.09028 |
| PA14_03100 | | 3.34357 | 2.34249 |

| | | | |
|---|---|---|---|
| PA14_03130 | hudA | | 0.59333 |
| PA14_03163 | | 8.2443 | 8.4527 |
| PA14_03166 | | 5.09239 | 4.58927 |
| PA14_03170 | | -0.4338 | 3.54164 |
| PA14_03190 | | -0.66877 | 3.69816 |
| PA14_03200 | tle3 | -0.63588 | 3.14756 |
| PA14_03210 | | | 8.00466 |
| PA14_03220 | vgrG2b | 5.58909 | 4.23763 |
| PA14_03240 | hcpC | 4.68834 | 2.93657 |
| PA14_03250 | | 4.04679 | 3.28821 |
| PA14_03265 | | 5.70501 | 7.12961 |
| PA14_03270 | | 4.67624 | 3.92116 |
| PA14_03285 | | 2.65399 | 2.54708 |
| PA14_03290 | | 3.74436 | 3.66532 |
| PA14_03300 | | 4.95216 | |
| PA14_03310 | | 4.79201 | 4.09698 |
| PA14_03320 | | 4.382 | 5.31221 |
| PA14_03330 | | 6.78543 | 7.03253 |
| PA14_03340 | | 5.46023 | 4.16731 |
| PA14_03350 | | 6.11299 | 5.67935 |
| PA14_03360 | | 5.39471 | 3.07723 |
| PA14_03370 | | 7.32409 | 7.87223 |
| PA14_03380 | | 1.3015 | |
| PA14_03510 | | | 0.50482 |
| PA14_03560 | | 0.49813 | |
| PA14_03670 | cysW | -0.80283 | -1.25911 |
| PA14_03680 | cysT | -0.82234 | -1.24626 |
| PA14_03700 | | -2.21831 | -2.37308 |
| PA14_03710 | oscA | -3.71829 | -3.83343 |
| PA14_03730 | | | -0.75885 |
| PA14_03800 | oprE | -1.3259 | |
| PA14_04010 | | | 0.54659 |
| PA14_04040 | | 0.4332 | |
| PA14_04070 | | -0.61733 | -0.56405 |
| PA14_04080 | | 0.46107 | |
| PA14_04100 | | | 0.83579 |
| PA14_04110 | serA | -1.07398 | -1.1832 |
| PA14_04140 | | | -0.46599 |
| PA14_04180 | | | 0.8979 |
| PA14_04290 | | 0.86195 | 0.94343 |
| PA14_04300 | | -0.78268 | |
| PA14_04310 | rpiA | | -0.57407 |
| PA14_04340 | | | 0.8034 |
| PA14_04350 | | | 1.47048 |
| PA14_04380 | | | -0.54173 |
| PA14_04410 | ptsP | -0.39613 | |
| PA14_04420 | | | |
| PA14_04440 | | | -0.58222 |

| | | | |
|---|---|---|---|
| PA14_04480 | | | -0.43277 |
| PA14_04510 | | | 0.59562 |
| PA14_04520 | | 0.45387 | |
| PA14_04530 | | 0.62308 | |
| PA14_04560 | | 0.46932 | |
| PA14_04590 | | -0.37696 | -0.52752 |
| PA14_04610 | | -0.77 | -0.93724 |
| PA14_04700 | | | 0.85398 |
| PA14_04710 | | 5.60827 | 5.22222 |
| PA14_04750 | | -0.7556 | |
| PA14_04780 | | 2.0837 | 1.8369 |
| PA14_04790 | | 2.57254 | 2.54674 |
| PA14_04810 | laoC | 2.0127 | 1.85546 |
| PA14_04820 | laoR | 0.96057 | 1.01055 |
| PA14_04830 | | -0.51953 | |
| PA14_04900 | rpoH | | -0.43437 |
| PA14_04940 | | 0.94691 | |
| PA14_04950 | mtgA | | -0.63804 |
| PA14_04970 | thiS | -0.61956 | -0.73718 |
| PA14_04980 | | | -0.4805 |
| PA14_05000 | trmB | | |
| PA14_05070 | metW | -0.89531 | -0.84184 |
| PA14_05080 | metX | -1.15954 | -1.35032 |
| PA14_05300 | | 0.54812 | 0.52862 |
| PA14_05440 | | | 1.06368 |
| PA14_05500 | | 0.47801 | |
| PA14_05530 | mexA | 0.59179 | 0.6682 |
| PA14_05560 | | 0.7892 | 0.44547 |
| PA14_05590 | metF | | |
| PA14_05600 | | -0.88368 | |
| PA14_05620 | sahH | -0.80795 | |
| PA14_05630 | | 0.94369 | |
| PA14_05640 | | | -0.59199 |
| PA14_05650 | | | -0.6311 |
| PA14_05820 | | 0.9523 | 1.65904 |
| PA14_05840 | | | 0.63684 |
| PA14_05890 | | | -0.70228 |
| PA14_05960 | | -0.45309 | -0.86817 |
| PA14_05970 | | 0.76958 | |
| PA14_06000 | clpG | 0.68633 | 1.2229 |
| PA14_06160 | | -0.40921 | |
| PA14_06170 | fiuA | | |
| PA14_06180 | | | -0.65935 |
| PA14_06190 | | | -0.82708 |
| PA14_06200 | | -0.77747 | -0.81463 |
| PA14_06210 | | -0.69539 | |
| PA14_06240 | | -1.56397 | -2.36769 |
| PA14_06270 | | 1.60635 | 1.85658 |

| | | | |
|---|---|---|---|
| PA14_06280 | pcaD | 2.42762 | 3.2279 |
| PA14_06290 | glcB | | 0.70329 |
| PA14_06330 | rarD | | -0.43224 |
| PA14_06340 | | | -0.46174 |
| PA14_06420 | | -1.0679 | -1.14096 |
| PA14_06510 | bioF | | 0.47431 |
| PA14_06540 | bioC | | 0.52281 |
| PA14_06810 | norC | | |
| PA14_06870 | dnr | | |
| PA14_06880 | | -0.49955 | -0.96079 |
| PA14_06890 | | | -0.77598 |
| PA14_06900 | | -0.84769 | -1.70063 |
| PA14_06920 | | -0.76169 | -1.36644 |
| PA14_07090 | metK | | |
| PA14_07240 | | 0.58836 | 0.51725 |
| PA14_07250 | | -1.36586 | -1.41708 |
| PA14_07270 | | | -0.59771 |
| PA14_07300 | | -1.977 | |
| PA14_07340 | | -0.45434 | -0.50909 |
| PA14_07360 | | | |
| PA14_07400 | | | |
| PA14_07420 | impA | 1.18835 | 2.27394 |
| PA14_07480 | ltrA | 7.93869 | 8.1434 |
| PA14_07500 | rmcA | 0.45448 | 0.77413 |
| PA14_07580 | plsY | | -0.39144 |
| PA14_07600 | folK | 0.59061 | |
| PA14_07650 | | | 0.53513 |
| PA14_07660 | | | 0.48452 |
| PA14_07680 | | | 0.66441 |
| PA14_07800 | | | -0.60716 |
| PA14_07840 | | | -0.44172 |
| PA14_07940 | trpE | 0.38399 | |
| PA14_07950 | prtN | | |
| PA14_07980 | | 1.3745 | 2.64967 |
| PA14_07990 | | | 0.99571 |
| PA14_08040 | | -0.49382 | |
| PA14_08140 | | | |
| PA14_08150 | | | 0.75022 |
| PA14_08160 | | | 0.81451 |
| PA14_08180 | | | 0.57526 |
| PA14_08190 | | | 0.67489 |
| PA14_08210 | | | 0.64641 |
| PA14_08220 | | | |
| PA14_08280 | | 0.81422 | 1.14085 |
| PA14_08370 | vfr | | -0.48651 |
| PA14_08380 | | | -0.60272 |
| PA14_08420 | | | |
| PA14_08450 | | | 0.44823 |

| | | | |
|---|---|---|---|
| PA14_08510 | erpA | | -0.62036 |
| PA14_08570 | | -2.13999 | |
| PA14_08600 | | -1.39005 | |
| PA14_08680 | tufB | -1.21516 | -1.29555 |
| PA14_08695 | secE | | -1.01444 |
| PA14_08710 | nusG | | -0.62196 |
| PA14_08810 | rpsG | -0.39997 | -0.45072 |
| PA14_08840 | rpsJ | -1.20215 | -1.278 |
| PA14_08850 | rplC | -0.95793 | -1.16068 |
| PA14_08860 | rplD | -0.95492 | -1.03128 |
| PA14_08870 | rplW | -1.01465 | -1.05916 |
| PA14_08880 | rplB | -0.48926 | |
| PA14_08890 | rpsS | -1.2197 | -1.1964 |
| PA14_08900 | rplV | -0.9287 | -0.88253 |
| PA14_08910 | rpsC | -0.85906 | -0.71501 |
| PA14_08920 | rplP | -0.96517 | -0.76677 |
| PA14_08930 | rpmC | -1.08631 | -1.52805 |
| PA14_08940 | rpsQ | -1.07869 | -1.0464 |
| PA14_08950 | rplN | -0.76458 | -0.61465 |
| PA14_08960 | rplX | -0.90278 | -0.79012 |
| PA14_08970 | rplE | -0.77824 | -0.74113 |
| PA14_08980 | rpsN | -0.46293 | -0.61821 |
| PA14_08990 | rpsH | | -0.58099 |
| PA14_09000 | rplF | | -0.41483 |
| PA14_09010 | rplR | | -0.48429 |
| PA14_09070 | rpmJ | -0.81625 | -1.16007 |
| PA14_09180 | uvrA | | -0.4123 |
| PA14_09350 | | -1.34476 | -1.23952 |
| PA14_09400 | phzS | 0.82919 | 5.05256 |
| PA14_09410 | phzG1 | -0.67484 | 2.46479 |
| PA14_09420 | phzF1 | 3.32201 | 4.8859 |
| PA14_09440 | phzE1 | 3.14773 | 4.83613 |
| PA14_09450 | phzD1 | 3.39466 | 5.26464 |
| PA14_09460 | phzC1 | -1.31995 | 1.60378 |
| PA14_09470 | phzB1 | 1.7085 | 4.2728 |
| PA14_09490 | phzM | 1.69944 | 2.03745 |
| PA14_09500 | opmD | 3.34115 | 4.84928 |
| PA14_09520 | mexI | 2.81882 | 4.27816 |
| PA14_09530 | mexH | 3.04176 | 4.76876 |
| PA14_09540 | mexG | 2.67426 | 4.4412 |
| PA14_09580 | nmoA | -0.50827 | |
| PA14_09700 | pqsL | | 0.78251 |
| PA14_09760 | | | -0.56563 |
| PA14_09770 | souR | 0.77383 | 0.57415 |
| PA14_09810 | | 0.52952 | |
| PA14_09880 | | 1.69113 | 0.83304 |
| PA14_09900 | prpL | 2.24617 | 2.51155 |
| PA14_09920 | | 0.75487 | |

| | | | |
|---|---|---|---|
| PA14_09950 | | | -0.82054 |
| PA14_09960 | | | -1.13306 |
| PA14_10050 | | 1.12041 | |
| PA14_10070 | | 3.05993 | 3.78453 |
| PA14_10090 | | 5.23474 | 5.08971 |
| PA14_10120 | | 5.75571 | 3.49919 |
| PA14_10140 | fepG | | -1.22542 |
| PA14_10160 | fepD | | -1.18389 |
| PA14_10170 | fepB | | -0.97398 |
| PA14_10180 | fepC | | -1.02876 |
| PA14_10190 | | -0.57522 | -0.8522 |
| PA14_10200 | | -0.50922 | -1.42752 |
| PA14_10210 | | -0.85592 | -2.52061 |
| PA14_10260 | | -1.02885 | -1.3516 |
| PA14_10330 | | 0.81242 | 0.92683 |
| PA14_10340 | | 3.36126 | 3.64104 |
| PA14_10350 | | 3.68716 | 3.71356 |
| PA14_10360 | | 3.80579 | 3.77453 |
| PA14_10370 | | 2.84924 | 3.75667 |
| PA14_10380 | | 6.14099 | 2.54977 |
| PA14_10400 | | 5.85287 | 5.01235 |
| PA14_10420 | tyrS | | 0.7507 |
| PA14_10530 | mapR | | 2.23379 |
| PA14_10540 | ccoG | 1.12665 | 3.4029 |
| PA14_10550 | | 0.98678 | 3.72941 |
| PA14_10560 | | 0.50235 | 3.61023 |
| PA14_10570 | hpaI | | 0.91991 |
| PA14_10680 | | 0.46762 | |
| PA14_10700 | bphP | 1.17977 | |
| PA14_10740 | | -0.60797 | |
| PA14_10770 | | | 0.56398 |
| PA14_10790 | | | 0.53547 |
| PA14_10830 | | 7.30851 | 7.25536 |
| PA14_10840 | | 5.5792 | 2.83108 |
| PA14_10910 | | 2.44006 | 2.40504 |
| PA14_10960 | | 3.06615 | 3.2765 |
| PA14_10970 | | 1.54616 | 1.48021 |
| PA14_10980 | | 5.15333 | 5.11485 |
| PA14_11120 | | 0.9461 | |
| PA14_11130 | | -0.37857 | |
| PA14_11140 | | 1.04902 | 2.91934 |
| PA14_11150 | | -1.58343 | |
| PA14_11160 | | -1.38701 | -0.65516 |
| PA14_11180 | | -0.46639 | -0.54655 |
| PA14_11190 | | | -0.61799 |
| PA14_11270 | oprG | | 0.69447 |
| PA14_11310 | | 0.46252 | |
| PA14_11380 | nrdR | -0.4833 | -0.56792 |

| | | | |
|---|---|---|---|
| PA14_11430 | ribH | -0.50227 | -0.82842 |
| PA14_11450 | nusB | -0.84495 | -0.98212 |
| PA14_11510 | ribA | -0.86082 | -1.17309 |
| PA14_11520 | | -0.64967 | -0.89331 |
| PA14_11530 | | -0.52724 | -1.07081 |
| PA14_11580 | | | 0.4681 |
| PA14_11590 | | -3.92248 | -3.87731 |
| PA14_11690 | ppa | -0.44887 | |
| PA14_11750 | | -0.79528 | -0.95265 |
| PA14_11760 | eutC | | 0.49959 |
| PA14_11770 | eutB | | 0.75086 |
| PA14_11790 | eat | | |
| PA14_11810 | | 0.76163 | |
| PA14_11910 | | | 0.50683 |
| PA14_11970 | | | -0.5624 |
| PA14_11980 | | -0.42646 | -0.61173 |
| PA14_12010 | proA | | -0.39152 |
| PA14_12140 | | 0.41917 | 0.70726 |
| PA14_12260 | | | 0.49582 |
| PA14_12270 | | | |
| PA14_12360 | | | -0.61591 |
| PA14_12370 | | -0.568 | |
| PA14_12420 | ladS | 2.08419 | 1.82334 |
| PA14_12430 | | -1.12515 | -1.6399 |
| PA14_12440 | | -1.03928 | -1.925 |
| PA14_12450 | | | -0.67159 |
| PA14_12490 | | -0.75827 | -1.29662 |
| PA14_12560 | | -0.51241 | -0.58081 |
| PA14_12740 | | 0.68089 | 1.0097 |
| PA14_12820 | | | -0.63003 |
| PA14_12870 | tesB | | -0.46426 |
| PA14_12900 | | 0.92512 | 1.48711 |
| PA14_12910 | | -8.5098 | -9.18869 |
| PA14_12920 | tauA | -4.39657 | -3.40407 |
| PA14_12940 | | -3.23824 | -0.93879 |
| PA14_12960 | | -2.52334 | |
| PA14_12970 | tauD | -1.55739 | |
| PA14_12990 | betT | | -0.61755 |
| PA14_13000 | | -0.79934 | -1.77692 |
| PA14_13010 | | -3.70377 | -3.82618 |
| PA14_13110 | | 1.54032 | 1.04196 |
| PA14_13130 | | 1.1673 | 0.88378 |
| PA14_13140 | | 0.95703 | 0.80799 |
| PA14_13150 | | | 0.66469 |
| PA14_13170 | copA1 | 0.37413 | 0.66843 |
| PA14_13200 | | 4.77558 | 4.27882 |
| PA14_13210 | | 3.5209 | 4.83488 |
| PA14_13220 | | 6.23835 | 5.96066 |

| | | | |
|---|---|---|---|
| PA14_13230 | moaC | -0.4237 | |
| PA14_13250 | moaE | -0.38439 | -0.55113 |
| PA14_13340 | | 1.00887 | 2.11718 |
| PA14_13350 | tsiT | | 3.2389 |
| PA14_13360 | | 1.59578 | 4.15478 |
| PA14_13370 | | 1.14148 | 3.40391 |
| PA14_13380 | | 1.34846 | 1.99411 |
| PA14_13410 | prfC | -0.85301 | -0.87584 |
| PA14_13450 | | | -0.45108 |
| PA14_13470 | mdrR1 | | |
| PA14_13490 | | | 1.79569 |
| PA14_13500 | | | -0.50004 |
| PA14_13590 | | 1.03241 | |
| PA14_13600 | | 0.91104 | |
| PA14_13610 | | 0.92878 | |
| PA14_13620 | nhaP | 0.96746 | |
| PA14_13630 | | | |
| PA14_13880 | | 1.56909 | 1.32831 |
| PA14_13950 | | | -0.71993 |
| PA14_14040 | rhl | -0.72914 | -0.75874 |
| PA14_14060 | | -0.39645 | -0.46913 |
| PA14_14080 | | -1.58603 | |
| PA14_14170 | | 1.64628 | 1.6475 |
| PA14_14200 | | | -0.77784 |
| PA14_14220 | | -2.62323 | -1.20797 |
| PA14_14230 | | | -1.7213 |
| PA14_14370 | | 2.99736 | 3.23145 |
| PA14_14380 | | -1.03801 | -0.76138 |
| PA14_14480 | | -0.39217 | |
| PA14_14540 | | 2.54741 | 3.47496 |
| PA14_14550 | | | -2.38278 |
| PA14_14590 | queA | 0.43328 | |
| PA14_14660 | | | -0.61071 |
| PA14_14690 | trmJ | | -0.60906 |
| PA14_14710 | iscR | | |
| PA14_14730 | iscS | | -0.59165 |
| PA14_14740 | iscU | | -0.59892 |
| PA14_14750 | iscA | | -0.58461 |
| PA14_14780 | hscA | 0.64804 | |
| PA14_14810 | iscX | | -0.5662 |
| PA14_14860 | | 0.54125 | 0.70786 |
| PA14_14880 | ispG | | 0.49399 |
| PA14_14890 | hisS | | 0.56905 |
| PA14_14900 | | 0.39249 | 0.47138 |
| PA14_14930 | engA | 0.84547 | 0.8949 |
| PA14_15130 | | -0.41386 | |
| PA14_15160 | | -1.93787 | -2.36478 |
| PA14_15290 | | -2.04225 | -1.30222 |

| | | | |
|---|---|---|---|
| PA14_15350 | | 6.76126 | 6.17164 |
| PA14_15360 | | 6.52044 | 5.39804 |
| PA14_15370 | | 7.56452 | 7.45363 |
| PA14_15380 | repA | 3.60055 | 3.17166 |
| PA14_15390 | | 3.73119 | 2.94017 |
| PA14_15400 | repC | 4.96295 | 4.80399 |
| PA14_15430 | | 4.73555 | 4.52067 |
| PA14_15435 | | 6.02751 | 4.62443 |
| PA14_15445 | merE | 5.68023 | 4.48378 |
| PA14_15450 | merD | 4.62449 | 4.44347 |
| PA14_15460 | merA | 6.94548 | 7.19544 |
| PA14_15470 | merP | 6.66569 | 7.00752 |
| PA14_15475 | merT | 3.08055 | 2.67689 |
| PA14_15480 | | 6.78455 | 6.3123 |
| PA14_15510 | traJ | 5.72204 | 3.7805 |
| PA14_15520 | trbJ | 5.55982 | 5.28026 |
| PA14_15540 | trbL | 1.68465 | 1.27321 |
| PA14_15560 | | 4.69272 | 4.48555 |
| PA14_15570 | | 8.39052 | 8.63759 |
| PA14_15580 | | 4.36193 | 4.03352 |
| PA14_15590 | | 6.17744 | 5.11673 |
| PA14_15600 | | 7.22723 | 7.51373 |
| PA14_15610 | | 6.88907 | 7.13434 |
| PA14_15620 | | 7.03455 | 7.40812 |
| PA14_15630 | | 2.01974 | 2.6986 |
| PA14_15650 | | 3.30163 | 2.55109 |
| PA14_15660 | | 8.0006 | 8.15863 |
| PA14_15670 | | 8.54165 | 8.66983 |
| PA14_15680 | | | -0.80374 |
| PA14_15750 | | -0.74567 | -1.106 |
| PA14_15770 | nagE | 1.62349 | |
| PA14_15780 | ptsP | 2.48277 | 1.35791 |
| PA14_15790 | | 2.29117 | 0.83303 |
| PA14_15810 | nagA | 2.67791 | 1.59637 |
| PA14_15820 | | 2.98416 | 2.51681 |
| PA14_15970 | rpsP | -1.13104 | -1.32038 |
| PA14_15980 | rimM | -1.21132 | -1.22157 |
| PA14_15990 | trmD | -0.60098 | -0.78358 |
| PA14_16000 | rplS | -0.50988 | -0.63925 |
| PA14_16100 | | 1.56962 | 2.53924 |
| PA14_16110 | | 1.0352 | 1.79624 |
| PA14_16140 | | -0.67257 | 1.7575 |
| PA14_16150 | | | 1.66886 |
| PA14_16160 | | -0.4509 | 2.65681 |
| PA14_16180 | | -0.79871 | 2.83129 |
| PA14_16190 | | -1.12129 | 2.38669 |
| PA14_16200 | | -1.52536 | 2.77798 |
| PA14_16210 | | | 0.8501 |

| | | | |
|---|---|---|---|
| PA14_16250 | lasB | 2.45384 | 2.47525 |
| PA14_16260 | | 0.59672 | 0.93911 |
| PA14_16270 | | -0.57358 | |
| PA14_16280 | nalC | 1.47418 | 1.93518 |
| PA14_16290 | | | 0.82593 |
| PA14_16310 | | 0.82015 | 0.80767 |
| PA14_16330 | | | |
| PA14_16340 | | | 1.06685 |
| PA14_16430 | wspA | -0.85708 | -0.62275 |
| PA14_16440 | | -0.49274 | -0.425 |
| PA14_16580 | | 0.85256 | 1.01748 |
| PA14_16590 | | 0.43035 | 0.75289 |
| PA14_16600 | | 0.67026 | 0.82935 |
| PA14_16660 | | 1.65383 | 1.97455 |
| PA14_16670 | cadR | -0.5379 | -0.57165 |
| PA14_16730 | | 0.38483 | |
| PA14_16750 | | -1.76606 | -1.64857 |
| PA14_16840 | | | |
| PA14_16950 | dapD | 0.43923 | |
| PA14_16970 | | -0.45615 | -0.82889 |
| PA14_16990 | | | -0.98165 |
| PA14_17000 | | | |
| PA14_17010 | | | -0.57127 |
| PA14_17060 | rpsB | -0.43066 | -0.5978 |
| PA14_17180 | lpxD | | 0.57563 |
| PA14_17230 | rnhB | 0.67002 | 1.01514 |
| PA14_17280 | tilS | 0.49467 | 0.52549 |
| PA14_17340 | ispD | 0.51739 | 0.48253 |
| PA14_17350 | yedF | 0.65876 | 1.12016 |
| PA14_17400 | adhC | 0.54882 | 0.80001 |
| PA14_17410 | fghA | | 0.65493 |
| PA14_17480 | rpoS | | 0.71589 |
| PA14_17490 | fdxA | -0.57427 | |
| PA14_17500 | mutS | -0.89155 | -0.83853 |
| PA14_17510 | | | 1.85051 |
| PA14_17530 | recA | -0.42675 | -0.68503 |
| PA14_17550 | | | 0.43025 |
| PA14_17570 | | | 0.51612 |
| PA14_17580 | | 0.79114 | 1.20422 |
| PA14_17660 | | | -0.50685 |
| PA14_17700 | rpmE2 | -0.64784 | -0.62867 |
| PA14_17710 | rpmJ2 | -0.89168 | -1.37927 |
| PA14_17720 | | | -0.46023 |
| PA14_17730 | | | |
| PA14_17740 | | 0.5784 | |
| PA14_17810 | | | -0.7361 |
| PA14_17850 | | | -0.70209 |
| PA14_17900 | metR | | -0.63448 |

| | | | |
|---|---|---|---|
| PA14_17960 | glpK | 0.96542 | |
| PA14_17980 | glpF | 1.0044 | 0.80529 |
| PA14_18040 | | 0.35064 | 0.46589 |
| PA14_18060 | | -1.80366 | -1.13672 |
| PA14_18100 | mmsR | | 0.55472 |
| PA14_18110 | mmsA | | -0.72943 |
| PA14_18180 | | | -0.50486 |
| PA14_18250 | ptsP | 0.43242 | |
| PA14_18320 | | | -0.47563 |
| PA14_18330 | arnT | -0.42564 | -1.02936 |
| PA14_18350 | arnA | -0.39975 | -0.63334 |
| PA14_18360 | arnC | -0.45521 | -0.71449 |
| PA14_18370 | arnB | -0.47603 | -1.20888 |
| PA14_18510 | algE | -0.51414 | |
| PA14_18580 | algD | | |
| PA14_18630 | | 2.32053 | 1.14269 |
| PA14_18650 | grxD | | -0.59024 |
| PA14_18660 | | 0.72581 | |
| PA14_18680 | | -0.75549 | -0.6015 |
| PA14_18720 | motY | -1.19388 | -1.00135 |
| PA14_18740 | argG | -0.85459 | -0.94558 |
| PA14_18750 | gloA1 | | -0.6671 |
| PA14_18800 | | 0.59054 | 1.12626 |
| PA14_18810 | | 2.25637 | 1.60487 |
| PA14_18820 | | | |
| PA14_18830 | purB | | |
| PA14_18880 | nth | | 0.8913 |
| PA14_18910 | | | 0.6465 |
| PA14_18920 | rsxC | | 0.43679 |
| PA14_18960 | tli5a | | 5.10761 |
| PA14_19010 | tsi3 | -1.03702 | 0.5687 |
| PA14_19020 | tse3 | -0.71194 | 2.02797 |
| PA14_19030 | | | 0.43186 |
| PA14_19140 | pheC | | 0.6156 |
| PA14_19170 | | | -1.04831 |
| PA14_19210 | | | -0.53248 |
| PA14_19230 | | | -0.74518 |
| PA14_19310 | | | 0.64413 |
| PA14_19320 | | 1.02181 | 0.58926 |
| PA14_19340 | | 0.54185 | |
| PA14_19350 | | 1.00533 | 0.97145 |
| PA14_19360 | ngg | 0.82511 | 0.93324 |
| PA14_19370 | | 1.0252 | 0.8821 |
| PA14_19380 | | -0.35231 | |
| PA14_19390 | | 0.83698 | 0.56046 |
| PA14_19450 | | -0.42376 | |
| PA14_19470 | mqoA | | |
| PA14_19480 | | -0.66411 | -0.69016 |

| | | | |
|---|---|---|---|
| PA14_19490 | | -5.20862 | -5.03897 |
| PA14_19500 | | -1.8251 | -0.80632 |
| PA14_19510 | | -0.74767 | |
| PA14_19520 | | | |
| PA14_19530 | ssuE | -4.72489 | -4.44338 |
| PA14_19540 | | -4.25168 | -3.29771 |
| PA14_19560 | ssuD | -3.27151 | -1.60409 |
| PA14_19570 | ssuC | -2.22744 | -0.81986 |
| PA14_19580 | ssuB | -1.9898 | |
| PA14_19680 | | | -0.92537 |
| PA14_19690 | | | -1.98528 |
| PA14_19700 | | | -0.84377 |
| PA14_19710 | | | |
| PA14_19810 | | -0.79091 | -1.15972 |
| PA14_19860 | | 0.59883 | |
| PA14_19920 | | | -0.60654 |
| PA14_19990 | | | -0.52143 |
| PA14_20010 | hasR | | 0.47595 |
| PA14_20120 | | | 0.59383 |
| PA14_20130 | | -0.45115 | |
| PA14_20140 | fpr | -0.41499 | |
| PA14_20170 | nosY | | |
| PA14_20200 | nosZ | | |
| PA14_20230 | nosR | | |
| PA14_20290 | amrZ | -0.93035 | -0.8113 |
| PA14_20300 | phnC | 0.55469 | 0.60602 |
| PA14_20400 | phnK | 0.588 | 0.81172 |
| PA14_20480 | | 0.96514 | 0.70912 |
| PA14_20510 | | 2.11096 | 2.22315 |
| PA14_20520 | | 2.37946 | 3.41537 |
| PA14_20530 | | 0.89086 | 1.48233 |
| PA14_20550 | | -0.75208 | |
| PA14_20600 | | 6.05341 | 4.51894 |
| PA14_20610 | lecB | | 3.56182 |
| PA14_20700 | flgZ | -0.85821 | -0.72871 |
| PA14_20730 | flgM | -0.91708 | -0.69244 |
| PA14_20760 | cheR | | -0.4287 |
| PA14_20770 | hsbA | 0.73529 | 0.58073 |
| PA14_20780 | hsbR | 0.91098 | 0.64022 |
| PA14_20820 | | -0.39821 | -0.57639 |
| PA14_20840 | | 0.6197 | 0.60616 |
| PA14_20850 | | | -0.48501 |
| PA14_20890 | rfaD | | 0.75248 |
| PA14_20900 | | -1.08912 | 0.99245 |
| PA14_20920 | | -1.59465 | 0.89964 |
| PA14_20940 | | -2.44225 | |
| PA14_20950 | fabH2 | -1.51466 | 0.92453 |
| PA14_20960 | | -2.10265 | 0.65577 |

| | | | |
|---|---|---|---|
| PA14_20970 | cyp23 | -1.91729 | 0.94104 |
| PA14_20980 | | -2.00837 | 0.70394 |
| PA14_21000 | | -1.82658 | 0.97002 |
| PA14_21010 | | -1.98985 | 0.8874 |
| PA14_21020 | | -1.70993 | 1.12215 |
| PA14_21030 | | | |
| PA14_21070 | | | 0.54549 |
| PA14_21120 | | 2.6962 | 5.2386 |
| PA14_21130 | | | 0.46841 |
| PA14_21140 | phnE | -3.35121 | -3.33212 |
| PA14_21150 | | -3.51268 | -3.28701 |
| PA14_21175 | | | 0.79228 |
| PA14_21240 | | -1.11043 | -1.45127 |
| PA14_21260 | | -0.37779 | -0.95188 |
| PA14_21290 | | | -0.4585 |
| PA14_21340 | fadD2 | 1.00585 | 0.65864 |
| PA14_21370 | fadD1 | | 0.42702 |
| PA14_21450 | tssI | 0.68561 | 4.94091 |
| PA14_21490 | | | 2.6207 |
| PA14_21520 | | | |
| PA14_21600 | | -0.80838 | -1.4752 |
| PA14_21620 | oprP | 0.78126 | 0.71231 |
| PA14_21630 | | | |
| PA14_21750 | | | |
| PA14_21760 | (capB) | | -0.82373 |
| PA14_21780 | | 0.8973 | |
| PA14_21790 | rdgC | | -0.85268 |
| PA14_21820 | | | |
| PA14_21830 | | | -0.6531 |
| PA14_21870 | | 0.6141 | 0.57721 |
| PA14_21940 | | 0.95644 | 1.35558 |
| PA14_21960 | | 0.43284 | 1.02303 |
| PA14_21970 | | 0.82352 | |
| PA14_21980 | | -0.47435 | -0.56484 |
| PA14_22000 | rluA | 0.98046 | 1.148 |
| PA14_22050 | htrB | | |
| PA14_22075 | | 3.20515 | 3.17474 |
| PA14_22080 | | 6.72296 | 6.48811 |
| PA14_22090 | | 2.37876 | 1.93928 |
| PA14_22120 | | 1.00153 | |
| PA14_22190 | | 3.9248 | 3.84193 |
| PA14_22270 | | 5.87314 | 3.42887 |
| PA14_22310 | | 0.52718 | |
| PA14_22370 | | 1.15077 | 0.99685 |
| PA14_22420 | | | 3.34499 |
| PA14_22450 | ppiA | | 0.42633 |
| PA14_22500 | | 2.16314 | 1.40701 |
| PA14_22510 | | 2.35756 | 2.26605 |

| | | | |
|---|---|---|---|
| PA14_22520 | | 3.54687 | 3.42905 |
| PA14_22530 | | 3.01342 | 3.59388 |
| PA14_22540 | | 1.07006 | 1.21842 |
| PA14_22550 | | 5.58079 | 3.0069 |
| PA14_22600 | | 2.96529 | 2.17586 |
| PA14_22650 | | | 0.63327 |
| PA14_22660 | | | 0.50871 |
| PA14_22670 | | | 0.48082 |
| PA14_22690 | trkH | 0.52086 | |
| PA14_22730 | cpxS | | |
| PA14_22770 | | -1.30107 | -1.06445 |
| PA14_22780 | yciL | | -0.50387 |
| PA14_22800 | yciB | -0.45118 | -0.80051 |
| PA14_22890 | gap | | 0.6155 |
| PA14_22990 | | 1.28694 | |
| PA14_23000 | | 1.54195 | 0.76231 |
| PA14_23010 | gltK | 0.97337 | 0.70689 |
| PA14_23090 | | -0.41868 | |
| PA14_23120 | | | -0.48519 |
| PA14_23130 | | | -0.56575 |
| PA14_23240 | | | -0.49198 |
| PA14_23260 | gyrA | | -0.39602 |
| PA14_23290 | hisC2 | -0.37916 | -0.57427 |
| PA14_23330 | rpsA | -0.44514 | -0.96261 |
| PA14_23350 | orfA | 7.81084 | 8.20094 |
| PA14_23360 | wzz | 4.75714 | 4.55344 |
| PA14_23370 | orfK | 6.24751 | 5.89033 |
| PA14_23380 | orfH | 5.36721 | 3.95632 |
| PA14_23390 | orfE | 2.80696 | 2.45641 |
| PA14_23400 | | 2.81462 | 2.65042 |
| PA14_23410 | | 2.36716 | 2.54043 |
| PA14_23420 | zbdP | 5.49207 | 4.72993 |
| PA14_23430 | hepP | 5.95238 | 5.4332 |
| PA14_23440 | | 4.46485 | 4.88468 |
| PA14_23450 | | 4.96195 | 5.15365 |
| PA14_23460 | orfN | 5.02991 | 4.85895 |
| PA14_23470 | | -0.65681 | -0.76667 |
| PA14_23510 | uvrB | -0.77312 | -0.84046 |
| PA14_23640 | | 0.51884 | |
| PA14_23680 | ibpA | | 1.12933 |
| PA14_23700 | | -0.38985 | |
| PA14_23750 | leuC | -0.74382 | -0.61515 |
| PA14_23840 | truA | 0.56114 | 0.76052 |
| PA14_23850 | trpF | | -0.68287 |
| PA14_23920 | purF | | -0.4717 |
| PA14_23970 | xcpQ | 0.82218 | 0.8565 |
| PA14_23980 | xcpP | 0.48051 | |
| PA14_23990 | xcpR | 1.0996 | 0.61388 |

| | | | |
|---|---|---|---|
| PA14_24010 | xcpS | 0.81156 | |
| PA14_24020 | xcpT | 0.85985 | 0.75949 |
| PA14_24040 | xcpU | 1.43495 | 0.9258 |
| PA14_24050 | xcpV | 1.68438 | 1.20724 |
| PA14_24060 | xcpW | 1.08917 | |
| PA14_24070 | xcpX | 0.86035 | |
| PA14_24080 | xcpY | 1.19821 | 0.87874 |
| PA14_24100 | xcpZ | 0.52562 | 0.64927 |
| PA14_24190 | | 0.75444 | |
| PA14_24270 | pepN | | 0.38414 |
| PA14_24290 | gbt | 0.65124 | 0.6753 |
| PA14_24300 | | 0.75315 | 0.55479 |
| PA14_24310 | | 0.57271 | -0.50243 |
| PA14_24330 | | 0.72686 | |
| PA14_24360 | | 5.14873 | 2.90141 |
| PA14_24445 | gdhB | 0.36318 | 1.00319 |
| PA14_24480 | pelA | | 1.44415 |
| PA14_24490 | pelB | 0.63445 | 2.00654 |
| PA14_24500 | pelC | | |
| PA14_24510 | pelD | 1.71892 | 3.09581 |
| PA14_24530 | pelE | 1.0534 | 2.958 |
| PA14_24550 | pelF | | 1.84956 |
| PA14_24560 | pelG | | 1.27313 |
| PA14_24600 | | | 1.03765 |
| PA14_24665 | rlmKL | -0.4161 | |
| PA14_24720 | | 0.88215 | 0.92343 |
| PA14_24730 | | 1.62663 | 1.43593 |
| PA14_24740 | | 0.46305 | 0.4907 |
| PA14_24760 | | | 0.49194 |
| PA14_24830 | | -0.50545 | -0.82735 |
| PA14_24850 | | 1.18697 | 1.69363 |
| PA14_24940 | | 1.59821 | 1.31816 |
| PA14_24950 | | 0.47957 | |
| PA14_24980 | | | |
| PA14_24990 | | -0.78465 | 2.37425 |
| PA14_25040 | | -0.6705 | -0.56645 |
| PA14_25160 | lexA | | -0.57607 |
| PA14_25250 | | -0.39393 | -0.46192 |
| PA14_25450 | | 0.42362 | |
| PA14_25520 | | -0.58845 | |
| PA14_25540 | ptpA | -0.40593 | -0.70836 |
| PA14_25560 | rne | | |
| PA14_25590 | | | -0.52377 |
| PA14_25600 | sppA | | -0.65854 |
| PA14_25620 | | | -0.47108 |
| PA14_25630 | rpmF | | -1.10351 |
| PA14_25640 | plsX | | -0.65301 |
| PA14_25670 | acpP | | -0.45337 |

| | | | |
|---|---|---|---|
| PA14_25690 | fabF1 | 0.7819 | 0.62757 |
| PA14_25710 | pabC | 0.58353 | 0.70037 |
| PA14_25920 | cobM | 0.51073 | |
| PA14_25980 | aroF | | 0.57263 |
| PA14_26010 | | -0.61604 | -0.81653 |
| PA14_26020 | paaP | 1.33154 | 3.75219 |
| PA14_26050 | | | -1.20187 |
| PA14_26060 | | -0.85055 | |
| PA14_26150 | | -1.10313 | -1.04066 |
| PA14_26160 | | -0.60713 | -2.61563 |
| PA14_26165 | | | -1.54552 |
| PA14_26190 | | 1.32475 | 1.61649 |
| PA14_26240 | hisJ | | -0.74241 |
| PA14_26310 | | -0.54503 | |
| PA14_26350 | | 0.39051 | 0.76252 |
| PA14_26420 | | | -0.55971 |
| PA14_26450 | mntP | | -0.7656 |
| PA14_26780 | | | -0.56439 |
| PA14_26810 | | | |
| PA14_26850 | | -0.50538 | -0.8263 |
| PA14_26860 | | 0.54709 | 0.47301 |
| PA14_26960 | | -0.36135 | -0.45966 |
| PA14_27000 | | -0.78305 | -0.6515 |
| PA14_27050 | | | -0.496 |
| PA14_27180 | | -1.01983 | -1.11622 |
| PA14_27220 | ohr | | 0.95849 |
| PA14_27230 | ohrR | -0.50299 | |
| PA14_27270 | | -0.91461 | |
| PA14_27390 | | 0.48378 | |
| PA14_27460 | tpm | -0.39046 | |
| PA14_27520 | | 0.62276 | 1.14212 |
| PA14_27530 | ospR | 0.6565 | 1.15072 |
| PA14_27550 | sagS | -0.36419 | |
| PA14_27590 | | -0.46625 | -0.78689 |
| PA14_27630 | rebP2 | 9.01057 | 11.27548 |
| PA14_27640 | rebP1 | 9.28837 | 11.67984 |
| PA14_27650 | | 6.05915 | 8.32756 |
| PA14_27660 | | 2.39161 | 5.0464 |
| PA14_27675 | | 4.52217 | 5.82 |
| PA14_27680 | | 6.18826 | 8.20263 |
| PA14_27690 | | 2.80912 | 6.38262 |
| PA14_27700 | | 2.89756 | 5.06064 |
| PA14_27730 | fadE | | 0.49425 |
| PA14_27740 | | -0.44582 | -0.73436 |
| PA14_27770 | | | 0.48908 |
| PA14_27800 | | | 0.46763 |
| PA14_27960 | tal | 0.74577 | |
| PA14_27980 | dusA | | -0.61653 |

| | | | |
|---|---|---|---|
| PA14_28000 | | | 0.90631 |
| PA14_28010 | | | 1.88871 |
| PA14_28030 | | 0.90423 | 0.81558 |
| PA14_28040 | | -1.17154 | -0.47611 |
| PA14_28070 | | -3.81804 | -4.09805 |
| PA14_28110 | bamI | | 1.08677 |
| PA14_28130 | | -0.98431 | |
| PA14_28150 | | -0.79036 | -1.0715 |
| PA14_28180 | | | -0.71333 |
| PA14_28200 | tsi4 | | 0.81061 |
| PA14_28210 | tse4 | | 2.1331 |
| PA14_28220 | | | 2.98141 |
| PA14_28240 | | | 1.60336 |
| PA14_28250 | | 7.23454 | 7.18145 |
| PA14_28290 | | -0.67904 | -0.6293 |
| PA14_28310 | | -0.76893 | -1.10478 |
| PA14_28320 | | 0.76455 | 0.67122 |
| PA14_28340 | | -0.55249 | -0.51966 |
| PA14_28350 | | 1.20871 | |
| PA14_28380 | | 0.51035 | |
| PA14_28400 | | | |
| PA14_28410 | | | 2.84861 |
| PA14_28440 | | | |
| PA14_28450 | eco | | 1.59961 |
| PA14_28460 | | | 0.52335 |
| PA14_28500 | | | 0.81986 |
| PA14_28520 | | 0.70237 | |
| PA14_28550 | | -1.66188 | -1.50079 |
| PA14_28580 | | | -1.26876 |
| PA14_28590 | map | | -0.94555 |
| PA14_28620 | | 0.46491 | |
| PA14_28660 | infC | -1.13868 | -1.50996 |
| PA14_28670 | rpmI | -1.60165 | -2.12103 |
| PA14_28680 | rplT | -0.77904 | -1.27676 |
| PA14_28720 | ihfA | -0.81483 | -0.82003 |
| PA14_28770 | | 4.30224 | 4.53685 |
| PA14_28800 | | 7.68038 | 7.67468 |
| PA14_28810 | | 4.94065 | 5.16713 |
| PA14_28820 | | 6.35205 | 6.41779 |
| PA14_28830 | | 7.11568 | 7.88391 |
| PA14_28840 | | 5.08699 | 3.46403 |
| PA14_28880 | | | 0.87535 |
| PA14_28895 | | | 0.85593 |
| PA14_28910 | | | 0.74754 |
| PA14_28920 | | | 0.95659 |
| PA14_28930 | | | 1.24926 |
| PA14_28940 | | -1.1362 | |
| PA14_28950 | | -0.77572 | |

| | | | |
|---|---|---|---|
| PA14_28970 | | 7.50634 | 7.20337 |
| PA14_28980 | | 8.65073 | 9.04196 |
| PA14_29000 | | -0.43925 | -0.48715 |
| PA14_29010 | | -0.85727 | -0.91587 |
| PA14_29020 | cpo | | 0.85382 |
| PA14_29030 | | 0.86385 | 1.03497 |
| PA14_29090 | | -2.24064 | -2.33929 |
| PA14_29130 | | -0.87889 | -0.74015 |
| PA14_29200 | tse2 | | 3.39231 |
| PA14_29210 | | -0.41902 | |
| PA14_29220 | | | |
| PA14_29230 | | 1.10302 | 1.99853 |
| PA14_29240 | | 1.01275 | 2.03335 |
| PA14_29250 | | | 0.91652 |
| PA14_29260 | | -0.37812 | |
| PA14_29280 | | -0.55768 | |
| PA14_29330 | | | 4.84396 |
| PA14_29340 | pfeE | -1.46038 | -2.36926 |
| PA14_29350 | pfeA | -2.42265 | -3.09951 |
| PA14_29390 | tssI | 0.50336 | 2.29622 |
| PA14_29400 | tse5 | | 2.37931 |
| PA14_29410 | | | 1.49906 |
| PA14_29420 | | 0.48837 | 0.51254 |
| PA14_29460 | | | 0.74529 |
| PA14_29530 | | | -1.56189 |
| PA14_29590 | mvaU | 0.41679 | 0.88332 |
| PA14_29600 | queD | | -0.57613 |
| PA14_29620 | norR | -0.42583 | -0.44544 |
| PA14_29640 | fhp | -1.00165 | -0.85768 |
| PA14_29710 | | | |
| PA14_29760 | | -2.10433 | -2.11195 |
| PA14_29800 | | -0.39393 | -0.44561 |
| PA14_29850 | nuoN | | 0.46918 |
| PA14_29860 | nuoM | | 0.64867 |
| PA14_30010 | nuoB | | -0.46286 |
| PA14_30070 | | | 0.77121 |
| PA14_30110 | purB | -0.49577 | -0.70288 |
| PA14_30180 | idh | | |
| PA14_30240 | infA | -0.5635 | -0.89989 |
| PA14_30260 | | | |
| PA14_30360 | | | 0.51363 |
| PA14_30410 | | -0.33795 | |
| PA14_30430 | | | 0.89056 |
| PA14_30460 | | -4.54611 | -4.2177 |
| PA14_30470 | | -3.79521 | -3.37652 |
| PA14_30490 | | -3.0203 | -2.07427 |
| PA14_30500 | | -2.1272 | -1.11184 |
| PA14_30520 | | -1.5144 | |

| | | | |
|---|---|---|---|
| PA14_30540 | | -1.66096 | |
| PA14_30550 | | -4.67966 | -4.59483 |
| PA14_30560 | gteE | | 1.06983 |
| PA14_30570 | | | 0.51605 |
| PA14_30620 | | 2.09932 | 1.54007 |
| PA14_30630 | pqsH | 1.23715 | 0.67359 |
| PA14_30650 | gacA | -0.87875 | -0.73517 |
| PA14_30690 | | 2.9396 | 2.78426 |
| PA14_30720 | | | -2.26953 |
| PA14_30730 | | -0.8135 | 0.71568 |
| PA14_30740 | | -0.77661 | -1.41505 |
| PA14_30820 | | | |
| PA14_30840 | | 0.42582 | 0.60634 |
| PA14_30850 | | 2.78542 | 2.66947 |
| PA14_30860 | trbG | 3.36555 | 3.56894 |
| PA14_30900 | trbJ | 4.24852 | 4.33975 |
| PA14_30910 | trbE | | 0.85383 |
| PA14_30940 | trbB | 3.9644 | 3.87029 |
| PA14_30960 | | 0.71757 | |
| PA14_30970 | | 3.99066 | 3.98026 |
| PA14_30980 | tli1 | | 3.76086 |
| PA14_30990 | | 7.21161 | 7.50687 |
| PA14_31000 | | 6.05884 | 3.81567 |
| PA14_31010 | | 5.21072 | 5.03148 |
| PA14_31030 | | 4.67625 | 4.40105 |
| PA14_31040 | | 4.499 | 4.25171 |
| PA14_31050 | | 5.29929 | 4.8062 |
| PA14_31060 | | | 4.94956 |
| PA14_31070 | | 1.82641 | 1.97223 |
| PA14_31100 | | 1.94458 | 2.8302 |
| PA14_31110 | | 0.86689 | |
| PA14_31150 | | 3.71928 | 3.081 |
| PA14_31160 | | 8.49535 | 9.11507 |
| PA14_31170 | | 6.06701 | 3.81043 |
| PA14_31190 | | 1.51387 | 1.06202 |
| PA14_31220 | | 1.35079 | 1.27653 |
| PA14_31230 | | 1.93148 | 1.8794 |
| PA14_31240 | | 6.59524 | 5.50353 |
| PA14_31270 | | 3.68173 | 3.9407 |
| PA14_31280 | | 5.04336 | 2.72559 |
| PA14_31290 | pa1L | 0.61305 | 4.67252 |
| PA14_31330 | | -4.2499 | -4.3461 |
| PA14_31350 | | 1.07652 | 1.84943 |
| PA14_31360 | | 1.49474 | 1.37128 |
| PA14_31380 | | | -0.97896 |
| PA14_31390 | | -2.56539 | -1.99618 |
| PA14_31400 | ctpH | -0.71503 | -2.44587 |
| PA14_31420 | | | 0.6762 |

| | | | |
|---|---|---|---|
| PA14_31440 | | | |
| PA14_31620 | | -0.64037 | -0.65447 |
| PA14_31640 | | -0.74938 | -0.73406 |
| PA14_31700 | | -0.8133 | 0.55019 |
| PA14_31720 | | -2.34849 | 2.24357 |
| PA14_31730 | | -2.22751 | 1.86566 |
| PA14_31740 | | -2.84674 | 0.65813 |
| PA14_31750 | | -0.54487 | 1.70393 |
| PA14_31760 | | -1.48563 | 1.12659 |
| PA14_31770 | | -0.56467 | -0.82026 |
| PA14_31820 | | -1.3463 | 0.6485 |
| PA14_31930 | | -1.59324 | -1.65423 |
| PA14_31960 | | 0.52996 | |
| PA14_31970 | czcC | -1.66295 | -1.4583 |
| PA14_32150 | antB | | |
| PA14_32160 | antA | 0.87638 | |
| PA14_32200 | catR | -1.10094 | -1.47406 |
| PA14_32270 | | 1.89441 | 2.19332 |
| PA14_32280 | | | 0.47938 |
| PA14_32290 | | | 0.83598 |
| PA14_32310 | | | |
| PA14_32330 | | 2.38859 | 2.49712 |
| PA14_32380 | oprN | 1.36475 | 2.31859 |
| PA14_32390 | mexF | 1.92922 | 3.32504 |
| PA14_32400 | mexE | 0.74339 | 2.83193 |
| PA14_32420 | mexS | | 0.52137 |
| PA14_32460 | | 0.44234 | 0.78062 |
| PA14_32470 | | -0.65403 | |
| PA14_32500 | | | -0.51744 |
| PA14_32520 | | -2.58407 | -2.19896 |
| PA14_32530 | | -1.3726 | -1.18125 |
| PA14_32540 | | -2.11608 | -1.75435 |
| PA14_32600 | | | 0.56135 |
| PA14_32630 | | | 1.15634 |
| PA14_32660 | | -0.75303 | -0.78582 |
| PA14_32690 | gtdA | -0.71914 | -0.74575 |
| PA14_32710 | | | -0.61657 |
| PA14_32770 | | | 1.32544 |
| PA14_32780 | tpsB1 | | |
| PA14_32790 | | -2.08855 | |
| PA14_32850 | | 3.70327 | 4.13214 |
| PA14_32860 | | | 1.43127 |
| PA14_32880 | | | 0.58925 |
| PA14_32905 | | | 0.71111 |
| PA14_32930 | | | 0.58615 |
| PA14_32950 | | 1.2424 | 1.97525 |
| PA14_32985 | gcvH2 | -0.90396 | -1.13145 |
| PA14_33000 | gcvP2 | -0.71544 | -1.12299 |

| | | | |
|---|---|---|---|
| PA14_33010 | glyA2 | -0.58779 | -0.92946 |
| PA14_33050 | | 2.07363 | 2.4376 |
| PA14_33060 | | 2.00759 | 1.26358 |
| PA14_33170 | | | 1.15211 |
| PA14_33190 | | -0.45759 | |
| PA14_33220 | | -0.56454 | |
| PA14_33270 | pvdG | -0.44401 | |
| PA14_33280 | pvdL | -0.46827 | |
| PA14_33290 | | 1.10364 | 0.99564 |
| PA14_33300 | cas6f | 7.52255 | 9.21041 |
| PA14_33310 | csy3 | 6.95119 | 8.90516 |
| PA14_33320 | csy2 | 7.19669 | 8.80681 |
| PA14_33330 | csy1 | 7.64528 | 9.26927 |
| PA14_33340 | cas3f | 6.30742 | 7.899 |
| PA14_33350 | cas1f | 6.91796 | 8.00503 |
| PA14_33370 | | -0.89301 | -1.01463 |
| PA14_33450 | treA | 0.77192 | 0.68532 |
| PA14_33460 | | 0.92885 | 0.59117 |
| PA14_33480 | sndH | 0.63865 | 0.76169 |
| PA14_33500 | pvdH | -0.49768 | |
| PA14_33510 | | -0.45647 | |
| PA14_33540 | | 0.43424 | 0.48013 |
| PA14_33680 | fpvA | | |
| PA14_33750 | ompQ | | |
| PA14_33820 | pvdQ | | 0.94987 |
| PA14_33830 | | | |
| PA14_33840 | | | |
| PA14_33900 | | -0.5174 | -0.60568 |
| PA14_33910 | | | 1.21412 |
| PA14_33920 | | -2.30769 | -1.0663 |
| PA14_33940 | | | 1.42439 |
| PA14_33960 | vgrG3 | | 1.34483 |
| PA14_33970 | tpe2 | 5.74793 | 7.87136 |
| PA14_33980 | tsd2 | 5.3162 | 7.85355 |
| PA14_33990 | clpV3 | -0.59053 | 0.94566 |
| PA14_34000 | hsiH3 | | 1.27309 |
| PA14_34010 | hsiG3 | | 1.77251 |
| PA14_34020 | hsiF3 | | 2.10123 |
| PA14_34030 | hcp3 | | 1.99139 |
| PA14_34050 | hsiC3 | | 2.14778 |
| PA14_34070 | hsiB3 | | 2.32192 |
| PA14_34100 | hsiJ3 | | 1.06324 |
| PA14_34110 | dotU3 | | 1.9899 |
| PA14_34130 | icmF3 | | 1.42208 |
| PA14_34140 | hsiA3 | -0.40818 | 1.04539 |
| PA14_34150 | sfa3 | -4.5599 | -4.11973 |
| PA14_34170 | | -2.30522 | -0.96872 |
| PA14_34320 | | | -0.73483 |

| | | | |
|---|---|---|---|
| PA14_34330 | | 0.38766 | |
| PA14_34340 | mtlZ | -1.33639 | -1.6165 |
| PA14_34390 | | | -0.67759 |
| PA14_34450 | | -0.56438 | -0.81591 |
| PA14_34460 | | -2.01441 | |
| PA14_34490 | | -0.89788 | |
| PA14_34500 | | -0.69529 | |
| PA14_34510 | | -1.00452 | |
| PA14_34520 | | -1.3837 | |
| PA14_34540 | | | 0.62322 |
| PA14_34600 | | 0.63646 | 0.79114 |
| PA14_34630 | gnuT | 0.91546 | 0.56301 |
| PA14_34670 | | -0.92436 | -0.64212 |
| PA14_34680 | | -0.57969 | -0.61097 |
| PA14_34690 | | -1.49157 | -1.79351 |
| PA14_34730 | | -3.8801 | -3.9561 |
| PA14_34740 | | -4.20297 | -4.40392 |
| PA14_34750 | | -5.22202 | -4.38484 |
| PA14_34770 | | -3.7607 | -2.55856 |
| PA14_34780 | | -3.1347 | -1.36539 |
| PA14_34790 | | -1.28346 | |
| PA14_34800 | | 1.1308 | 1.17692 |
| PA14_34810 | | -0.43904 | 0.54116 |
| PA14_34820 | | -0.53876 | 0.91286 |
| PA14_34830 | | -0.66553 | 0.85699 |
| PA14_34840 | | -0.67696 | 0.54222 |
| PA14_34850 | | -0.45951 | |
| PA14_34870 | chiC | 2.56267 | 4.23469 |
| PA14_34880 | | -4.88579 | -4.32401 |
| PA14_34900 | | -4.4898 | -3.99195 |
| PA14_34920 | | -4.46018 | -3.42029 |
| PA14_34930 | | | |
| PA14_34960 | oprB | 0.9408 | 1.30437 |
| PA14_34970 | gcd | | 0.87346 |
| PA14_34990 | | 0.63803 | 1.26081 |
| PA14_35050 | | | -0.8204 |
| PA14_35060 | | -0.76498 | -0.93275 |
| PA14_35150 | | | 0.76225 |
| PA14_35160 | pumA | 0.83251 | 2.02798 |
| PA14_35190 | | | |
| PA14_35200 | | | 0.58428 |
| PA14_35400 | pvcC | 0.9185 | |
| PA14_35440 | ansA | -0.91591 | -0.51642 |
| PA14_35620 | pslK | -0.54184 | -0.71406 |
| PA14_35630 | pslJ | -1.3333 | -0.80224 |
| PA14_35640 | pslI | -1.51775 | -0.71816 |
| PA14_35650 | pslH | -1.45942 | -0.63747 |
| PA14_35670 | pslG | -1.57646 | -1.07067 |

| | | | |
|---|---|---|---|
| PA14_35680 | psIF | -3.10361 | -3.16284 |
| PA14_35690 | psIE | -3.44548 | -3.43142 |
| PA14_35710 | | 5.28561 | 4.67684 |
| PA14_35720 | | 4.82735 | 4.93468 |
| PA14_35730 | | 1.83246 | 2.17962 |
| PA14_35740 | | 4.50718 | 3.63985 |
| PA14_35750 | tnpA | 5.29985 | 4.30796 |
| PA14_35760 | | 3.2653 | 2.79592 |
| PA14_35770 | | 3.30017 | 3.56952 |
| PA14_35780 | | 1.85621 | 5.53869 |
| PA14_35790 | | 4.4391 | 7.51059 |
| PA14_35800 | | 7.02785 | 8.45276 |
| PA14_35810 | | | 5.04673 |
| PA14_35820 | tnpS | 8.56759 | 9.50512 |
| PA14_35830 | tnpT | 7.41744 | 8.19581 |
| PA14_35840 | | 9.26078 | 10.05458 |
| PA14_35850 | | 9.76632 | 10.20457 |
| PA14_35860 | | 2.42195 | 2.38876 |
| PA14_35880 | | 4.87366 | 4.89435 |
| PA14_35890 | | 6.93699 | 7.04583 |
| PA14_35900 | | 6.07361 | 4.83542 |
| PA14_35920 | | 1.27205 | 0.88252 |
| PA14_35940 | | 0.97574 | 1.38871 |
| PA14_35970 | | 1.27559 | 2.32951 |
| PA14_35990 | | 0.86825 | |
| PA14_36000 | prpR | 5.99189 | 4.69476 |
| PA14_36010 | | 3.67878 | 3.5915 |
| PA14_36020 | | -1.38762 | -1.53835 |
| PA14_36030 | | 2.44791 | 2.05737 |
| PA14_36180 | | | -0.47111 |
| PA14_36200 | | -2.91839 | -1.0518 |
| PA14_36220 | | -0.82204 | |
| PA14_36250 | | | 0.57632 |
| PA14_36270 | | -0.74029 | |
| PA14_36280 | | | 0.98953 |
| PA14_36290 | | | 1.29445 |
| PA14_36300 | | | 1.81436 |
| PA14_36310 | hcnC | | 2.25543 |
| PA14_36320 | hcnB | | 2.32514 |
| PA14_36330 | hcnA | | 1.70439 |
| PA14_36345 | exoY | 1.67408 | 0.96045 |
| PA14_36360 | | 0.61666 | |
| PA14_36390 | | 0.67429 | |
| PA14_36450 | | 0.68965 | |
| PA14_36570 | glgA | 0.53613 | 0.62062 |
| PA14_36580 | treZ | 1.21986 | 1.10392 |
| PA14_36590 | malQ | 1.50458 | 1.24381 |
| PA14_36605 | | 1.22665 | 1.12302 |

| | | | |
|---|---|---|---|
| PA14_36620 | | 0.76232 | 0.72198 |
| PA14_36630 | glgX | 0.83254 | 0.74103 |
| PA14_36650 | | 1.34794 | |
| PA14_36660 | | 1.18287 | 0.74207 |
| PA14_36670 | | 1.18442 | 0.7174 |
| PA14_36680 | | 0.79803 | 0.62379 |
| PA14_36690 | clsB | 0.83368 | 0.71747 |
| PA14_36700 | | 0.62876 | |
| PA14_36740 | | 0.74059 | 0.74041 |
| PA14_36810 | katE | 1.17553 | 1.17438 |
| PA14_36840 | glgP | 0.51759 | |
| PA14_36870 | | | 0.63345 |
| PA14_36910 | ligD | 0.72315 | |
| PA14_36920 | | 0.79257 | |
| PA14_37070 | | | 0.9049 |
| PA14_37120 | | -0.6242 | -0.83795 |
| PA14_37130 | | -0.74288 | -0.76839 |
| PA14_37140 | | -1.91707 | -1.53957 |
| PA14_37170 | | 5.83864 | 4.20247 |
| PA14_37250 | | -2.53846 | -2.43473 |
| PA14_37410 | | -1.33246 | -0.95692 |
| PA14_37420 | | | 3.13516 |
| PA14_37430 | | | 1.51172 |
| PA14_37440 | | | 2.91566 |
| PA14_37470 | | | 3.40955 |
| PA14_37490 | | | 3.30298 |
| PA14_37510 | | | 3.45722 |
| PA14_37560 | asnB | | 2.94481 |
| PA14_37570 | | -0.60013 | 1.47663 |
| PA14_37660 | | | |
| PA14_37670 | | 2.12684 | 2.27084 |
| PA14_37680 | | 1.57031 | 1.16864 |
| PA14_37690 | | | 0.8851 |
| PA14_37710 | fusA2 | | 0.48031 |
| PA14_37730 | | -0.44064 | |
| PA14_37745 | | 2.3563 | 2.8332 |
| PA14_37760 | | 2.57262 | 2.9115 |
| PA14_37770 | | 2.3356 | 2.88012 |
| PA14_37780 | | 2.6486 | 3.11654 |
| PA14_37820 | | -0.61577 | |
| PA14_37830 | | -3.40797 | -1.52707 |
| PA14_37870 | sppB | -1.02831 | -0.60987 |
| PA14_37910 | | | -0.74262 |
| PA14_37940 | cynR | -0.86453 | -0.87054 |
| PA14_37965 | cynS | | |
| PA14_37980 | | | |
| PA14_37990 | | -1.96877 | -0.63242 |
| PA14_38010 | | | |

| | | | |
|---|---|---|---|
| PA14_38040 | cmrA | | 0.69234 |
| PA14_38140 | | -0.43583 | -1.36416 |
| PA14_38200 | | 0.37858 | 0.66309 |
| PA14_38260 | | 0.44983 | 0.62171 |
| PA14_38270 | | 0.92978 | 1.1123 |
| PA14_38320 | | 0.73757 | 0.80502 |
| PA14_38350 | galU | 0.51104 | |
| PA14_38370 | | -1.54934 | -1.45441 |
| PA14_38420 | | | -0.95664 |
| PA14_38430 | gnyR | 0.67098 | 0.83057 |
| PA14_38440 | gnyD | 0.4684 | |
| PA14_38470 | gnyH | 0.52276 | 0.81371 |
| PA14_38490 | gnyL | | 0.57214 |
| PA14_38510 | hmgA | | -0.75219 |
| PA14_38530 | fahA | | -0.51085 |
| PA14_38570 | hbcR | -1.29062 | -1.3981 |
| PA14_38590 | bdhA | -1.25214 | -0.8194 |
| PA14_38680 | | | -0.54879 |
| PA14_38780 | pqqE | | 0.83819 |
| PA14_38970 | nahK | -4.50903 | -0.56312 |
| PA14_38990 | nosP | -3.94973 | |
| PA14_39050 | braZ | 1.6265 | 0.854 |
| PA14_39060 | | 1.35035 | 3.35854 |
| PA14_39080 | | | -1.52615 |
| PA14_39090 | | | 0.80955 |
| PA14_39100 | | 0.94768 | |
| PA14_39130 | | | |
| PA14_39180 | | 0.49309 | 0.77995 |
| PA14_39240 | | -0.52737 | |
| PA14_39270 | | | 0.96509 |
| PA14_39280 | rbsK | -1.49048 | -1.24937 |
| PA14_39330 | rbsA | | |
| PA14_39350 | rbsB | 1.18619 | |
| PA14_39440 | | | 0.78899 |
| PA14_39460 | | | |
| PA14_39470 | | 4.35808 | 4.11071 |
| PA14_39480 | | 5.60349 | 3.53436 |
| PA14_39520 | | | 0.58837 |
| PA14_39560 | | | 0.88578 |
| PA14_39590 | metE | -3.3076 | -3.59808 |
| PA14_39620 | | -0.65879 | -0.68467 |
| PA14_39630 | | -0.80369 | -0.80459 |
| PA14_39640 | cobN | | -0.65387 |
| PA14_39770 | | -0.47619 | -0.52117 |
| PA14_39780 | | 2.0339 | 3.33702 |
| PA14_39880 | | | 4.21481 |
| PA14_39890 | phzF2 | 3.36183 | 4.93625 |
| PA14_39945 | phzC2 | 1.41543 | 3.57506 |

| | | | |
|---|---|---|---|
| PA14_39960 | phzB2 | 3.04792 | 4.99065 |
| PA14_39970 | phzA2 | 4.5315 | 4.80025 |
| PA14_39990 | | | 1.08415 |
| PA14_40010 | | 0.57263 | 2.77692 |
| PA14_40020 | | 1.52128 | 3.77691 |
| PA14_40030 | | 1.23771 | 3.5467 |
| PA14_40040 | | 0.61993 | 2.62909 |
| PA14_40050 | | 0.52021 | 2.72457 |
| PA14_40060 | | 1.80589 | 4.37681 |
| PA14_40100 | | -0.60866 | 1.0442 |
| PA14_40110 | | -0.68608 | |
| PA14_40180 | | | 0.83839 |
| PA14_40200 | | | 0.8304 |
| PA14_40240 | | -2.20601 | -1.66566 |
| PA14_40260 | | 0.61417 | 1.59374 |
| PA14_40290 | lasA | 4.53112 | 5.46836 |
| PA14_40300 | | 0.50515 | 0.95893 |
| PA14_40310 | acpP | 0.99174 | 1.00125 |
| PA14_40470 | | 5.38461 | 2.69043 |
| PA14_40560 | | | 0.55624 |
| PA14_40630 | nfuA | -0.48147 | -0.51482 |
| PA14_40650 | tsi1 | | |
| PA14_40660 | tse1 | | 1.29122 |
| PA14_40670 | metH | -2.07209 | -1.7221 |
| PA14_40690 | | -2.71708 | -2.04181 |
| PA14_40700 | | | -0.6869 |
| PA14_40730 | | -0.52761 | -0.70293 |
| PA14_40770 | cysI | -0.9598 | |
| PA14_40780 | | -1.01322 | |
| PA14_40790 | | -0.41337 | -1.04984 |
| PA14_40840 | sohB | | -0.46143 |
| PA14_40880 | | 0.46011 | 0.64307 |
| PA14_40890 | | | 0.65876 |
| PA14_40910 | | -0.65041 | -0.55235 |
| PA14_41030 | | -0.68418 | -1.07873 |
| PA14_41090 | mltD | 0.40766 | |
| PA14_41130 | nppA2 | | 0.69182 |
| PA14_41160 | nppD | | 0.86392 |
| PA14_41170 | fabI | | 0.70185 |
| PA14_41250 | tig | | -0.51051 |
| PA14_41270 | parS | -0.51169 | -0.47464 |
| PA14_41420 | | | 1.21962 |
| PA14_41440 | | | 0.79289 |
| PA14_41470 | acnB | -0.71116 | |
| PA14_41500 | | 0.85901 | 0.92739 |
| PA14_41560 | | -2.47083 | -2.54808 |
| PA14_41563 | cobA | -0.7597 | -0.86662 |
| PA14_41575 | sigX | 0.72169 | |

| | | | |
|---|---|---|---|
| PA14_41730 | | | |
| PA14_41810 | | 0.48965 | |
| PA14_41840 | cysH | -1.50967 | -1.79235 |
| PA14_41870 | cysB | -1.10646 | -0.95568 |
| PA14_41960 | | -2.37623 | -1.19748 |
| PA14_41990 | | | 0.87456 |
| PA14_42030 | | -0.96338 | -0.97383 |
| PA14_42050 | | -0.58593 | -1.12447 |
| PA14_42060 | | -2.25379 | -2.61211 |
| PA14_42130 | | | 0.49694 |
| PA14_42180 | | | -0.41325 |
| PA14_42200 | | -0.70733 | -0.9998 |
| PA14_42220 | | | |
| PA14_42270 | pscJ | | |
| PA14_42290 | pscH | | |
| PA14_42340 | pscD | | |
| PA14_42350 | pscC | | -0.76938 |
| PA14_42360 | pscB | | -1.73668 |
| PA14_42380 | exsD | | -1.03025 |
| PA14_42390 | exsA | -0.62691 | -1.63428 |
| PA14_42400 | exsB | | -1.23469 |
| PA14_42410 | exsE | | -0.89167 |
| PA14_42430 | exsC | 1.28477 | |
| PA14_42440 | popD | 1.93363 | -0.78008 |
| PA14_42450 | popB | 1.90051 | -1.16079 |
| PA14_42460 | pcrH | 1.36892 | |
| PA14_42470 | pcrV | 1.34927 | -0.8886 |
| PA14_42480 | pcrG | 2.30312 | |
| PA14_42520 | pscX | | |
| PA14_42550 | popN | | |
| PA14_42570 | pscN | | |
| PA14_42600 | pscP | | -0.72996 |
| PA14_42610 | pscQ | | |
| PA14_42620 | pscR | -6.02062 | -7.97201 |
| PA14_42670 | | 0.59668 | |
| PA14_42780 | | | -0.69186 |
| PA14_42860 | mhr | | 0.70763 |
| PA14_42870 | | | -0.43093 |
| PA14_42880 | stk1 | 1.33753 | 5.29255 |
| PA14_42960 | lip2.2 | | 2.69891 |
| PA14_43000 | hsiG2 | 0.80774 | 7.05699 |
| PA14_43020 | hsiF2 | 0.77309 | 7.35911 |
| PA14_43030 | hsiC2 | | 8.25299 |
| PA14_43040 | hsiB2 | | 7.34404 |
| PA14_43050 | hsiA2 | 3.18766 | 6.10131 |
| PA14_43070 | hcp2 | 4.3358 | 9.17491 |
| PA14_43080 | vgrG14 | | 7.0889 |
| PA14_43090 | tap | 2.64059 | 8.57082 |

| | | | |
|---|---|---|---|
| PA14_43100 | rhsP2 | | 3.21788 |
| PA14_43190 | | | 0.86874 |
| PA14_43230 | | | |
| PA14_43240 | | -0.38017 | |
| PA14_43250 | | | |
| PA14_43270 | mnmH | -0.32272 | |
| PA14_43310 | | -0.5984 | |
| PA14_43420 | | 0.69894 | 0.88827 |
| PA14_43440 | | -0.62647 | -0.82996 |
| PA14_43510 | | 0.58593 | |
| PA14_43600 | | | -0.45735 |
| PA14_43650 | | -0.63278 | |
| PA14_43660 | | | |
| PA14_43670 | | | |
| PA14_43710 | | -1.50487 | -1.49783 |
| PA14_43720 | bosR | -2.39505 | -2.37066 |
| PA14_43760 | | -0.6392 | |
| PA14_43830 | panB | | |
| PA14_43850 | hptG | | 0.7035 |
| PA14_43880 | | | -0.81047 |
| PA14_44020 | sdhB | -0.92545 | -0.52768 |
| PA14_44050 | sdhD | -0.50733 | -0.92413 |
| PA14_44060 | sdhC | -0.6891 | -1.066 |
| PA14_44090 | | -0.6345 | -1.03114 |
| PA14_44110 | | -1.06655 | -2.12292 |
| PA14_44120 | | 0.46005 | |
| PA14_44150 | | | |
| PA14_44160 | | | -0.46482 |
| PA14_44180 | | -0.92617 | -1.36459 |
| PA14_44210 | | 0.89317 | 0.88353 |
| PA14_44240 | | 0.69576 | |
| PA14_44311 | cprA | | -0.45576 |
| PA14_44390 | | | |
| PA14_44400 | ccoP | | 0.71913 |
| PA14_44420 | ccoG | | -0.61211 |
| PA14_44450 | ccoS | -0.87309 | -1.09392 |
| PA14_44480 | pemB | -1.35515 | -1.88887 |
| PA14_44490 | anr | -0.81357 | -0.59925 |
| PA14_44640 | | -0.56733 | -0.62985 |
| PA14_44650 | | | 0.62386 |
| PA14_44660 | ligA | 0.52737 | 0.77941 |
| PA14_44800 | | | -0.66734 |
| PA14_44830 | puuE | | |
| PA14_44840 | uraD | | |
| PA14_44850 | alc | 0.56711 | |
| PA14_44880 | | 0.72528 | |
| PA14_44910 | | | 5.10772 |
| PA14_44950 | | | 2.01905 |

| | | | |
|---|---|---|---|
| PA14_44970 | moaA2 | -0.50719 | -0.95301 |
| PA14_45100 | muiA | | 1.01776 |
| PA14_45110 | cysP | -2.59874 | -2.90128 |
| PA14_45120 | | 0.63371 | 0.63279 |
| PA14_45130 | eutH | | |
| PA14_45240 | | | -0.68205 |
| PA14_45400 | | -0.40091 | |
| PA14_45410 | | -0.59075 | |
| PA14_45440 | | | -0.88612 |
| PA14_45500 | | -0.75245 | -0.69226 |
| PA14_45590 | | -0.33587 | |
| PA14_45700 | | | 1.25555 |
| PA14_45710 | | 1.06322 | 1.34026 |
| PA14_45760 | fliQ | -0.61329 | -0.52853 |
| PA14_45790 | fliN | -1.12659 | -1.0505 |
| PA14_45800 | fliM | -0.54953 | -0.63118 |
| PA14_45810 | fliL | -0.52391 | -0.67989 |
| PA14_45830 | | -1.68645 | -1.40648 |
| PA14_45840 | | -0.37994 | -0.51692 |
| PA14_45940 | lasI | -1.5257 | -0.53915 |
| PA14_46030 | bdlA | -2.06516 | -1.44529 |
| PA14_46070 | gbuA | | -2.52333 |
| PA14_46080 | | -0.58376 | -2.22007 |
| PA14_46100 | | | -1.03446 |
| PA14_46110 | | | -0.88931 |
| PA14_46120 | | | -0.64142 |
| PA14_46230 | aphA | | 0.54439 |
| PA14_46300 | | 0.74393 | 0.48831 |
| PA14_46340 | | | -0.49806 |
| PA14_46380 | | | 0.63529 |
| PA14_46400 | | -1.12117 | -0.99186 |
| PA14_46430 | | | 0.88276 |
| PA14_46440 | | | 0.47827 |
| PA14_46450 | aceK | | 0.47112 |
| PA14_46460 | | 3.63888 | 4.17852 |
| PA14_46510 | | 5.80207 | 5.74967 |
| PA14_46520 | | | 4.09329 |
| PA14_46530 | | 3.61862 | 4.7731 |
| PA14_46540 | | 5.87046 | 5.60902 |
| PA14_46550 | | 7.37417 | 8.13785 |
| PA14_46560 | | 1.97111 | 2.40113 |
| PA14_46570 | | 7.58055 | 7.70365 |
| PA14_46580 | | 3.87321 | 3.22761 |
| PA14_46590 | | 3.58197 | 3.14728 |
| PA14_46610 | | 4.37745 | 4.48259 |
| PA14_46620 | | 6.27044 | 5.32771 |
| PA14_46630 | | -0.57642 | -0.5817 |
| PA14_46640 | | | -0.74478 |

| | | | |
|---|---|---|---|
| PA14_46650 | | | -1.25508 |
| PA14_46660 | | -0.48096 | -1.97305 |
| PA14_46700 | | | -0.56798 |
| PA14_46720 | | | 0.66404 |
| PA14_46750 | | 0.60273 | 1.05104 |
| PA14_46810 | | 0.928 | 1.03635 |
| PA14_46820 | | 0.57996 | 1.06314 |
| PA14_46830 | | 0.52941 | 0.83011 |
| PA14_46850 | | 1.42302 | |
| PA14_46860 | | 1.42025 | 0.53852 |
| PA14_46880 | | 0.58152 | 0.51976 |
| PA14_46890 | | | 0.55348 |
| PA14_46920 | | 0.56018 | 0.68229 |
| PA14_46970 | ansB | | 0.76192 |
| PA14_47030 | | -0.39961 | |
| PA14_47090 | | | |
| PA14_47100 | ilvA2 | 0.54472 | 0.46247 |
| PA14_47110 | | 0.98556 | 0.67662 |
| PA14_47120 | | | 0.90836 |
| PA14_47150 | cyoE | | -1.09613 |
| PA14_47160 | cyoD | | -1.77184 |
| PA14_47180 | cyoC | | -0.9806 |
| PA14_47190 | cyoB | | -0.71619 |
| PA14_47210 | cyoA | | -1.54671 |
| PA14_47230 | | | -0.57114 |
| PA14_47320 | | -0.34327 | |
| PA14_47370 | lepB | | |
| PA14_47390 | | -0.36986 | -0.42563 |
| PA14_47400 | | -0.60456 | -0.84016 |
| PA14_47520 | | 0.45312 | 1.2933 |
| PA14_47530 | | 1.37252 | 2.0059 |
| PA14_47550 | | 0.6725 | |
| PA14_47580 | | -0.75438 | |
| PA14_47640 | | | -1.16286 |
| PA14_47690 | cobQ | | 0.62473 |
| PA14_47720 | cobC | | 0.59928 |
| PA14_47730 | cobD | | 0.70263 |
| PA14_47760 | cobB | | 0.74092 |
| PA14_47810 | | -0.89493 | -1.50009 |
| PA14_47860 | | 3.27935 | 3.22449 |
| PA14_47900 | | -0.53919 | |
| PA14_47930 | | -0.61545 | |
| PA14_47940 | | -0.63584 | -0.87537 |
| PA14_48010 | | 1.45652 | 1.05218 |
| PA14_48040 | aprI | 0.71814 | 0.53182 |
| PA14_48060 | aprA | 0.94769 | 1.27088 |
| PA14_48090 | aprF | | -0.78257 |
| PA14_48100 | aprE | | -0.698 |

| | | | |
|---|---|---|---|
| PA14_48115 | aprD | | -0.53374 |
| PA14_48150 | qslA | -6.68518 | -5.5418 |
| PA14_48450 | | 5.75638 | 5.01124 |
| PA14_48460 | | 2.75769 | 2.64546 |
| PA14_48470 | aguB | 1.43696 | 1.05231 |
| PA14_48490 | | 4.14312 | 3.78539 |
| PA14_48510 | dnaB | 3.46949 | 3.3045 |
| PA14_48520 | | 0.43037 | |
| PA14_48530 | | | 0.87138 |
| PA14_48570 | | | 0.58108 |
| PA14_48590 | | | 1.16776 |
| PA14_48750 | | | |
| PA14_48760 | | | 0.82982 |
| PA14_48830 | ddaR | | 0.96693 |
| PA14_48860 | | 0.42261 | |
| PA14_48880 | | 4.97417 | 4.9779 |
| PA14_48890 | | -1.73386 | -2.04058 |
| PA14_48930 | | 0.65396 | 0.62486 |
| PA14_48940 | coaB | -2.82857 | -3.99278 |
| PA14_48950 | | -1.47045 | -2.6799 |
| PA14_48960 | | -2.15004 | -2.37904 |
| PA14_48970 | | -2.47343 | -3.7072 |
| PA14_48980 | | -5.03445 | -7.16463 |
| PA14_49030 | | 2.64797 | 2.62943 |
| PA14_49130 | dctA | | -0.90489 |
| PA14_49160 | | 0.35163 | |
| PA14_49170 | phoQ | 0.35432 | |
| PA14_49200 | oprH | -0.42299 | |
| PA14_49230 | napD | | 1.13337 |
| PA14_49250 | napA | | 0.95484 |
| PA14_49290 | | | |
| PA14_49300 | | 3.57509 | 2.95596 |
| PA14_49310 | | 4.26144 | 3.60163 |
| PA14_49320 | | 1.78753 | 1.91392 |
| PA14_49410 | | | -0.61202 |
| PA14_49480 | | 2.33058 | 4.627 |
| PA14_49500 | | 0.88567 | 3.31212 |
| PA14_49510 | | 6.20958 | 6.24197 |
| PA14_49520 | pyoS3A | 3.55062 | 4.34437 |
| PA14_49530 | | | 0.58401 |
| PA14_49540 | | | 4.37829 |
| PA14_49560 | toxA | 1.21164 | 0.63292 |
| PA14_49570 | | -1.71918 | -2.76308 |
| PA14_49650 | | | -0.56031 |
| PA14_49700 | | | -0.4846 |
| PA14_49710 | hchA | -1.45275 | |
| PA14_49720 | | -0.71667 | |
| PA14_49740 | | 0.95666 | 1.91029 |

| | | | |
|---|---|---|---|
| PA14_49750 | | | -0.58633 |
| PA14_49760 | rhlC | -1.45086 | -0.76414 |
| PA14_49790 | | | -0.58855 |
| PA14_49800 | | | -0.70131 |
| PA14_49870 | def | -0.87395 | -0.80857 |
| PA14_49880 | yfiR | 1.2167 | 1.46443 |
| PA14_49890 | tobB | 0.58303 | 0.59491 |
| PA14_49900 | | 0.5925 | 1.23344 |
| PA14_49920 | | 0.63512 | 0.61375 |
| PA14_49990 | | -0.63615 | -0.75234 |
| PA14_50030 | | | -0.61517 |
| PA14_50110 | fliH | | -0.45025 |
| PA14_50220 | fleQ | -0.49413 | |
| PA14_50240 | fleP | -1.32287 | -1.25792 |
| PA14_50250 | fliS | -1.11395 | -0.91314 |
| PA14_50270 | fliD | -0.71444 | -0.42286 |
| PA14_50280 | | -0.52724 | -0.46115 |
| PA14_50290 | fliC | -1.27968 | -1.21036 |
| PA14_50300 | | -0.4966 | |
| PA14_50310 | | -0.54279 | |
| PA14_50320 | | -0.68558 | |
| PA14_50330 | | -1.34442 | -0.83668 |
| PA14_50340 | flgL | -0.90828 | |
| PA14_50360 | flgK | -0.95219 | |
| PA14_50380 | flgJ | -0.6357 | |
| PA14_50410 | flgI | -0.99023 | -0.76029 |
| PA14_50420 | flgH | -0.82541 | -0.57925 |
| PA14_50430 | flgG | -1.14067 | -0.78143 |
| PA14_50440 | flgF | -1.25849 | -0.85098 |
| PA14_50450 | flgE | -1.53177 | -0.60899 |
| PA14_50460 | flgD | -0.95068 | |
| PA14_50470 | flgC | -1.16183 | -0.69785 |
| PA14_50480 | flgB | -1.31892 | -0.74934 |
| PA14_50500 | | | 0.74043 |
| PA14_50570 | | | 1.44996 |
| PA14_50590 | | | 1.62794 |
| PA14_50610 | | -0.53465 | -0.60632 |
| PA14_50710 | shaD | | -0.51867 |
| PA14_50720 | shaE | -0.49124 | -0.66545 |
| PA14_50730 | shaF | -0.5487 | -1.05498 |
| PA14_50760 | | | -0.61296 |
| PA14_50820 | | 1.23659 | 1.06024 |
| PA14_50850 | | | -0.77 |
| PA14_50860 | | 0.58192 | 0.68515 |
| PA14_50880 | | | |
| PA14_50980 | pac | | 0.41184 |
| PA14_51080 | | -1.08421 | -1.06702 |
| PA14_51090 | | -1.05207 | -1.01386 |

| | | | |
|---|---|---|---|
| PA14_51100 | | -0.71753 | -0.52725 |
| PA14_51110 | | -1.50012 | -1.93866 |
| PA14_51120 | | -2.34163 | -2.47047 |
| PA14_51330 | nadA | -0.64889 | -0.9606 |
| PA14_51340 | mvfR | -0.31144 | |
| PA14_51350 | phnB | 2.57971 | 3.81032 |
| PA14_51360 | phnA | 2.49165 | 3.77357 |
| PA14_51380 | pqsE | 2.46617 | 3.66069 |
| PA14_51390 | pqsD | 2.62385 | 3.99755 |
| PA14_51410 | pqsC | 3.06462 | 4.45668 |
| PA14_51420 | pqsB | 3.27367 | 4.50951 |
| PA14_51430 | pqsA | 2.94075 | 3.78845 |
| PA14_51440 | ogt | -1.35124 | -1.82844 |
| PA14_51450 | cupC3 | 0.75488 | 0.7191 |
| PA14_51460 | cupC2 | | |
| PA14_51490 | | 0.5706 | |
| PA14_51500 | | 0.68422 | 1.38274 |
| PA14_51520 | spcU | 5.80831 | 3.32338 |
| PA14_51530 | exoU | 6.71974 | 5.36287 |
| PA14_51540 | | -1.63823 | -1.95387 |
| PA14_51550 | | 4.4745 | 4.24715 |
| PA14_51560 | | 5.6047 | 4.97607 |
| PA14_51610 | | 3.01305 | 3.20789 |
| PA14_51640 | | 1.53179 | 0.89561 |
| PA14_51650 | | 2.27027 | 1.47472 |
| PA14_51680 | queE | | |
| PA14_51710 | oprL | -0.39699 | |
| PA14_51880 | oprD | | 1.01056 |
| PA14_51950 | | -1.27144 | |
| PA14_52010 | hda | | -0.4518 |
| PA14_52090 | | | -0.68359 |
| PA14_52120 | | | -1.36384 |
| PA14_52210 | cysM | | -0.48449 |
| PA14_52250 | pirR | | -0.80252 |
| PA14_52270 | ldhA | | 0.44545 |
| PA14_52330 | | | -0.59636 |
| PA14_52460 | mgtE | | -0.70948 |
| PA14_52490 | | | 0.78834 |
| PA14_52500 | | | 0.91967 |
| PA14_52510 | | | 1.05414 |
| PA14_52550 | | -3.63714 | -1.27374 |
| PA14_52570 | rsmA | | -8.42682 |
| PA14_52580 | lysC | -0.51861 | |
| PA14_52670 | astD | | 0.52922 |
| PA14_52730 | | | -0.59479 |
| PA14_52750 | aotP | 0.6329 | 0.72957 |
| PA14_52760 | | 0.79064 | 1.07819 |
| PA14_52770 | aotM | 0.78349 | 0.62589 |

| | | | |
|---|---|---|---|
| PA14_52780 | aotQ | 1.12696 | 0.79247 |
| PA14_52820 | | 1.04902 | 1.26609 |
| PA14_52870 | | -0.67412 | -1.23083 |
| PA14_52880 | | | -1.05302 |
| PA14_52910 | | | -0.75545 |
| PA14_52920 | | -0.55226 | -1.04995 |
| PA14_52930 | | -0.81179 | -1.09709 |
| PA14_52980 | phhR | | |
| PA14_53020 | pbpG | | -0.54398 |
| PA14_53030 | arfB | | -0.52344 |
| PA14_53040 | | | 0.64499 |
| PA14_53050 | aroP2 | -2.30359 | -2.39731 |
| PA14_53070 | hpd | 1.04422 | 0.99911 |
| PA14_53110 | | 0.52673 | |
| PA14_53150 | | 1.18009 | 0.54192 |
| PA14_53160 | | 1.26879 | 0.99602 |
| PA14_53180 | | 0.84158 | |
| PA14_53190 | bolA | 0.59305 | |
| PA14_53200 | | 0.39458 | |
| PA14_53210 | | 1.36975 | 2.48111 |
| PA14_53220 | fumC2 | 0.40571 | 1.30682 |
| PA14_53230 | | | 0.56841 |
| PA14_53250 | cpbD | 2.39222 | 3.46092 |
| PA14_53260 | | 0.91689 | |
| PA14_53270 | | 0.73238 | 0.7496 |
| PA14_53300 | | | 0.94643 |
| PA14_53340 | cerN | 0.61692 | |
| PA14_53400 | | -0.73774 | |
| PA14_53410 | | -1.10415 | -0.87029 |
| PA14_53470 | | | 0.47395 |
| PA14_53480 | pta | | 0.6871 |
| PA14_53490 | | 1.01347 | |
| PA14_53510 | | -0.43577 | -0.58842 |
| PA14_53520 | oruR | -0.8598 | -1.34067 |
| PA14_53530 | | | 0.73583 |
| PA14_53570 | | 4.02526 | 4.41345 |
| PA14_53580 | | 3.85526 | 3.75709 |
| PA14_53590 | | 4.64968 | 3.886 |
| PA14_53600 | | -2.5056 | -3.12846 |
| PA14_53610 | | 0.71425 | |
| PA14_53640 | | | |
| PA14_53690 | | -1.84942 | -2.40469 |
| PA14_53700 | | | 0.69887 |
| PA14_53790 | | 1.19937 | |
| PA14_53820 | ampDh3 | 0.38937 | 1.09097 |
| PA14_53840 | | | -0.79454 |
| PA14_53870 | | -0.65648 | |
| PA14_53880 | | | -0.47064 |

| | | | |
|---|---|---|---|
| PA14_54000 | prpD | 0.34392 | 1.28355 |
| PA14_54050 | | 6.24185 | 4.55411 |
| PA14_54070 | | 0.52866 | |
| PA14_54110 | | | -1.1821 |
| PA14_54120 | azoR1 | -1.75891 | -1.29995 |
| PA14_54130 | | -4.05542 | -4.0586 |
| PA14_54150 | putP | 0.6011 | |
| PA14_54170 | putA | | |
| PA14_54190 | pruR | | -0.50027 |
| PA14_54210 | lon | 0.43087 | 1.15475 |
| PA14_54240 | | | 0.72174 |
| PA14_54320 | era | 0.42448 | 0.52588 |
| PA14_54330 | rnc | 0.48535 | 0.61838 |
| PA14_54350 | lepB | 0.37831 | |
| PA14_54370 | lepA | 0.4471 | |
| PA14_54390 | mucD | 1.06074 | 0.77822 |
| PA14_54410 | mucB | | |
| PA14_54430 | algU | | -0.72565 |
| PA14_54450 | nadB | -0.42189 | -0.47069 |
| PA14_54470 | | | -0.79369 |
| PA14_54500 | | 0.50146 | |
| PA14_54520 | | 0.57873 | |
| PA14_54570 | | | |
| PA14_54580 | | | -0.60575 |
| PA14_54590 | ung | | |
| PA14_54600 | | -0.52832 | 0.5133 |
| PA14_54610 | | -0.54709 | -1.07851 |
| PA14_54630 | | 0.47968 | 0.67721 |
| PA14_54660 | | 0.7974 | 0.96089 |
| PA14_54690 | | | 0.91775 |
| PA14_54720 | | | -1.62454 |
| PA14_54730 | | -0.98666 | -2.64965 |
| PA14_54740 | | -0.558 | -2.08785 |
| PA14_54750 | | | -1.7095 |
| PA14_54760 | | -0.61129 | |
| PA14_54770 | | | 0.95743 |
| PA14_54810 | | | |
| PA14_54830 | phaG | 0.85562 | 2.52567 |
| PA14_54850 | | 7.76174 | 7.94068 |
| PA14_54860 | | | 2.97174 |
| PA14_54880 | cysE | 5.46368 | 5.32193 |
| PA14_54890 | | 5.5746 | 3.04207 |
| PA14_54900 | | 6.16776 | 4.34312 |
| PA14_54910 | | 5.1681 | 4.79508 |
| PA14_54920 | | 5.83614 | 3.18272 |
| PA14_54930 | | 6.26179 | 3.3937 |
| PA14_54940 | | 5.34326 | 4.72273 |
| PA14_54950 | | 5.74861 | 5.24136 |

| | | | |
|---|---|---|---|
| PA14_54960 | ybtQ | 7.2675 | 6.03099 |
| PA14_54970 | | 8.38529 | 7.4164 |
| PA14_54980 | | 9.20662 | 8.60624 |
| PA14_55000 | | 7.85448 | 6.98714 |
| PA14_55020 | | 7.6662 | 5.78484 |
| PA14_55030 | | 8.02022 | 7.31145 |
| PA14_55040 | | 7.53566 | 7.20088 |
| PA14_55050 | | 1.71717 | 1.50687 |
| PA14_55070 | | 8.25707 | 8.13527 |
| PA14_55100 | | | -0.87617 |
| PA14_55200 | | | 0.71778 |
| PA14_55400 | | 2.02686 | 1.9383 |
| PA14_55410 | | 3.73222 | 4.24526 |
| PA14_55440 | hxcR | 1.55214 | 1.28253 |
| PA14_55450 | hxcQ | | |
| PA14_55480 | hxcX | | |
| PA14_55530 | hxcW | 2.17583 | 1.41893 |
| PA14_55550 | vreI | 2.05688 | 1.14752 |
| PA14_55560 | vreA | 1.74372 | 0.72661 |
| PA14_55580 | nemO | | -0.68309 |
| PA14_55631 | | -1.96656 | |
| PA14_55637 | | -1.75885 | |
| PA14_55650 | sbcC | | -0.43649 |
| PA14_55800 | fppA | | |
| PA14_55810 | pprB | 0.41678 | 0.97557 |
| PA14_55960 | pctC | -2.04162 | -1.75267 |
| PA14_56000 | pctA | -0.54758 | -1.40295 |
| PA14_56010 | pctB | -1.47707 | -1.15497 |
| PA14_56040 | | 0.47026 | 0.71897 |
| PA14_56050 | | 0.52574 | |
| PA14_56060 | purU | | -0.49989 |
| PA14_56070 | mvaT | | 0.85175 |
| PA14_56090 | | | 0.56318 |
| PA14_56130 | | -0.55484 | |
| PA14_56160 | | | |
| PA14_56180 | | | -0.56076 |
| PA14_56190 | | | |
| PA14_56220 | | | 0.86853 |
| PA14_56260 | | | 0.51967 |
| PA14_56340 | | 0.37931 | |
| PA14_56370 | | | 0.74845 |
| PA14_56480 | | | -0.75259 |
| PA14_56510 | | | -1.26912 |
| PA14_56540 | | | 0.70361 |
| PA14_56640 | | | 1.54806 |
| PA14_56660 | | 0.4813 | 3.21094 |
| PA14_56670 | | | -2.70817 |
| PA14_56680 | feoB | | -2.34173 |

| | | | |
|---|---|---|---|
| PA14_56690 | feoA | | -1.39152 |
| PA14_56830 | | | 0.54758 |
| PA14_56840 | | 0.48534 | |
| PA14_56870 | | | 0.50367 |
| PA14_56890 | mexW | | 0.6592 |
| PA14_56980 | crcB | 0.47986 | |
| PA14_56990 | | 2.12212 | 2.35298 |
| PA14_57010 | groEL | | 0.7085 |
| PA14_57020 | groES | | 0.73941 |
| PA14_57030 | fxsA | | 0.59162 |
| PA14_57060 | | 0.55896 | 0.38765 |
| PA14_57070 | | 0.46876 | |
| PA14_57100 | ampG | 0.49213 | 0.44279 |
| PA14_57110 | | | |
| PA14_57160 | apbA | | 0.7149 |
| PA14_57170 | | 0.42026 | 0.72284 |
| PA14_57180 | | | 0.99581 |
| PA14_57220 | secA | | -0.51742 |
| PA14_57240 | | 0.5067 | |
| PA14_57460 | mraZ | | -0.58414 |
| PA14_57470 | rsmI | | -0.5296 |
| PA14_57630 | | 1.09572 | 0.76793 |
| PA14_57650 | zapE | | -0.57044 |
| PA14_57720 | cysD | -1.08628 | -1.26783 |
| PA14_57770 | hisC1 | -0.44192 | |
| PA14_57820 | | | -0.45761 |
| PA14_57890 | kdsD | -0.34428 | -0.79866 |
| PA14_57950 | raiA | -0.50515 | -0.44451 |
| PA14_57980 | | 0.84 | |
| PA14_58000 | | -0.49333 | -0.44827 |
| PA14_58130 | mreC | | |
| PA14_58230 | | -1.31947 | 1.28716 |
| PA14_58240 | | -1.22045 | 1.07793 |
| PA14_58250 | magD | -1.72112 | 1.26963 |
| PA14_58260 | | -1.65009 | 1.22589 |
| PA14_58270 | | -1.61412 | 1.37208 |
| PA14_58290 | | -0.78993 | 0.44058 |
| PA14_58330 | | | 1.25353 |
| PA14_58350 | dppA1 | 1.14078 | 1.34909 |
| PA14_58360 | dppA2 | 0.69298 | 0.99328 |
| PA14_58390 | dppA3 | 4.19971 | 3.09079 |
| PA14_58410 | opdP | 2.14603 | |
| PA14_58420 | dppA4 | 1.3445 | 0.89584 |
| PA14_58540 | | 0.53318 | |
| PA14_58570 | piuA | -0.45187 | |
| PA14_58610 | | 0.66072 | |
| PA14_58630 | | 0.61824 | 0.81223 |
| PA14_58650 | | | -0.5355 |

| | | | |
|---|---|---|---|
| PA14_58690 | | -0.55894 | -1.67834 |
| PA14_58730 | pilA | 5.31459 | 5.06766 |
| PA14_58760 | pilC | 1.33751 | 0.83561 |
| PA14_58770 | pilD | | -0.61254 |
| PA14_58820 | | | -0.66913 |
| PA14_58830 | | -0.87891 | -1.10866 |
| PA14_58880 | zapE | -1.11873 | -1.32198 |
| PA14_58910 | | 1.77788 | 1.60791 |
| PA14_58920 | | 1.66426 | 2.15448 |
| PA14_58930 | | | 0.80132 |
| PA14_59050 | NdpA2 | 3.50587 | 4.57746 |
| PA14_59070 | | 5.09858 | 2.6201 |
| PA14_59090 | | 3.86512 | 3.71514 |
| PA14_59100 | | 3.93272 | 3.83805 |
| PA14_59120 | | 3.84867 | 3.58672 |
| PA14_59130 | | 6.23374 | 6.27713 |
| PA14_59180 | topA | 2.12287 | 2.1633 |
| PA14_59190 | | 3.55883 | 3.89812 |
| PA14_59200 | | 2.20039 | 2.06582 |
| PA14_59210 | | 2.02835 | 1.74308 |
| PA14_59240 | pilL2 | 1.54864 | 0.81013 |
| PA14_59250 | pilN2 | 0.97228 | |
| PA14_59270 | pilO2 | 3.6704 | 3.61065 |
| PA14_59290 | pilQ2 | 3.88621 | 3.81887 |
| PA14_59310 | pilR2 | 4.54121 | 4.13214 |
| PA14_59320 | pilS2 | 1.5027 | 2.21191 |
| PA14_59340 | pilT2 | 3.91683 | 3.98538 |
| PA14_59350 | pilV2 | 1.88291 | 1.60695 |
| PA14_59360 | pilM2 | 0.66739 | 0.94089 |
| PA14_59380 | | 8.30211 | 8.48315 |
| PA14_59400 | | 6.78109 | 6.34746 |
| PA14_59430 | | 3.93312 | 4.90913 |
| PA14_59440 | | 4.69784 | 4.6664 |
| PA14_59470 | | 6.33448 | 6.13648 |
| PA14_59480 | | 5.86889 | 3.99149 |
| PA14_59490 | vgrG4b | | 4.44545 |
| PA14_59500 | | 5.11023 | 5.01396 |
| PA14_59510 | | 6.80032 | 6.52328 |
| PA14_59520 | | 5.37354 | 3.11239 |
| PA14_59530 | | 5.17592 | 4.52363 |
| PA14_59540 | | 1.50466 | 0.55313 |
| PA14_59550 | | 4.38122 | 4.47932 |
| PA14_59580 | | 7.05704 | 7.1751 |
| PA14_59590 | | 3.14008 | 3.24166 |
| PA14_59610 | | 4.96681 | 4.93457 |
| PA14_59620 | | 6.0347 | 4.87058 |
| PA14_59630 | | 3.61719 | 3.92376 |
| PA14_59690 | traD | 2.50196 | 2.64354 |

| | | | |
|---|---|---|---|
| PA14_59710 | cupD1 | 4.23909 | 4.88483 |
| PA14_59720 | cupD2 | 1.52506 | 0.92005 |
| PA14_59735 | cupD3 | 1.15532 | 1.30804 |
| PA14_59750 | cupD4 | 1.4174 | 1.06655 |
| PA14_59760 | cupD5 | 2.31226 | 2.18235 |
| PA14_59770 | rcsB | 5.48886 | 5.13624 |
| PA14_59780 | rcsC | 5.64386 | 4.83891 |
| PA14_59790 | pvrR | 6.09351 | 5.18705 |
| PA14_59800 | pvrS | 5.58787 | 2.86625 |
| PA14_59820 | | 3.93833 | 4.15226 |
| PA14_59830 | | 3.81156 | 3.64841 |
| PA14_59845 | | 5.27155 | 5.29946 |
| PA14_59900 | | 1.82894 | 1.34094 |
| PA14_59910 | | 4.12341 | 3.80321 |
| PA14_59920 | | 1.04348 | 1.01118 |
| PA14_59930 | | 3.26799 | 4.00281 |
| PA14_59940 | | 3.25855 | 3.60993 |
| PA14_59960 | dsbA2 | -1.54297 | -1.37139 |
| PA14_59970 | | 4.84959 | 4.67475 |
| PA14_59990 | | 1.35054 | |
| PA14_60000 | | 3.40219 | 3.09712 |
| PA14_60020 | | 2.42619 | 1.69002 |
| PA14_60030 | | 7.07431 | 6.96244 |
| PA14_60040 | | 6.91517 | 7.06694 |
| PA14_60050 | | | 4.30636 |
| PA14_60070 | | 4.67683 | 3.74534 |
| PA14_60080 | | 7.4586 | 8.30604 |
| PA14_60090 | | 4.14409 | 4.10866 |
| PA14_60100 | | 4.60052 | 4.69184 |
| PA14_60110 | | | 3.85083 |
| PA14_60120 | dcd2 | 8.63599 | 8.34629 |
| PA14_60130 | | 4.62558 | 3.96605 |
| PA14_60140 | | 4.07067 | 3.8182 |
| PA14_60200 | pgeF | | -0.54038 |
| PA14_60210 | rluD | | -0.56517 |
| PA14_60270 | thiO | 1.95766 | 1.38922 |
| PA14_60280 | fimU | 3.48599 | 3.65907 |
| PA14_60290 | pilW | 4.09445 | 4.00747 |
| PA14_60300 | pilX | 4.84821 | 4.78395 |
| PA14_60310 | pilY1 | 5.29447 | 3.25305 |
| PA14_60320 | pilE | 4.72786 | 4.70922 |
| PA14_60370 | ileS | -0.3947 | |
| PA14_60380 | ribF | | -0.49871 |
| PA14_60445 | obgW | | -0.61509 |
| PA14_60460 | rplU | | -0.74943 |
| PA14_60480 | | -1.68901 | -1.61073 |
| PA14_60490 | | 0.43919 | |
| PA14_60530 | | | -0.56629 |

| | | | |
|---|---|---|---|
| PA14_60540 | | -0.39708 | -0.81628 |
| PA14_60560 | | | 0.86322 |
| PA14_60570 | | | 1.04793 |
| PA14_60580 | | | 0.90594 |
| PA14_60630 | | -2.25127 | -2.59446 |
| PA14_60650 | | -2.91996 | -3.17665 |
| PA14_60660 | | -2.45949 | -2.51715 |
| PA14_60670 | rtcA | -2.26687 | -1.79514 |
| PA14_60730 | | 2.18261 | 1.87922 |
| PA14_60750 | (pra) | 2.72794 | 2.88153 |
| PA14_60760 | | 0.97083 | 1.15632 |
| PA14_60770 | | 0.76463 | 0.9295 |
| PA14_60780 | | 1.1387 | 0.54231 |
| PA14_60790 | | 1.03698 | |
| PA14_60810 | escR | | 1.33372 |
| PA14_60850 | mexC | 1.15163 | 0.79714 |
| PA14_60870 | morA | | |
| PA14_60890 | glyA | | -0.58559 |
| PA14_60900 | | 1.3704 | 0.98859 |
| PA14_60920 | yjiA | 1.03992 | 0.71006 |
| PA14_60950 | | | 0.46458 |
| PA14_60960 | | | 1.75307 |
| PA14_61000 | | | 0.70362 |
| PA14_61050 | mscL | | 0.83127 |
| PA14_61090 | | -0.56862 | -0.67411 |
| PA14_61130 | | | -0.94735 |
| PA14_61140 | | -0.48988 | -1.01938 |
| PA14_61150 | | | -0.6268 |
| PA14_61170 | | | -0.64099 |
| PA14_61190 | cdrB | 2.75495 | 3.55244 |
| PA14_61200 | | 3.70665 | 4.65121 |
| PA14_61270 | | -1.15812 | |
| PA14_61290 | | | 0.58736 |
| PA14_61300 | | -0.85131 | -0.83724 |
| PA14_61370 | | -1.18112 | -1.77444 |
| PA14_61410 | | 0.75635 | 1.05279 |
| PA14_61440 | | -0.70418 | -0.99042 |
| PA14_61500 | | 0.92671 | 2.96687 |
| PA14_61510 | | | 1.04198 |
| PA14_61520 | | | 1.75721 |
| PA14_61530 | | | 0.84538 |
| PA14_61550 | | | 0.93134 |
| PA14_61650 | pagL | | 0.55884 |
| PA14_61710 | hemA | 0.43674 | |
| PA14_61780 | | 0.38769 | 0.48604 |
| PA14_61840 | | -4.27612 | -8.11652 |
| PA14_61850 | | 0.62743 | 0.83675 |
| PA14_61860 | can | | -1.23667 |

| | | | |
|---|---|---|---|
| PA14_61870 | | | -1.41286 |
| PA14_61950 | | -1.67649 | -0.72567 |
| PA14_61960 | mksF | | -0.51304 |
| PA14_62000 | (hitA) | -0.73268 | -0.592 |
| PA14_62020 | | 0.69354 | 0.54118 |
| PA14_62060 | | | 2.58296 |
| PA14_62090 | | -2.26837 | |
| PA14_62150 | ilvH | -0.43254 | -0.54231 |
| PA14_62160 | ilvI | | -0.45176 |
| PA14_62250 | | | 0.79909 |
| PA14_62300 | | | -0.48054 |
| PA14_62330 | | | |
| PA14_62390 | | 2.8244 | 2.52278 |
| PA14_62430 | | | -0.80111 |
| PA14_62440 | | 0.42661 | |
| PA14_62510 | gluQRS | 0.76461 | |
| PA14_62620 | pgi | -0.30886 | -0.36139 |
| PA14_62680 | | 0.78023 | |
| PA14_62690 | | 2.06284 | 1.92058 |
| PA14_62730 | truB | 0.48332 | |
| PA14_62780 | rimP | | |
| PA14_62810 | secG | | -0.68602 |
| PA14_62840 | glmM | 0.55236 | 0.46791 |
| PA14_62930 | carA | | -0.58291 |
| PA14_62960 | dnaJ | | 0.85785 |
| PA14_62970 | dnaK | | 0.56305 |
| PA14_62990 | grpE | | 1.0782 |
| PA14_63010 | recN | | -0.56314 |
| PA14_63030 | omlA | -0.59323 | -0.94452 |
| PA14_63170 | cueR | 0.48901 | |
| PA14_63230 | | -0.39494 | -0.84861 |
| PA14_63240 | yedA | | |
| PA14_63340 | | -0.75342 | -0.521 |
| PA14_63390 | | -1.43942 | -1.02886 |
| PA14_63410 | | 1.17437 | 1.46993 |
| PA14_63420 | | 0.99466 | 1.63182 |
| PA14_63430 | | 0.87152 | 1.38271 |
| PA14_63450 | | 6.79417 | 7.03156 |
| PA14_63460 | | 5.06412 | 4.6354 |
| PA14_63470 | | | 0.50905 |
| PA14_63480 | | -2.23954 | -2.10733 |
| PA14_63605 | fdnG | 0.38531 | 0.66048 |
| PA14_63640 | fadH2 | 0.56757 | |
| PA14_63860 | | -2.36728 | -2.31608 |
| PA14_63900 | | -0.64456 | |
| PA14_64000 | | | 0.59275 |
| PA14_64050 | gcbA | -0.44423 | -0.69308 |
| PA14_64060 | ctpL | -0.58186 | -1.16816 |

| | | | |
|---|---|---|---|
| PA14_64090 | aroQ1 | 0.63099 | 0.62857 |
| PA14_64110 | accC | | 0.56494 |
| PA14_64120 | | | |
| PA14_64140 | prmA | 0.72875 | |
| PA14_64170 | | | -0.65793 |
| PA14_64180 | dusB | | -0.52274 |
| PA14_64230 | retS | -0.4112 | |
| PA14_64335 | ureD | 0.48859 | |
| PA14_64370 | ureB | | 0.64912 |
| PA14_64390 | ureC | 0.54395 | 1.05013 |
| PA14_64400 | | | -0.89302 |
| PA14_64410 | | 1.02841 | 0.86747 |
| PA14_64430 | | 4.82266 | 4.02665 |
| PA14_64440 | | 0.36181 | |
| PA14_64460 | | | 0.57097 |
| PA14_64530 | | 1.72182 | 3.76747 |
| PA14_64550 | | | 0.89327 |
| PA14_64560 | | | 1.20151 |
| PA14_64620 | | | -0.85317 |
| PA14_64850 | | 0.46969 | 0.75205 |
| PA14_64860 | | 0.68358 | 0.70694 |
| PA14_64870 | | 1.13681 | 1.32978 |
| PA14_64880 | livM | 0.89876 | 1.13923 |
| PA14_64890 | | 1.15356 | 0.90157 |
| PA14_64900 | | 0.94102 | 0.9804 |
| PA14_64920 | | 0.42361 | 0.92732 |
| PA14_64930 | | 0.76852 | 0.70291 |
| PA14_64940 | | | 0.63324 |
| PA14_64950 | pncA | -2.00099 | -2.59145 |
| PA14_64960 | pncB1 | -2.1235 | -1.43037 |
| PA14_64980 | nadE | -1.24273 | -1.05692 |
| PA14_64990 | choE | | -0.63829 |
| PA14_65000 | azu | | 0.76788 |
| PA14_65030 | | 0.47803 | |
| PA14_65040 | | | 0.79171 |
| PA14_65090 | | 1.48294 | 1.01885 |
| PA14_65170 | rpsR | -0.95452 | -1.01772 |
| PA14_65180 | rpsF | -0.85125 | -0.97012 |
| PA14_65230 | purA | 0.84059 | 0.6766 |
| PA14_65250 | hisZ | 1.10865 | 0.64639 |
| PA14_65320 | miaA | -0.42091 | -0.75031 |
| PA14_65400 | queG | | -0.47356 |
| PA14_65470 | | 0.64334 | |
| PA14_65560 | serB | | -0.56428 |
| PA14_65590 | | 0.50909 | |
| PA14_65740 | thiC | -0.52078 | -2.63481 |
| PA14_65750 | opmH | 0.41108 | |
| PA14_65860 | | 2.20036 | 1.78018 |

| | | | |
|---|---|---|---|
| PA14_65920 | | | |
| PA14_66040 | | | -0.50914 |
| PA14_66060 | rfaE | | -0.46909 |
| PA14_66080 | msbA | | -0.74511 |
| PA14_66100 | | 3.23402 | 2.9374 |
| PA14_66140 | dnpA | 0.54002 | |
| PA14_66190 | | 0.56864 | 0.85462 |
| PA14_66200 | | 0.80731 | 0.88366 |
| PA14_66210 | | 0.84267 | 0.99726 |
| PA14_66220 | waaP | 1.23534 | 1.56085 |
| PA14_66230 | waaG | 1.02853 | 1.23359 |
| PA14_66240 | waaC | 1.50725 | 1.36004 |
| PA14_66250 | waaF | 1.23846 | 0.99203 |
| PA14_66290 | aceE | | |
| PA14_66340 | | | -0.42727 |
| PA14_66420 | | -0.88743 | -1.16554 |
| PA14_66440 | metY | -0.87061 | |
| PA14_66450 | | | |
| PA14_66460 | | 0.73126 | 2.33639 |
| PA14_66480 | | -0.39577 | -0.57709 |
| PA14_66510 | | -0.58555 | |
| PA14_66530 | | | -0.82437 |
| PA14_66600 | aroB | -0.351 | |
| PA14_66620 | pilQ | 0.69068 | 0.60166 |
| PA14_66630 | pilP | 0.97618 | 0.77859 |
| PA14_66640 | pilO | 0.84801 | 0.74166 |
| PA14_66690 | | 0.7232 | 0.4941 |
| PA14_66700 | | 0.43518 | |
| PA14_66710 | | -0.40592 | -0.65354 |
| PA14_66760 | | | 0.47395 |
| PA14_66770 | hslV | | 1.05535 |
| PA14_66790 | hslU | | 0.83064 |
| PA14_66820 | phaC1 | 0.71562 | |
| PA14_66830 | phaD | 0.8162 | 0.54341 |
| PA14_66840 | phaC2 | 1.38029 | 2.12408 |
| PA14_66850 | | 0.74196 | 1.57361 |
| PA14_66950 | hisE | -0.48035 | -0.67922 |
| PA14_67180 | | | 2.34804 |
| PA14_67190 | tli5b3 | | 2.85842 |
| PA14_67200 | tli5b2 | 1.4265 | 4.32441 |
| PA14_67210 | | 3.5464 | 6.2068 |
| PA14_67240 | hutG | | 0.49724 |
| PA14_67250 | hutI | 0.38119 | 0.76538 |
| PA14_67260 | | | 0.55723 |
| PA14_67280 | | 1.24619 | 1.35858 |
| PA14_67340 | | 1.3845 | 0.69296 |
| PA14_67380 | | -1.65146 | -1.91357 |
| PA14_67400 | | -1.37237 | -1.40758 |

| | | | |
|---|---|---|---|
| PA14_67440 | | 0.91501 | |
| PA14_67510 | estA | | 1.38309 |
| PA14_67520 | | -1.05461 | 1.62067 |
| PA14_67530 | | | 2.42945 |
| PA14_67540 | | 1.8609 | 2.36439 |
| PA14_67620 | | | |
| PA14_67670 | ntrB | | |
| PA14_67780 | | -0.47063 | -0.86307 |
| PA14_67830 | | | 1.23169 |
| PA14_67850 | | -0.78418 | -0.93253 |
| PA14_68030 | | -0.3996 | |
| PA14_68070 | | 0.76622 | 0.63189 |
| PA14_68190 | rmlD | | 0.4141 |
| PA14_68200 | rmlA | | 0.69527 |
| PA14_68210 | rmlC | | |
| PA14_68260 | dctP | 0.99599 | |
| PA14_68280 | dctQ | 1.12413 | |
| PA14_68290 | dctM | 1.75998 | |
| PA14_68300 | arcD | | 1.14222 |
| PA14_68330 | arcA | 0.45167 | 0.93374 |
| PA14_68350 | arcC | 0.61068 | 0.62417 |
| PA14_68360 | fabY | 0.53656 | 0.77888 |
| PA14_68380 | nudE | | -0.47355 |
| PA14_68390 | yrfG | | -0.51959 |
| PA14_68400 | lysM | | 0.52519 |
| PA14_68430 | fdhD | | 2.27276 |
| PA14_68440 | | | 3.28542 |
| PA14_68460 | | 2.12297 | 0.72845 |
| PA14_68470 | | | 1.29231 |
| PA14_68480 | | | -0.6155 |
| PA14_68490 | | -0.76129 | -0.78316 |
| PA14_68510 | | -0.66728 | -0.91518 |
| PA14_68530 | | | -0.7287 |
| PA14_68630 | | -0.84376 | -1.12335 |
| PA14_68670 | | -0.42666 | -0.45079 |
| PA14_68800 | | -0.72721 | |
| PA14_68810 | | | |
| PA14_68830 | | | 0.71231 |
| PA14_68850 | gcvP1 | | 0.43873 |
| PA14_68930 | | 0.54696 | 0.6895 |
| PA14_68940 | | 0.92024 | 1.63541 |
| PA14_68970 | | 0.44885 | |
| PA14_69030 | | -0.4132 | -0.50699 |
| PA14_69060 | | 0.9908 | 1.02638 |
| PA14_69070 | rbbA | 1.18181 | 1.55912 |
| PA14_69090 | | | 0.79583 |
| PA14_69190 | rho | | |
| PA14_69240 | hemB | | -0.55647 |

| | | | |
|---|---|---|---|
| PA14_69300 | | | -0.44534 |
| PA14_69320 | | 1.14536 | 1.11973 |
| PA14_69370 | algP | | 0.5276 |
| PA14_69450 | hemC | -0.43845 | -0.45428 |
| PA14_69500 | argH | -0.74973 | -0.97698 |
| PA14_69510 | | 6.00621 | 7.18469 |
| PA14_69520 | | 5.10138 | 8.07222 |
| PA14_69540 | ccoN | | 5.03974 |
| PA14_69550 | | 0.64003 | 5.71451 |
| PA14_69580 | | -1.01333 | -0.68182 |
| PA14_69620 | | | 0.44994 |
| PA14_69630 | rnk | 0.62197 | |
| PA14_69720 | | | 0.5471 |
| PA14_69810 | glnK | -1.18976 | -0.88913 |
| PA14_69840 | | | 1.60545 |
| PA14_69850 | betT | -0.3613 | |
| PA14_69900 | | 1.18217 | |
| PA14_69950 | | 0.55057 | 0.63113 |
| PA14_69970 | cycB | 0.46808 | |
| PA14_69980 | | -0.43326 | -0.44131 |
| PA14_69990 | dadX | 0.64111 | 0.61664 |
| PA14_70040 | dadA | -0.55363 | |
| PA14_70050 | | | 0.64572 |
| PA14_70120 | | 0.5237 | |
| PA14_70140 | | | -0.51862 |
| PA14_70160 | | -0.50547 | -1.31227 |
| PA14_70170 | | | -0.91363 |
| PA14_70180 | rpmG | | -0.71904 |
| PA14_70190 | rpmB | | -0.74063 |
| PA14_70230 | radC | -0.49846 | |
| PA14_70370 | pyrE | -0.57076 | -0.89484 |
| PA14_70390 | crc | -0.46115 | -0.78956 |
| PA14_70450 | rpoZ | -0.63451 | -0.91036 |
| PA14_70480 | | -0.43398 | -0.70637 |
| PA14_70690 | glcD | -0.59062 | -0.81745 |
| PA14_70710 | glcC | 0.37721 | 0.98025 |
| PA14_70800 | phoU | | 0.60342 |
| PA14_70810 | pstB | | |
| PA14_70850 | pstC | | 0.71294 |
| PA14_70860 | pstS | | |
| PA14_70880 | | -0.89329 | |
| PA14_70910 | | -2.6358 | |
| PA14_70950 | betB | -0.44978 | |
| PA14_70970 | betI | -0.48456 | -0.60635 |
| PA14_71000 | choV | | 0.4778 |
| PA14_71090 | | | 0.64943 |
| PA14_71100 | | -0.96447 | |
| PA14_71220 | cls | 0.97154 | 0.61926 |

| | | | |
|---|---|---|---|
| PA14_71230 | | | |
| PA14_71410 | gbcA | -2.30126 | -2.64625 |
| PA14_71460 | glyA1 | -4.40835 | -4.41428 |
| PA14_71620 | purE | 0.61126 | 0.53724 |
| PA14_71630 | adhA | | 0.67738 |
| PA14_71750 | pycR | | -0.46471 |
| PA14_71830 | | -0.82011 | |
| PA14_71840 | | -0.48985 | |
| PA14_71920 | wbpY | 0.74452 | 0.63963 |
| PA14_71930 | wbpX | 0.77116 | 0.84666 |
| PA14_71960 | wzm | 0.73881 | |
| PA14_71990 | gmd | 1.12289 | 0.53314 |
| PA14_72000 | rmd | 0.56036 | |
| PA14_72010 | | 0.5666 | |
| PA14_72040 | | 0.60527 | |
| PA14_72080 | | -0.54213 | -0.75794 |
| PA14_72090 | | -0.6778 | -0.70023 |
| PA14_72110 | | | -0.50229 |
| PA14_72150 | | | 0.58657 |
| PA14_72170 | | | -0.68784 |
| PA14_72280 | (citA) | | |
| PA14_72340 | gltP | 1.02014 | |
| PA14_72350 | | 2.68362 | 1.75713 |
| PA14_72360 | | 1.66367 | |
| PA14_72410 | | | -0.71655 |
| PA14_72450 | dsbA1 | -0.48452 | -0.72951 |
| PA14_72480 | engB | -0.50167 | -0.54836 |
| PA14_72500 | | -1.07174 | -0.47464 |
| PA14_72550 | | -0.3811 | -0.4515 |
| PA14_72560 | np20 | -0.38322 | |
| PA14_72630 | | | -0.75895 |
| PA14_72650 | qapR | -0.52705 | -1.03127 |
| PA14_72760 | | 0.97316 | 0.75713 |
| PA14_72770 | | | -0.57873 |
| PA14_72800 | | | -0.39837 |
| PA14_72810 | | -0.45664 | -0.5394 |
| PA14_72840 | | -0.43723 | |
| PA14_72850 | | -1.47291 | -1.49182 |
| PA14_72870 | | -2.51183 | -2.61962 |
| PA14_72880 | | -1.34072 | -1.15309 |
| PA14_72930 | | 1.39369 | 1.26854 |
| PA14_72940 | | -0.87447 | -0.74831 |
| PA14_72960 | | | -1.72978 |
| PA14_72990 | | -0.50858 | -0.75659 |
| PA14_73060 | | | -0.59747 |
| PA14_73150 | | -0.90394 | -0.67237 |
| PA14_73160 | | -0.8187 | -1.41427 |
| PA14_73230 | atpC | | 0.58343 |

| | | | |
|---|---|---|---|
| PA14_73240 | atpD | | 0.74527 |
| PA14_73250 | atpG | | 0.54249 |
| PA14_73330 | parB | 0.54835 | 0.77348 |
| PA14_73350 | soj | 0.93307 | 1.1848 |
| PA14_73390 | trmE | 0.69083 | 0.54871 |

| *rsmA* vs *nahK* ($log_2$FC ) |
|---:|
| 1.77389 |
| 0.16238 |
| 0.7178 |
| 0.62589 |
| 1.88396 |
| 3.21153 |
| -1.79702 |
| 2.26211 |
| 2.02215 |
| 1.96703 |
| 1.71351 |
| 0.42953 |
| 2.36173 |
| 1.24251 |
| 1.49325 |
| 1.7153 |
| 2.19308 |
| 2.24703 |
| 2.21132 |
| 2.09561 |
| 2.4172 |
| 2.32803 |
| 2.35632 |
| 1.2232 |
| 2.69009 |
| 2.88118 |
| 3.86391 |
| 2.73281 |
| 3.24957 |
| 3.36229 |
| 2.99475 |
| 3.301 |
| 3.3652 |
| 1.40217 |

| |
|---|
| 0.68609 |
| 2.37816 |
| 2.43681 |
| 1.28242 |
| 3.13099 |
| 3.50065 |
| 3.34165 |
| 3.2826 |
| 2.77911 |
| 2.84984 |
| -0.1448 |
| 1.42381 |
| -0.29776 |
| -0.63322 |
| -0.16985 |
| -1.01363 |
| -1.02036 |
| 0.25938 |
| 0.20634 |
| 0.32363 |
| 0.16936 |
| 0.16002 |
| 0.24752 |
| 1.50003 |
| 0.67726 |
| 0.42887 |

| |
|---|
| 0.20677 |
| 4.23424 |
| 4.58184 |
| 6.49151 |
| 0.82889 |
| 0.20862 |
| 1.29722 |
| 1.29414 |
| 0.85559 |
| 0.16058 |
| 0.64589 |
| 0.38341 |
| -0.22942 |
| 1.14798 |
| 0.7083 |
| 0.7143 |
| -0.3417 |
| 1.27329 |
| 0.12123 |

| |
|---|
| 0.25745 |
| -0.24767 |
| -0.21857 |
| 0.49216 |
| -0.18606 |
| 0.5445 |
| 0.91926 |
| 0.66181 |
| -0.39309 |
| -0.53448 |
| -0.67309 |
| -0.3561 |
| -0.19856 |

| |
|---|
| -0.6326 |
| 0.3128 |
| -0.27667 |
| 0.29604 |
| 2.72802 |
| 0.21145 |
| 0.20985 |
| 0.17578 |
| 0.49659 |
| 0.96456 |
| 0.24214 |
| 0.88888 |
| 0.76243 |
| 0.36826 |
| 0.17993 |
| 0.27816 |
| 1.10453 |

| |
|---|
| 1.79498 |
| 0.81068 |
| -0.466 |
| 3.66434 |
| 3.74401 |
| 1.03777 |
| 1.05751 |
| 1.41106 |
| 3.72116 |
| 3.62734 |
| 1.24541 |
| 1.23474 |
| 1.5823 |
| 1.61538 |
| 0.18085 |

| |
|---|
| -0.43361 |
| -0.42017 |
| -0.37534 |
| -0.19589 |
| -0.18742 |
| 1.84989 |
| 1.80051 |
| 2.17853 |
| 2.30279 |
| -0.78119 |
| -1.00598 |
| -0.22777 |
| 1.44798 |
| 0.59283 |

| |
|---|
| -0.30627 |
| -0.27091 |
| 0.41416 |
| -0.31398 |
| -0.28781 |
| 0.34749 |
| 1.33324 |
| 0.78167 |
| 1.6135 |
| -0.44524 |

| |
|---|
| 0.23169 |
| 2.57777 |
| 2.16869 |
| 1.80644 |
| 0.12162 |
| 0.79476 |
| 1.44738 |
| -0.21558 |
| -0.16696 |
| -0.41544 |
| -0.73465 |
| 0.18465 |
| -1.17907 |
| 1.96935 |
| -1.65359 |
| -1.29108 |
| -0.15623 |
| -0.21102 |

| |
|---|
| -0.94279 |
| -0.80266 |
| -1.04976 |
| -0.44502 |
| 0.49654 |
| 2.82997 |
| 2.68441 |
| 3.53148 |
| 4.11632 |
| 4.0879 |
| 4.71393 |
| 1.42526 |

| |
|---|
| 0.441 |
| 0.09952 |
| 1.42131 |
| 0.74417 |
| 2.21756 |
| 0.59062 |
| 2.39856 |
| 0.26192 |
| -0.19856 |
| -0.90775 |
| -0.29708 |
| -0.34028 |

| |
|---|
| -0.10702 |
| -0.22048 |
| -0.24952 |
| -0.65444 |
| -0.64094 |
| -0.20194 |
| 4.60418 |
| 2.54449 |
| 3.27679 |
| 0.17201 |
| -0.37794 |
| -0.13518 |

| |
|---|
| 0.17348 |
| 0.22549 |
| 0.65945 |
| 0.30214 |
| 0.62822 |
| 0.27611 |
| 0.44758 |
| -0.27854 |
| -0.5185 |
| -0.63564 |
| 0.49711 |
| -0.41438 |
| -0.51063 |
| -0.75693 |
| 0.68999 |
| 0.32835 |
| 3.18464 |
| 0.37404 |
| 2.84328 |
| 3.11671 |
| 2.90473 |
| 3.11064 |
| 3.13711 |

| |
|---|
| 3.35915 |
| 3.32307 |
| 3.41158 |
| 3.47713 |
| 3.45587 |
| 0.12937 |
| 2.50088 |
| 1.12259 |
| 4.81995 |
| 2.54482 |
| 0.90426 |
| 0.21624 |
| 0.1145 |
| -0.21534 |
| -0.17727 |
| -0.17925 |
| 1.54806 |

| |
|---|
| 0.11788 |
| -0.4651 |
| 0.14227 |
| 0.51073 |
| -0.0884 |

| |
|---|
| -0.16686 |
| -0.18611 |
| -0.36412 |
| -1.59666 |
| -0.52274 |
| 0.11589 |
| 1.11563 |
| 0.4114 |
| 0.75103 |
| 0.26156 |
| 0.30956 |
| 0.9093 |
| 0.20687 |
| 0.18433 |
| 0.20576 |
| 3.48349 |
| -0.20463 |
| -0.43723 |
| -0.31609 |

| |
|---|
| 1.32192 |
| -0.63576 |
| -0.44096 |
| -0.17361 |
| -0.30429 |
| -0.24753 |
| -0.16112 |
| 1.12666 |
| 0.14709 |
| -0.35241 |
| 1.9747 |
| 2.22971 |
| 2.18488 |
| 1.60524 |
| 2.0381 |
| 1.80866 |
| 2.50598 |
| 0.49203 |
| 0.09879 |
| -0.27814 |
| -0.13943 |

| |
|---|
| 1.67916 |
| 2.88934 |
| 0.15366 |
| 1.80587 |
| 2.15116 |
| 1.35038 |
| 2.28877 |
| 1.94592 |
| 0.28923 |
| 2.03786 |
| -0.08775 |
| 1.13684 |
| 0.21268 |
| 0.27402 |
| -0.21359 |
| -0.27722 |
| -0.09163 |
| 0.25824 |
| 1.84556 |
| 1.69063 |
| 1.81572 |
| 1.433 |
| 1.84621 |
| 0.40051 |

| |
|---|
| 0.46696 |
| 3.64259 |
| 0.15312 |
| 0.26784 |
| 0.46119 |
| 4.40326 |
| 1.49859 |
| 2.36761 |
| 0.97209 |
| 0.15023 |
| -0.15195 |
| 0.15213 |
| 0.48723 |
| 0.25354 |
| 0.37214 |
| 0.9342 |

| |
|---|
| 1.18912 |
| 0.69668 |
| 0.94832 |
| 1.96116 |
| 0.76422 |
| 3.58205 |
| 4.55505 |
| -0.80236 |
| 3.02999 |
| 0.51809 |
| -1.10451 |
| 0.80013 |

| |
|---|
| 0.18464 |
| 1.74599 |
| 4.7503 |
| 4.39285 |
| 4.02029 |
| 2.98268 |
| 3.10563 |
| 2.70269 |
| -0.3742 |
| -0.38046 |
| 0.27981 |
| 1.19498 |
| 0.2217 |
| 0.45315 |
| 1.18899 |
| 1.5272 |
| 0.9916 |
| 2.18438 |
| 0.45442 |
| 1.54024 |
| 2.15367 |
| 1.35666 |
| 1.41283 |
| 1.78766 |
| 1.00835 |
| 0.1586 |

| |
|---|
| 1.92506 |
| 0.14526 |
| 1.55192 |
| 1.87485 |
| 1.51435 |
| 1.48133 |
| 1.50369 |
| 0.85053 |
| 0.16691 |
| 0.37489 |
| 0.17837 |
| 0.29184 |
| 0.17638 |
| 0.39235 |
| 0.57539 |
| 1.67678 |
| 2.05978 |
| 2.33731 |
| 2.20122 |
| 2.29702 |
| 2.52629 |
| 2.59556 |
| 2.81615 |
| 3.0219 |
| 2.70612 |
| 1.13434 |
| 1.89488 |
| 2.03387 |
| 1.93091 |

| |
|---|
| 1.60578 |
| 1.34034 |
| 1.44828 |
| 1.7407 |
| 1.28611 |
| 1.27456 |
| 0.29954 |
| 1.48637 |
| 1.96139 |
| 2.00155 |
| 1.69897 |
| 1.0738 |
| 1.487 |
| 0.31177 |
| 0.15686 |
| 0.26412 |
| 0.24284 |
| -0.12941 |
| 0.53505 |
| 0.28205 |
| 0.15724 |

| |
|---|
| 1.0252 |
| 1.9604 |
| 2.20985 |
| 2.58198 |
| 2.96239 |
| 1.32081 |
| 1.74504 |
| 0.65492 |
| 1.43308 |
| 0.18494 |
| 1.12562 |
| 1.54322 |
| 1.56865 |
| 0.90472 |
| 1.58946 |
| 1.73888 |
| 0.74519 |

| |
|---|
| 2.46778 |
| 2.32353 |
| 2.83512 |
| 3.21931 |
| 3.23847 |
| 3.24347 |
| 3.45734 |
| 2.67669 |
| -0.27324 |
| 0.27113 |
| 1.17721 |
| 0.67315 |
| 1.56846 |
| 0.27924 |
| 0.44217 |
| 0.22857 |
| 0.38035 |

| |
|---|
| 0.93387 |
| -0.44204 |
| -0.16095 |
| 3.39852 |
| 3.20362 |
| 0.89893 |
| 0.86658 |
| -0.15231 |
| 1.10266 |
| -0.20726 |
| -0.20094 |
| 0.32418 |
| 1.50588 |
| 0.36969 |
| 0.4046 |
| 4.92799 |
| 1.0029 |
| 1.39799 |

| |
|---|
| 1.75542 |
| 1.33823 |
| 0.47749 |
| 1.56355 |
| 0.90445 |
| 0.61474 |
| 1.30534 |
| 0.29702 |
| 2.0095 |
| 1.47148 |
| 0.1893 |
| 0.61466 |
| 1.83068 |
| 1.2232 |
| 2.8233 |
| 0.53458 |
| 0.91359 |
| -0.37587 |
| 0.16615 |
| 0.13645 |
| 0.56362 |
| 1.09707 |
| 0.24229 |
| -0.76579 |

| |
|---|
| 0.12882 |
| -0.33352 |
| -0.237 |
| -0.55132 |
| -1.08497 |
| -0.89228 |
| -1.27705 |
| -1.02897 |
| -0.31224 |
| -2.13335 |
| -1.60398 |
| -1.85384 |
| -3.18776 |
| -3.43408 |
| -2.79356 |
| -3.0175 |
| -0.58549 |
| -0.4341 |
| -0.6539 |
| -0.94741 |
| -0.68837 |
| -0.54338 |
| -0.19466 |
| 0.81932 |
| 4.42753 |
| 1.02012 |
| 5.19604 |
| 5.4499 |
| 5.6767 |
| 5.43381 |
| 3.32377 |
| 4.71908 |
| 4.90715 |
| 5.01581 |

| |
|---|
| 5.03279 |
| 0.36536 |
| 0.65701 |
| 1.25958 |
| 0.87874 |
| 0.2077 |
| 0.41361 |
| 0.30073 |
| 0.51433 |
| 0.15161 |
| -0.30225 |
| 0.16774 |
| 0.29329 |
| -0.20414 |
| -0.12772 |
| -0.22066 |
| 4.98589 |
| 1.85225 |

| |
|---|
| 1.20066 |
| -0.41028 |
| 1.20585 |
| 0.41605 |
| -0.64457 |
| -0.32159 |
| -0.44955 |
| -0.19975 |
| 0.34626 |
| 0.34159 |
| 0.2825 |
| 1.74593 |
| 0.47378 |

| |
|---|
| -0.58372 |
| -0.48912 |
| -0.25093 |
| 0.30957 |
| -0.63722 |
| -0.92829 |
| -0.25577 |
| -0.89989 |
| -0.91779 |
| 0.50338 |
| -0.26583 |
| -0.24099 |

| |
|---|
| -0.21412 |
| 0.34886 |
| 0.36971 |
| 0.24312 |
| 1.40536 |
| -0.86285 |
| -2.39101 |
| 0.22784 |
| -0.28886 |
| 1.88424 |
| 1.77046 |
| 5.31715 |
| 0.62821 |
| 0.35949 |

| |
|---|
| 0.44668 |
| 0.13082 |
| 0.2337 |
| 0.18121 |
| 0.53678 |
| 0.58583 |
| 0.26935 |
| 1.57746 |
| 1.7586 |
| 0.16682 |

| |
|---|
| 0.32497 |
| 0.42028 |
| 0.40988 |
| 0.56952 |
| 0.74001 |
| 0.50543 |
| 0.17471 |
| -0.59284 |
| -0.58489 |
| 0.36136 |
| 0.28142 |
| -0.96941 |
| -0.14992 |
| 0.16864 |
| -0.34018 |
| 0.61141 |
| 0.33266 |
| -7.40994 |

| |
|---|
| -0.22129 |
| 0.58912 |
| -0.16911 |
| 0.8996 |
| 0.36447 |
| 0.29219 |
| -0.24584 |
| 1.79045 |
| 0.41874 |
| 0.54873 |
| -0.57835 |
| 0.1539 |
| -0.62308 |

| |
|---|
| 0.78376 |
| -0.6322 |
| -0.11836 |
| 0.14744 |
| 0.20319 |
| -0.13684 |
| -0.77493 |
| -0.41851 |
| -0.26951 |
| 1.10972 |
| 1.60682 |
| -0.44751 |
| 0.92966 |
| 1.31882 |
| 0.12758 |
| 0.86085 |
| 6.04134 |

| |
|---|
| -0.8353 |
| -0.31504 |
| -0.36526 |
| -0.77143 |
| -0.17793 |
| -0.2462 |
| -0.43516 |
| 1.66138 |
| 2.02877 |
| -0.42826 |
| -0.61967 |
| 0.97005 |
| 0.45848 |
| -0.1963 |
| 0.36799 |
| 0.25069 |
| -0.38542 |
| -0.93313 |
| 0.52008 |
| 1.24101 |
| 2.27574 |
| -2.59717 |
| -2.65383 |

| |
|---|
| -1.63011 |
| 0.67774 |
| -0.12742 |
| -0.16405 |
| -0.1275 |
| 3.13236 |
| 2.70556 |
| 3.34492 |
| 3.32487 |
| 3.27684 |
| 1.5631 |
| 0.65997 |
| -0.39518 |
| -0.55809 |
| -0.77956 |

| |
|---|
| -0.7316 |
| -0.57819 |
| -0.38489 |
| 4.95535 |
| -0.21945 |

| |
|---|
| -0.3737 |
| 3.47856 |
| 1.00769 |
| 0.41466 |
| 6.0638 |

| |
|---|
| 0.68879 |
| 1.1285 |
| -0.26959 |
| -0.14448 |
| 1.47104 |
| 0.69264 |
| 0.46957 |
| 0.20119 |
| 0.2212 |
| 0.61888 |
| 1.09452 |
| -2.34576 |
| -1.01251 |

| |
|---|
| -1.33561 |
| 1.95523 |
| 0.33969 |
| -0.19463 |
| -0.12713 |
| 0.17523 |
| 0.18183 |
| -0.12235 |
| -0.16351 |

| |
|---|
| 0.21379 |
| 0.6317 |
| -0.30279 |
| 0.25454 |
| 1.52574 |
| 0.24983 |
| 0.20614 |
| 0.10293 |
| -0.08527 |
| -0.35526 |
| -0.12709 |
| -0.34639 |
| -1.59622 |

| |
|---|
| -0.2255 |
| -0.10863 |
| 0.14956 |
| 1.71067 |
| 0.3387 |
| -0.29807 |
| 0.23927 |
| 0.19961 |
| -0.12141 |
| 0.19142 |
| 0.15081 |
| 2.39955 |
| 2.15178 |
| 2.62311 |
| 3.17664 |

| |
|---|
| 1.16413 |
| 3.26263 |
| 2.73546 |
| 0.23993 |
| 0.12567 |
| 1.63017 |
| 0.64288 |
| 0.32595 |
| -0.7098 |
| -1.47915 |
| -1.52555 |
| 0.33471 |
| 1.08374 |
| 2.17015 |
| 2.87945 |
| 0.54828 |
| -0.16149 |
| 0.18203 |
| 0.14979 |
| 0.17335 |
| -0.17632 |

| |
|---|
| 1.03693 |
| 2.94101 |
| 3.70617 |
| 4.87921 |
| 1.24298 |
| 1.6407 |
| -0.87032 |
| 0.20012 |
| -0.48735 |
| -0.33185 |
| -0.50321 |
| 0.1091 |
| 0.19703 |
| 1.02912 |
| 1.89879 |
| 1.47218 |

| |
|---|
| 0.26018 |
| 1.08399 |
| 1.12285 |
| -0.19005 |
| -1.10041 |
| -1.06879 |
| -0.31775 |
| 0.19702 |

| |
|---|
| 0.18576 |
