## Supplementary Table 5 for "Pseudomonas aeruginosa uses kinases NahK and RetS to control the motile-biofilm switch"

| Locus Tag | Gene | *nosPnahK* vs wildtype ($log_2$ FC) *LB | *nosPnahK* vs wildtype ($log_2$ FC) *SMM |
|---|---|---|---|
| PA14_00180 | | 0.98381 | 1.76476 |
| PA14_00190 | fmt | 0.86849 | 1.3047 |
| PA14_00640 | phzH | 1.81782 | 2.34853 |
| PA14_00780 | | -1.25109 | -1.04605 |
| PA14_01120 | tsi6 | 6.6995 | 6.15006 |
| PA14_01130 | | 7.79994 | 8.08239 |
| PA14_01200 | | 2.31488 | 3.04583 |
| PA14_01220 | | 9.24986 | 9.73591 |
| PA14_01230 | | -1.66073 | -1.04285 |
| PA14_01240 | | -1.05883 | -1.04201 |
| PA14_01330 | | 1.08032 | 1.87592 |
| PA14_01460 | | -1.74204 | -0.99848 |
| PA14_01510 | | -1.44826 | -1.38039 |
| PA14_01770 | | 8.25268 | 8.55898 |
| PA14_02090 | | 0.95388 | 1.63896 |
| PA14_02100 | | 1.52257 | 1.77298 |
| PA14_02200 | | -1.57472 | -1.56798 |
| PA14_02220 | | -1.4101 | -1.36114 |
| PA14_02230 | cheW | -1.03656 | -1.4687 |
| PA14_02250 | cheA | -1.43646 | -1.27794 |
| PA14_02570 | mdcC | 1.25442 | -2.48223 |
| PA14_03163 | | 12.94657 | 12.24989 |
| PA14_03166 | | 10.52907 | 10.16491 |
| PA14_03180 | | 6.30204 | 4.87402 |
| PA14_03250 | | 10.82279 | 10.66423 |
| PA14_03265 | | 10.49247 | 10.68648 |
| PA14_03270 | | 12.40353 | 11.23572 |
| PA14_03285 | | 10.18195 | 10.10956 |
| PA14_03290 | | 1.92675 | 1.18005 |
| PA14_03300 | | 1.61705 | 1.25444 |
| PA14_03310 | | 8.6243 | 8.06514 |
| PA14_03320 | | 10.80248 | 10.36406 |
| PA14_03330 | | 9.31708 | 8.22007 |
| PA14_03340 | | 9.47942 | 8.82598 |
| PA14_03350 | | 13.15019 | 11.19666 |
| PA14_03360 | | 11.98839 | 10.63503 |
| PA14_03370 | | 11.26127 | 11.43391 |
| PA14_03380 | | 10.82593 | 10.84093 |
| PA14_03390 | | 5.24953 | 5.38753 |
| PA14_03470 | | 0.90815 | 0.87398 |
| PA14_04300 | | -0.85958 | -1.02899 |
| PA14_04710 | | 9.30092 | 7.21569 |
| PA14_04780 | laoA | 1.45465 | 1.76075 |
| PA14_05810 | amaB | 1.86485 | 3.29212 |
| PA14_05890 | | -3.12741 | -1.2919 |

| | | | |
|---|---|---|---|
| PA14_05970 | | 2.20177 | 1.19949 |
| PA14_06270 | | 1.66811 | 1.65747 |
| PA14_06530 | | -2.2165 | -1.15587 |
| PA14_07480 | | 13.89554 | 15.91811 |
| PA14_07550 | | 1.32092 | -2.37216 |
| PA14_07600 | | 1.41171 | 1.11696 |
| PA14_07630 | | -1.79023 | -1.59676 |
| PA14_07850 | | -1.09656 | -0.85433 |
| PA14_08830 | tufA | 0.91555 | 1.31769 |
| PA14_09280 | pchF | -2.12169 | -2.65518 |
| PA14_09400 | phzS | 2.27715 | 2.13639 |
| PA14_09420 | phzF1 | 2.72815 | 4.07002 |
| PA14_09440 | phzE1 | 2.93939 | 3.92497 |
| PA14_09450 | phzD1 | 2.82524 | 3.79163 |
| PA14_09470 | phzB1 | 3.59308 | 3.37544 |
| PA14_09490 | phzM | 1.84174 | 1.99368 |
| PA14_09520 | mexI | 2.53391 | 3.16249 |
| PA14_09880 | | 5.21658 | 3.05883 |
| PA14_10070 | | 6.49148 | 6.99172 |
| PA14_10090 | | 9.06748 | 10.24473 |
| PA14_10120 | | 7.76507 | 7.62819 |
| PA14_10340 | | 1.20432 | 3.44982 |
| PA14_10350 | | 1.08577 | 3.58907 |
| PA14_10370 | | 4.05764 | 6.88297 |
| PA14_10380 | | 3.39748 | 5.66805 |
| PA14_10560 | | -1.24986 | -0.85923 |
| PA14_10830 | | 9.71512 | 10.58036 |
| PA14_10840 | | 8.52511 | 9.19501 |
| PA14_10960 | | 6.72056 | 6.28661 |
| PA14_10970 | | 5.41938 | 5.20163 |
| PA14_10980 | | 7.5291 | 7.47583 |
| PA14_11140 | | 2.01939 | 1.66399 |
| PA14_11590 | | -3.53626 | -3.28772 |
| PA14_11970 | | -0.94612 | -0.74057 |
| PA14_12110 | | 1.35635 | 1.20992 |
| PA14_12900 | | 1.33947 | 1.52666 |
| PA14_13200 | | 8.43041 | 8.3798 |
| PA14_13210 | | 7.73632 | 10.18325 |
| PA14_13220 | | 6.17392 | 7.52169 |
| PA14_13340 | | 1.9183 | 2.16515 |
| PA14_13970 | | -2.1731 | -1.10723 |
| PA14_14100 | | -1.10005 | -0.81705 |
| PA14_14300 | | 7.0206 | 7.03037 |
| PA14_14310 | | 7.69164 | 7.88822 |
| PA14_14330 | | -2.21185 | -2.18333 |
| PA14_14420 | | 2.09329 | 2.6257 |
| PA14_14430 | | -2.046 | -0.89171 |
| PA14_14540 | | 7.18771 | 7.01572 |

| | | | |
|---|---|---|---|
| PA14_14550 | | 12.48048 | 12.01734 |
| PA14_14890 | hisS | 0.89472 | 1.41625 |
| PA14_14900 | | 1.11876 | 1.02345 |
| PA14_14930 | engA | 0.83609 | 0.73705 |
| PA14_15290 | | -2.59921 | -1.26323 |
| PA14_15350 | | 11.3346 | 11.38394 |
| PA14_15360 | | 11.29647 | 9.81032 |
| PA14_15380 | repA | 6.76047 | 7.55113 |
| PA14_15400 | | 8.62623 | 9.44019 |
| PA14_15430 | | 7.45597 | 7.4707 |
| PA14_15435 | | 9.13164 | 10.07629 |
| PA14_15450 | merD | 7.23383 | 7.99836 |
| PA14_15460 | merA | 10.8417 | 10.62176 |
| PA14_15470 | merP | 7.59216 | 6.49445 |
| PA14_15475 | merT | 6.74446 | 6.91278 |
| PA14_15480 | merR | 9.73294 | 10.90855 |
| PA14_15490 | | 5.83944 | 6.05163 |
| PA14_15500 | | 8.13876 | 8.64086 |
| PA14_15510 | traJ | 7.69636 | 8.44969 |
| PA14_15520 | trbJ | 7.10715 | 6.74791 |
| PA14_15540 | | 10.65757 | 9.02198 |
| PA14_15560 | | 10.09498 | 10.84732 |
| PA14_15570 | | 9.6678 | 10.35741 |
| PA14_15580 | | 11.66875 | 11.72762 |
| PA14_15590 | | 13.01793 | 11.36157 |
| PA14_15600 | | 10.42735 | 11.75782 |
| PA14_15610 | | 13.14728 | 12.66641 |
| PA14_15620 | | 5.51515 | 5.6815 |
| PA14_15630 | | 8.35938 | 9.3752 |
| PA14_15650 | | 8.32523 | 8.47661 |
| PA14_15660 | | 7.18251 | 8.20074 |
| PA14_16010 | | 2.29174 | -1.64948 |
| PA14_16150 | | -1.02001 | -0.9218 |
| PA14_16280 | | 1.40042 | 1.01458 |
| PA14_16290 | | 1.55215 | 2.21009 |
| PA14_16750 | | -1.4934 | -2.26872 |
| PA14_17310 | kdsA | 0.85034 | 1.1794 |
| PA14_18630 | | 1.82108 | 1.61592 |
| PA14_18800 | | 1.03213 | 1.68199 |
| PA14_18810 | | 1.52071 | 1.41913 |
| PA14_19140 | pheC | 1.13895 | 1.32463 |
| PA14_19920 | | -3.65095 | -1.84898 |
| PA14_20460 | | 2.15972 | 1.7835 |
| PA14_20510 | | 3.33277 | 3.23202 |
| PA14_20520 | | 5.51675 | 6.82835 |
| PA14_20530 | | 9.92716 | 9.49309 |
| PA14_20600 | | 9.03327 | 8.64404 |
| PA14_21120 | | 2.13607 | 3.80736 |

| | | | |
|---|---|---|---|
| PA14_21140 | | -2.48855 | -5.65939 |
| PA14_21150 | | -4.49288 | -7.31736 |
| PA14_21240 | | -0.94438 | -1.13089 |
| PA14_21280 | | -0.99354 | -0.87333 |
| PA14_21600 | | -2.85141 | -2.66422 |
| PA14_22075 | | 1.60445 | 2.96745 |
| PA14_22080 | | 10.80712 | 9.9421 |
| PA14_22090 | | 7.55531 | 8.1181 |
| PA14_22100 | | 6.37659 | 7.04395 |
| PA14_22110 | | 6.73074 | 6.7153 |
| PA14_22120 | | 8.19446 | 8.85923 |
| PA14_22140 | | 5.93778 | 5.15791 |
| PA14_22190 | | 8.63146 | 9.01587 |
| PA14_22210 | | 5.88134 | 5.00381 |
| PA14_22230 | | 5.48035 | 5.58003 |
| PA14_22240 | | 5.52918 | 5.81745 |
| PA14_22250 | | 5.63277 | 6.58038 |
| PA14_22270 | | 10.43357 | 10.28436 |
| PA14_22290 | vacJ | 1.45532 | 1.60308 |
| PA14_22310 | | 1.09587 | 1.12664 |
| PA14_22340 | | -1.87139 | -3.71121 |
| PA14_22350 | actP | -1.62907 | -4.93854 |
| PA14_22500 | | 6.98497 | 6.78584 |
| PA14_22520 | | 7.2477 | 7.08857 |
| PA14_22530 | | 5.9061 | 4.86976 |
| PA14_22540 | | 4.46883 | 5.75085 |
| PA14_22550 | | 8.72623 | 8.86153 |
| PA14_22980 | | 1.3366 | 1.23543 |
| PA14_23200 | | 1.24092 | 1.27265 |
| PA14_23220 | ubiG | 0.81637 | 1.39895 |
| PA14_23350 | orfA | 11.37969 | 9.34459 |
| PA14_23360 | wzz | 13.48783 | 12.58701 |
| PA14_23370 | orfK | 13.61684 | 9.14093 |
| PA14_23380 | orfH | 14.35999 | 10.60071 |
| PA14_23390 | orfE | 12.42962 | 12.17274 |
| PA14_23400 | | 12.8241 | 12.04697 |
| PA14_23410 | orfJ | 12.46056 | 11.86468 |
| PA14_23420 | zbdP | 11.93096 | 11.04796 |
| PA14_23430 | hepP | 12.81144 | 9.81223 |
| PA14_23440 | orfL | 12.32081 | 10.14426 |
| PA14_23450 | orfM | 11.90372 | 12.09887 |
| PA14_23460 | orfN | 11.16829 | 13.63715 |
| PA14_24050 | xcpV | 1.58232 | 1.48545 |
| PA14_24100 | xcpZ | 1.06883 | 1.68616 |
| PA14_24180 | | -0.96044 | -0.90176 |
| PA14_24360 | | 9.78616 | 10.38121 |
| PA14_24690 | | 1.29697 | 1.04703 |
| PA14_25370 | | 1.56772 | 1.72123 |

| | | | |
|---|---|---|---|
| PA14_25690 | fabF1 | 0.96882 | 0.90294 |
| PA14_26060 | | -1.81264 | -1.73848 |
| PA14_26280 | | -1.21013 | -1.05709 |
| PA14_27100 | lipA | -1.03656 | -1.79137 |
| PA14_27630 | | 8.55679 | 12.20759 |
| PA14_27640 | | 7.97419 | 10.99047 |
| PA14_27650 | | 7.40425 | 8.45783 |
| PA14_27660 | | 5.96929 | 8.15996 |
| PA14_27675 | | 8.25489 | 8.63713 |
| PA14_27700 | | 5.6217 | 5.59897 |
| PA14_28130 | | -2.08636 | -1.0015 |
| PA14_28140 | | -1.80753 | -1.54655 |
| PA14_28150 | | -2.88617 | -1.64859 |
| PA14_28240 | | 7.11567 | 6.29053 |
| PA14_28250 | | 12.04442 | 8.26223 |
| PA14_28360 | | 2.32927 | 2.29031 |
| PA14_28370 | | 1.72061 | 1.81684 |
| PA14_28490 | | -0.75326 | 1.27268 |
| PA14_28750 | | 5.05549 | 6.0569 |
| PA14_28760 | | -2.33544 | -2.6652 |
| PA14_28770 | | 11.90879 | 12.92369 |
| PA14_28780 | | 10.2328 | 10.22422 |
| PA14_28790 | | 6.64652 | 6.8393 |
| PA14_28800 | | 12.21054 | 10.85059 |
| PA14_28810 | | 11.84085 | 14.81279 |
| PA14_28820 | | 12.42346 | 10.9244 |
| PA14_28830 | | 13.59379 | 12.42569 |
| PA14_28840 | | 14.53501 | 12.60423 |
| PA14_28850 | | 6.22106 | 6.41457 |
| PA14_28870 | | 5.3193 | 6.62934 |
| PA14_29210 | | -1.37534 | -1.45828 |
| PA14_29330 | | 10.38555 | 13.15416 |
| PA14_29470 | | -1.54648 | -3.20941 |
| PA14_29660 | | -1.5629 | 1.99334 |
| PA14_30100 | | 1.52214 | 2.01332 |
| PA14_30430 | | 2.26519 | 2.6453 |
| PA14_30440 | | 2.3295 | 3.6553 |
| PA14_30690 | | 10.33846 | 11.73653 |
| PA14_30740 | | -2.0214 | -2.47563 |
| PA14_30850 | | 6.33714 | 7.41807 |
| PA14_30860 | | 6.03215 | 7.28841 |
| PA14_30900 | | 6.71972 | 7.49246 |
| PA14_30910 | | 7.13935 | 6.66699 |
| PA14_30940 | | 5.14812 | 5.88352 |
| PA14_30960 | | 5.21005 | 6.23595 |
| PA14_30970 | | 10.21409 | 10.54822 |
| PA14_30980 | | 9.64926 | 8.59579 |
| PA14_30990 | | 8.3664 | 7.92214 |

| | | | |
|---|---|---|---|
| PA14_31000 | | 8.38754 | 8.45523 |
| PA14_31010 | | 9.12278 | 9.36308 |
| PA14_31030 | | 8.90391 | 8.48971 |
| PA14_31040 | | 8.91608 | 8.98186 |
| PA14_31050 | | 11.66058 | 10.40136 |
| PA14_31060 | | 10.15809 | 10.33087 |
| PA14_31070 | | 6.11681 | 7.45038 |
| PA14_31150 | | 7.15612 | 7.89564 |
| PA14_31160 | | 7.46564 | 8.05422 |
| PA14_31170 | | 5.56348 | 6.44724 |
| PA14_31180 | | 7.37585 | 7.96654 |
| PA14_31190 | | 6.59917 | 7.99042 |
| PA14_31230 | | 5.83904 | 6.45612 |
| PA14_31240 | | 10.80148 | 12.50918 |
| PA14_31270 | | 8.27856 | 8.84573 |
| PA14_31280 | | 8.75201 | 8.82224 |
| PA14_31330 | | -3.8276 | -5.68238 |
| PA14_31390 | | -3.23684 | -2.22364 |
| PA14_31540 | | -1.42131 | -2.17948 |
| PA14_31720 | | -1.01923 | -1.24109 |
| PA14_31930 | | -2.0811 | -2.65456 |
| PA14_32270 | | 1.95849 | 1.72309 |
| PA14_32390 | mexF | 1.28868 | 1.54389 |
| PA14_32520 | | -2.07782 | -2.6958 |
| PA14_32530 | | -1.32462 | -1.76284 |
| PA14_32820 | | 6.44567 | 9.59426 |
| PA14_32830 | | 10.46693 | 5.8792 |
| PA14_32840 | | 5.38021 | 4.86482 |
| PA14_32850 | | 8.34091 | 7.68868 |
| PA14_32950 | | 1.03822 | 1.5381 |
| PA14_33060 | | 2.11353 | 5.61508 |
| PA14_33190 | | -1.86267 | -1.60795 |
| PA14_33300 | | 10.67588 | 10.9274 |
| PA14_33310 | | 11.52428 | 11.97621 |
| PA14_33320 | | 10.92657 | 9.82189 |
| PA14_33330 | | 12.7134 | 11.60899 |
| PA14_33340 | | 10.651 | 11.47616 |
| PA14_33350 | | 11.51102 | 10.0288 |
| PA14_33360 | | 5.0315 | 6.22363 |
| PA14_33920 | | -2.71031 | -4.01961 |
| PA14_33970 | | 10.00887 | 10.13391 |
| PA14_33980 | | 9.70308 | 8.57192 |
| PA14_34690 | | -1.88891 | -2.45162 |
| PA14_34880 | | 1.23188 | 2.42169 |
| PA14_35160 | | 1.50749 | 1.49978 |
| PA14_35590 | pslM | -3.05721 | -2.04017 |
| PA14_35620 | pslK | -2.99789 | -1.05048 |
| PA14_35630 | pslJ | -3.78781 | -1.8368 |

| | | | |
|---|---|---|---|
| PA14_35640 | pslI | -3.21825 | -2.07222 |
| PA14_35650 | pslH | -3.79525 | -2.06927 |
| PA14_35670 | pslG | -3.4603 | -1.30981 |
| PA14_35680 | pslF | -5.00294 | -4.5331 |
| PA14_35690 | pslE | -5.79228 | -5.05447 |
| PA14_35700 | | 8.06867 | 8.93681 |
| PA14_35710 | | 8.37273 | 11.87487 |
| PA14_35720 | | 10.16628 | 11.44845 |
| PA14_35730 | | 7.37055 | 8.46978 |
| PA14_35740 | | 11.63314 | 11.60448 |
| PA14_35750 | | 9.01696 | 9.68232 |
| PA14_35760 | | 12.43094 | 11.97671 |
| PA14_35770 | | 11.99893 | 10.78767 |
| PA14_35780 | | 11.42972 | 10.02719 |
| PA14_35790 | | 11.59059 | 11.32859 |
| PA14_35800 | | 12.94576 | 11.98822 |
| PA14_35810 | | 13.40293 | 11.31751 |
| PA14_35820 | tnpS | 13.03456 | 10.8148 |
| PA14_35830 | tnpT | 11.84016 | 11.82975 |
| PA14_35840 | | 11.71435 | 12.57581 |
| PA14_35850 | | 12.99149 | 11.79689 |
| PA14_35860 | | 7.65593 | 9.41046 |
| PA14_35880 | | 9.71066 | 11.51017 |
| PA14_35890 | | 8.8713 | 9.58514 |
| PA14_35900 | | 7.91613 | 9.12726 |
| PA14_35920 | | 6.67685 | 8.53789 |
| PA14_35940 | | 6.2258 | 8.52663 |
| PA14_35950 | | 5.24661 | 7.62617 |
| PA14_35970 | | 5.60246 | 7.40854 |
| PA14_36000 | prpR | 9.63146 | 9.69748 |
| PA14_36010 | | 7.79952 | 7.87689 |
| PA14_36020 | | -1.46049 | -1.39329 |
| PA14_36030 | | 5.0463 | 5.15932 |
| PA14_36260 | | 1.26444 | 1.56953 |
| PA14_36270 | | -1.19825 | -1.32596 |
| PA14_36345 | exoY | 1.52354 | 2.68358 |
| PA14_36400 | | -2.23982 | -1.27834 |
| PA14_37130 | | -1.19999 | -1.26144 |
| PA14_37140 | | -2.77419 | -2.20901 |
| PA14_37170 | | 7.57105 | 7.87883 |
| PA14_37250 | | -2.21591 | -4.57813 |
| PA14_37610 | | -0.82139 | -1.2541 |
| PA14_37670 | | 4.11353 | 3.09661 |
| PA14_37680 | | 2.39314 | 2.13175 |
| PA14_37770 | | 1.24073 | 1.11112 |
| PA14_37780 | | 1.28327 | 1.31887 |
| PA14_37870 | sppB | -2.34249 | -3.23045 |
| PA14_38140 | | -1.07611 | -1.80144 |

| | | | | |
|---|---|---|---|---|
| PA14_38320 | | | 1.61883 | 2.62193 |
| PA14_38610 | | | 1.01149 | -1.16011 |
| PA14_38840 | | | -6.64294 | -8.77921 |
| PA14_38970 | | | -7.69935 | -8.04113 |
| PA14_38990 | | | -5.21479 | -5.14587 |
| PA14_39100 | | | 3.50309 | 2.05624 |
| PA14_39470 | | | 12.28222 | 10.6538 |
| PA14_39480 | | | 13.24377 | 12.81443 |
| PA14_39880 | phzG2 | | 5.74754 | 7.12698 |
| PA14_39945 | phzC2 | | 3.01713 | 3.2077 |
| PA14_39960 | phzB2 | | 2.68444 | 3.70953 |
| PA14_39970 | phzA2 | | 3.6652 | 3.51983 |
| PA14_40110 | | | -1.70194 | -1.17821 |
| PA14_40470 | | | 9.25856 | 9.13383 |
| PA14_41560 | | | -1.67718 | 2.63358 |
| PA14_42050 | | | -2.33726 | -2.89891 |
| PA14_42060 | | | -2.39514 | -2.54417 |
| PA14_42140 | | | -1.12238 | 2.69193 |
| PA14_42150 | | | -1.10931 | 1.86095 |
| PA14_42620 | pscR | | -3.54959 | -4.11101 |
| PA14_43070 | hcp2 | | 2.97289 | 5.781 |
| PA14_43080 | vgrG14 | | 9.31777 | 7.89561 |
| PA14_43090 | tap | | 8.67862 | 7.01246 |
| PA14_43100 | rhsP2 | | 12.78321 | 9.15907 |
| PA14_43510 | | | 0.98189 | 1.11808 |
| PA14_43610 | | | -1.38481 | -2.36607 |
| PA14_43710 | | | 1.27249 | -1.76095 |
| PA14_43720 | | | -1.63925 | -2.69632 |
| PA14_44210 | | | 1.54408 | 2.19113 |
| PA14_44270 | | | 1.19544 | 1.48287 |
| PA14_44311 | | | 2.96683 | 0.899 |
| PA14_44480 | pemB | | -1.15287 | -1.67487 |
| PA14_44590 | | | -1.39567 | -1.52222 |
| PA14_44710 | xdhA | | -1.46135 | -0.80575 |
| PA14_44760 | | | -1.65147 | -1.1415 |
| PA14_45120 | | | 1.83716 | 3.86079 |
| PA14_46070 | gbuA | | -2.50319 | -3.19005 |
| PA14_46460 | | | 10.30216 | 9.96475 |
| PA14_46510 | | | 13.01702 | 10.35413 |
| PA14_46520 | | | 13.77302 | 13.34136 |
| PA14_46530 | | | 10.74582 | 10.7691 |
| PA14_46540 | | | 11.37707 | 13.58222 |
| PA14_46550 | | | 12.28496 | 12.96631 |
| PA14_46570 | | | 9.81566 | 9.53861 |
| PA14_46590 | | | 7.64889 | 7.85842 |
| PA14_46600 | | | 6.02915 | 5.38196 |
| PA14_46610 | | | 7.26434 | 7.73491 |
| PA14_46620 | | | 8.81191 | 9.04573 |

| | | | |
|---|---|---|---|
| PA14_46810 | | 0.93586 | 1.01274 |
| PA14_47030 | | -1.3879 | -2.00275 |
| PA14_47530 | | -1.54208 | -1.33107 |
| PA14_48040 | aprI | 1.77867 | 1.49177 |
| PA14_48060 | aprA | 1.9693 | 1.334 |
| PA14_48100 | aprE | 1.03283 | 1.0112 |
| PA14_48150 | qslA | -5.98958 | -6.90386 |
| PA14_48450 | | 8.54554 | 8.69833 |
| PA14_48460 | | 6.36688 | 6.53751 |
| PA14_48470 | | 6.22187 | 5.37381 |
| PA14_48490 | | 6.57295 | 5.14643 |
| PA14_48500 | | 7.60379 | 6.71471 |
| PA14_48850 | | -2.37536 | -1.41179 |
| PA14_48880 | | 9.09195 | 10.15669 |
| PA14_48930 | | 4.90145 | 6.20845 |
| PA14_48950 | | -1.47633 | -3.35421 |
| PA14_48970 | | -2.20423 | -4.71297 |
| PA14_48980 | | -2.47643 | -3.93161 |
| PA14_48990 | | -4.13951 | -3.35331 |
| PA14_49000 | | -5.98595 | -5.69395 |
| PA14_49010 | xisF5 | 5.58792 | 7.98662 |
| PA14_49020 | pf5r | 10.20301 | 10.61424 |
| PA14_49030 | | 13.01689 | 12.15702 |
| PA14_49300 | | 1.35835 | 4.09081 |
| PA14_49320 | | 1.02486 | 0.95457 |
| PA14_49480 | | 13.20184 | 10.80322 |
| PA14_49510 | pyoS3I | 10.84769 | 12.58621 |
| PA14_49520 | pyoS3A | 11.13198 | 12.37609 |
| PA14_49540 | | 8.96655 | 8.6506 |
| PA14_49570 | | -1.80733 | -1.90265 |
| PA14_49710 | | -2.35038 | -2.76959 |
| PA14_49740 | | 1.1387 | 1.36133 |
| PA14_49760 | rhlC | -0.96538 | -1.04858 |
| PA14_49920 | | 1.42654 | 2.41559 |
| PA14_49990 | | 0.84831 | 0.98371 |
| PA14_50290 | fliC | -0.7959 | -1.36156 |
| PA14_50610 | | -1.42248 | -1.14126 |
| PA14_51100 | | -1.93494 | -0.94181 |
| PA14_51120 | | -2.20352 | -1.45368 |
| PA14_51350 | phnB | 0.79964 | 2.31752 |
| PA14_51450 | cupC3 | 1.59473 | 1.37147 |
| PA14_51520 | spcU | 10.8079 | 9.47903 |
| PA14_51530 | exoU | 10.80538 | 11.29311 |
| PA14_51540 | | -2.42043 | -2.69086 |
| PA14_51550 | | 9.32482 | 9.99248 |
| PA14_51560 | | 10.29369 | 9.98094 |
| PA14_51570 | | 3.39452 | 2.99436 |
| PA14_51620 | | 7.03748 | 4.25835 |

| | | | | |
|---|---|---|---|---|
| PA14_51630 | | | 1.63304 | 6.80793 |
| PA14_51640 | | | 5.42764 | 6.16404 |
| PA14_51830 | | | -0.92423 | 2.48072 |
| PA14_52480 | | | 2.11472 | 2.41397 |
| PA14_52760 | | | 0.8405 | 1.25372 |
| PA14_52780 | aotQ | | 1.18642 | 1.06361 |
| PA14_52800 | acsA | | -1.09047 | -4.40603 |
| PA14_52810 | | | -1.23164 | -1.8775 |
| PA14_52900 | | | -1.63471 | -2.61627 |
| PA14_52930 | | | -1.35402 | -2.13662 |
| PA14_53190 | bolA | | 1.46608 | 1.25843 |
| PA14_53200 | | | 1.57568 | 1.14654 |
| PA14_53250 | cpbD | | 1.28608 | 3.29383 |
| PA14_53300 | | | 1.41574 | 3.41324 |
| PA14_53570 | | | 8.19676 | 9.02319 |
| PA14_53580 | | | 9.77104 | 10.09935 |
| PA14_53590 | | | 12.86679 | 10.88938 |
| PA14_53600 | | | 10.42953 | 9.39666 |
| PA14_53610 | | | 8.46945 | 6.86248 |
| PA14_53620 | | | -3.80991 | -3.08666 |
| PA14_53820 | | | 0.95378 | 2.46201 |
| PA14_53920 | | | -0.91055 | -1.40671 |
| PA14_54040 | | | -1.00094 | -1.27413 |
| PA14_54050 | | | 9.37143 | 9.47648 |
| PA14_54070 | | | 1.14308 | 1.1998 |
| PA14_54080 | | | -1.89046 | -0.93517 |
| PA14_54130 | | | -3.46283 | -3.48331 |
| PA14_54600 | | | -1.06382 | -1.68111 |
| PA14_54740 | | | -1.44624 | -1.01624 |
| PA14_54750 | | | 7.2137 | 9.07115 |
| PA14_54830 | | | 0.9937 | 3.87635 |
| PA14_54850 | | | 7.13334 | 7.52065 |
| PA14_54860 | | | 8.84235 | 11.08172 |
| PA14_54870 | | | 7.73612 | 11.156 |
| PA14_54880 | | | 9.1443 | 10.20819 |
| PA14_54890 | | | 10.24303 | 9.99306 |
| PA14_54900 | | | 8.79331 | 10.34835 |
| PA14_54910 | | | 8.43389 | 9.29687 |
| PA14_54920 | | | 10.84772 | 11.75818 |
| PA14_54930 | | | 9.95228 | 9.63731 |
| PA14_54940 | | | 10.20105 | 11.15257 |
| PA14_54950 | | | 8.52906 | 10.20685 |
| PA14_54960 | | | 7.80895 | 11.18782 |
| PA14_54970 | | | 8.09063 | 9.55695 |
| PA14_54980 | ybtQ | | 8.3394 | 12.37403 |
| PA14_55000 | | | 5.52638 | 10.20493 |
| PA14_55020 | | | 5.61163 | 10.88447 |
| PA14_55030 | | | 5.88098 | 9.4722 |

| | | | | |
|---|---|---|---|---|
| PA14_55050 | | | 10.80521 | 10.84394 |
| PA14_55060 | | | 1.66766 | 1.49526 |
| PA14_55070 | | | 6.91482 | 9.81884 |
| PA14_55080 | | | 12.08243 | 11.93003 |
| PA14_55090 | | | 7.78576 | 11.13089 |
| PA14_55400 | | | 3.29105 | 3.53416 |
| PA14_55410 | lapC | | 4.4447 | 6.62141 |
| PA14_55750 | | | -1.75748 | -3.98784 |
| PA14_55790 | | | -1.77287 | -1.19325 |
| PA14_56590 | | | -1.13726 | 2.00628 |
| PA14_56910 | | | -2.46109 | -1.22953 |
| PA14_56990 | | | 1.36499 | 2.73968 |
| PA14_57180 | | | 0.9949 | 2.02605 |
| PA14_57210 | argJ | | 0.83598 | 0.97229 |
| PA14_57980 | | | 1.59772 | 1.30942 |
| PA14_58390 | dppA3 | | 2.50392 | 1.78536 |
| PA14_58410 | opdP | | 3.5477 | 2.18652 |
| PA14_58630 | | | 1.32555 | 0.93236 |
| PA14_58690 | | | -1.00856 | -1.15247 |
| PA14_58720 | | | 11.84636 | 9.01865 |
| PA14_58730 | pilA | | 4.25267 | 4.71041 |
| PA14_58740 | | | 5.95607 | 6.49993 |
| PA14_58760 | pilC | | 13.32375 | 10.55957 |
| PA14_58880 | | | -1.31821 | -2.21469 |
| PA14_58920 | | | 7.19466 | 6.49441 |
| PA14_58930 | | | 5.94927 | 5.0206 |
| PA14_58960 | | | 6.36977 | 5.5973 |
| PA14_58970 | | | 6.12583 | 5.96686 |
| PA14_58990 | | | 7.87025 | 7.70674 |
| PA14_59000 | | | 7.11285 | 6.80609 |
| PA14_59020 | | | 6.79162 | 6.28477 |
| PA14_59050 | | | 7.62657 | 7.64545 |
| PA14_59060 | | | 5.2725 | 4.91305 |
| PA14_59070 | | | 9.89615 | 9.70298 |
| PA14_59090 | | | 7.38392 | 7.41122 |
| PA14_59100 | | | 7.55064 | 6.49355 |
| PA14_59110 | | | 5.1862 | 5.50632 |
| PA14_59120 | | | 5.74818 | 5.91697 |
| PA14_59130 | | | 8.123 | 6.15814 |
| PA14_59140 | | | 7.99586 | 6.90435 |
| PA14_59150 | | | 9.62371 | 8.82606 |
| PA14_59160 | crpP | | 5.36793 | 5.39793 |
| PA14_59170 | | | 10.0725 | 11.00578 |
| PA14_59180 | | | 8.77565 | 8.71336 |
| PA14_59190 | | | 11.92877 | 11.71992 |
| PA14_59200 | | | 8.80344 | 8.6235 |
| PA14_59210 | | | 7.53022 | 6.95885 |
| PA14_59220 | | | 1.01618 | 3.47227 |

| | | | |
|---|---|---|---|
| PA14_59250 | pilN2 | 6.21531 | 5.16406 |
| PA14_59270 | pilO2 | 6.26154 | 5.6486 |
| PA14_59290 | pilQ2 | 6.5357 | 5.68011 |
| PA14_59310 | pilR2 | 6.64937 | 6.14654 |
| PA14_59320 | pilS2 | 6.09251 | 5.57524 |
| PA14_59340 | pilT2 | 7.80804 | 7.99915 |
| PA14_59350 | pilV2 | 8.38006 | 8.98351 |
| PA14_59360 | pilM2 | 5.87654 | 5.58562 |
| PA14_59370 | | 9.20595 | 10.8364 |
| PA14_59380 | | 11.27246 | 9.17262 |
| PA14_59390 | | 11.0447 | 10.97372 |
| PA14_59400 | | 11.24059 | 12.31276 |
| PA14_59430 | | 8.53266 | 6.76002 |
| PA14_59440 | | 8.31583 | 8.4345 |
| PA14_59470 | | 10.09778 | 7.43152 |
| PA14_59480 | | 9.84636 | 7.70923 |
| PA14_59490 | | 8.50815 | 6.67537 |
| PA14_59500 | | 8.1113 | 5.66894 |
| PA14_59520 | | 7.93798 | 5.05508 |
| PA14_59530 | | 8.02143 | 5.76593 |
| PA14_59540 | | 7.96387 | 8.11204 |
| PA14_59550 | | 11.60501 | 11.1299 |
| PA14_59580 | | 10.55363 | 8.7278 |
| PA14_59590 | | 9.71345 | 10.47865 |
| PA14_59600 | | 6.70712 | 7.56611 |
| PA14_59610 | | 11.72455 | 12.51894 |
| PA14_59620 | | 10.55626 | 12.37033 |
| PA14_59630 | | 9.63541 | 10.29847 |
| PA14_59690 | | 7.83742 | 6.85592 |
| PA14_59710 | cupD1 | 7.98793 | 7.89371 |
| PA14_59750 | cupD4 | 6.1579 | 6.79985 |
| PA14_59760 | cupD5 | 5.78148 | 5.93111 |
| PA14_59770 | rcsB | 11.03661 | 10.25279 |
| PA14_59780 | rcsC | 12.52762 | 12.70017 |
| PA14_59790 | pvrR | 12.21471 | 11.93332 |
| PA14_59800 | pvrS | 10.72519 | 11.64549 |
| PA14_59820 | | 6.68565 | 6.93345 |
| PA14_59830 | | 7.20409 | 9.04776 |
| PA14_59840 | | 13.82793 | 12.27121 |
| PA14_59845 | | 12.54698 | 14.35753 |
| PA14_59900 | | 6.24819 | 5.70651 |
| PA14_59910 | | 8.24941 | 6.24941 |
| PA14_59920 | | 7.18685 | 7.17106 |
| PA14_59930 | | 5.64044 | 4.98388 |
| PA14_59940 | | 9.51098 | 9.03989 |
| PA14_59950 | | 5.76008 | 5.16751 |
| PA14_59960 | dsbA2 | -2.82388 | -4.62428 |
| PA14_59970 | | 5.30887 | 7.01227 |

| | | | |
|---|---|---|---|
| PA14_59990 | | 6.23734 | 5.97323 |
| PA14_60000 | | 8.75768 | 8.37736 |
| PA14_60020 | | 7.82116 | 8.22629 |
| PA14_60030 | | 10.3238 | 11.8008 |
| PA14_60040 | | 8.95866 | 9.92144 |
| PA14_60050 | | 8.7814 | 9.53927 |
| PA14_60060 | | 7.17261 | 7.42098 |
| PA14_60070 | | 10.32896 | 13.13534 |
| PA14_60080 | | 11.23223 | 11.91647 |
| PA14_60090 | | 9.93402 | 9.42741 |
| PA14_60100 | dtd | 10.59938 | 8.69942 |
| PA14_60110 | | 10.8199 | 12.43884 |
| PA14_60120 | dcd2 | 10.40979 | 12.37983 |
| PA14_60130 | | 10.75588 | 10.87955 |
| PA14_60140 | | 9.36501 | 9.36461 |
| PA14_60190 | clpB | -0.97076 | 1.17256 |
| PA14_60280 | fimU | 9.77711 | 13.41414 |
| PA14_60290 | pilW | 10.04598 | 12.42867 |
| PA14_60300 | pilX | 9.20486 | 8.33658 |
| PA14_60310 | pilY1 | 11.77405 | 13.74464 |
| PA14_60320 | pilE | 9.93762 | 11.6408 |
| PA14_60730 | | 3.74494 | 2.39919 |
| PA14_60750 | pra | 2.78973 | 3.16474 |
| PA14_60790 | | 1.16159 | 1.49753 |
| PA14_60900 | | 1.46234 | 2.23609 |
| PA14_61130 | | -1.22775 | -1.6347 |
| PA14_61200 | | 2.02173 | 4.01681 |
| PA14_61300 | | -0.82801 | -1.83643 |
| PA14_61870 | | -0.87522 | -1.20207 |
| PA14_61920 | | -1.19876 | -1.66871 |
| PA14_61940 | | -1.26901 | -1.47515 |
| PA14_62390 | | 2.39492 | 3.69029 |
| PA14_62690 | | 1.11684 | 1.91335 |
| PA14_62720 | rpsO | 0.89148 | 0.9825 |
| PA14_63110 | | 2.41317 | 0.74815 |
| PA14_63120 | | 2.92937 | 0.82683 |
| PA14_63160 | pmrB | 1.73569 | 0.95042 |
| PA14_63170 | | 1.17947 | 0.72123 |
| PA14_63200 | | -1.31909 | -1.50039 |
| PA14_63330 | | -2.77842 | -0.94057 |
| PA14_63380 | | -1.41474 | -2.72883 |
| PA14_63410 | | 2.15821 | 1.24162 |
| PA14_63450 | | 7.76257 | 7.47448 |
| PA14_64335 | ureD | 1.28537 | 3.1361 |
| PA14_64430 | | 9.25392 | 6.06048 |
| PA14_64530 | | 4.69322 | 2.67565 |
| PA14_64950 | pncA | -3.16109 | -4.37915 |
| PA14_64960 | pncB1 | -1.99051 | -2.80178 |

| | | | |
|---|---|---|---|
| PA14_64980 | nadE | -1.09889 | -1.52361 |
| PA14_65090 | | 1.1867 | 1.79595 |
| PA14_65250 | hisZ | 1.36363 | 1.94344 |
| PA14_65260 | | 1.73988 | 2.54121 |
| PA14_66100 | | 4.20721 | 4.44304 |
| PA14_66200 | | 1.30783 | 1.67768 |
| PA14_66210 | | 2.28064 | 1.53025 |
| PA14_66220 | waaP | 2.09678 | 1.30522 |
| PA14_66240 | waaC | 1.56359 | 1.22834 |
| PA14_66250 | waaF | 1.55876 | 1.65009 |
| PA14_66540 | | -1.92891 | -2.06378 |
| PA14_66830 | phaD | 0.85145 | 1.23274 |
| PA14_67210 | tli5b1 | 11.79405 | 12.38838 |
| PA14_67370 | | -1.90826 | -1.13494 |
| PA14_67540 | | 1.33116 | 2.87552 |
| PA14_67580 | thiI | 0.92458 | -1.30617 |
| PA14_67850 | | -1.2737 | -1.71517 |
| PA14_69320 | | 1.46782 | 1.65308 |
| PA14_69330 | | -1.04308 | -0.84235 |
| PA14_69390 | algQ | -1.02669 | -0.88167 |
| PA14_69510 | | 9.48964 | 10.22279 |
| PA14_69520 | | 12.08608 | 11.82878 |
| PA14_69540 | | 8.6477 | 5.93566 |
| PA14_69630 | rnk | 2.07997 | 1.08512 |
| PA14_71160 | | -3.04819 | -2.9205 |
| PA14_71880 | | -1.01944 | -1.00542 |
| PA14_71890 | | -0.94684 | -1.84268 |
| PA14_72220 | | -2.20884 | -1.06648 |
| PA14_72370 | | 1.03286 | 1.59575 |
| PA14_72760 | | 1.67339 | 2.76762 |
| PA14_72870 | | -1.16287 | -1.61418 |
| PA14_72880 | | -2.20896 | -2.14689 |
| PA14_72890 | | -1.15177 | -1.06657 |
| PA14_73350 | soj | 0.90061 | 1.25917 |
| PA14_73400 | trmE | 1.21094 | 1.95881 |
